## Supplementary material for "Disordered proteins interact with the chemical environment to tune their protective function during drying": File S1: Synergy Paper Equations.docx

$$\mathrm{TFE}_{\mathrm{conformation}}=\sum_{k=1}^{n} \mathrm{RASA}_{k}*\mathrm{TFE}_{k}$$

$${\Delta G}_{\mathrm{Dimerization}}=\mathrm{TFE}_{\mathrm{Monomers}}-\mathrm{TFE}_{\mathrm{Dimer}}$$

$${\Delta G}_{\mathrm{Folding}}=\sum_{k=1}^{n} \mathrm{TFE}_{k}-\sum_{k=1}^{n} \mathrm{RASA}_{k}*\mathrm{TFE}_{k}$$
