## Supplementary figures and images for "Disordered proteins interact with the chemical environment to tune their protective function during drying"

### CAHSD_Beatine_6_3.pdf

CAHS D + Betaine 6mg 3

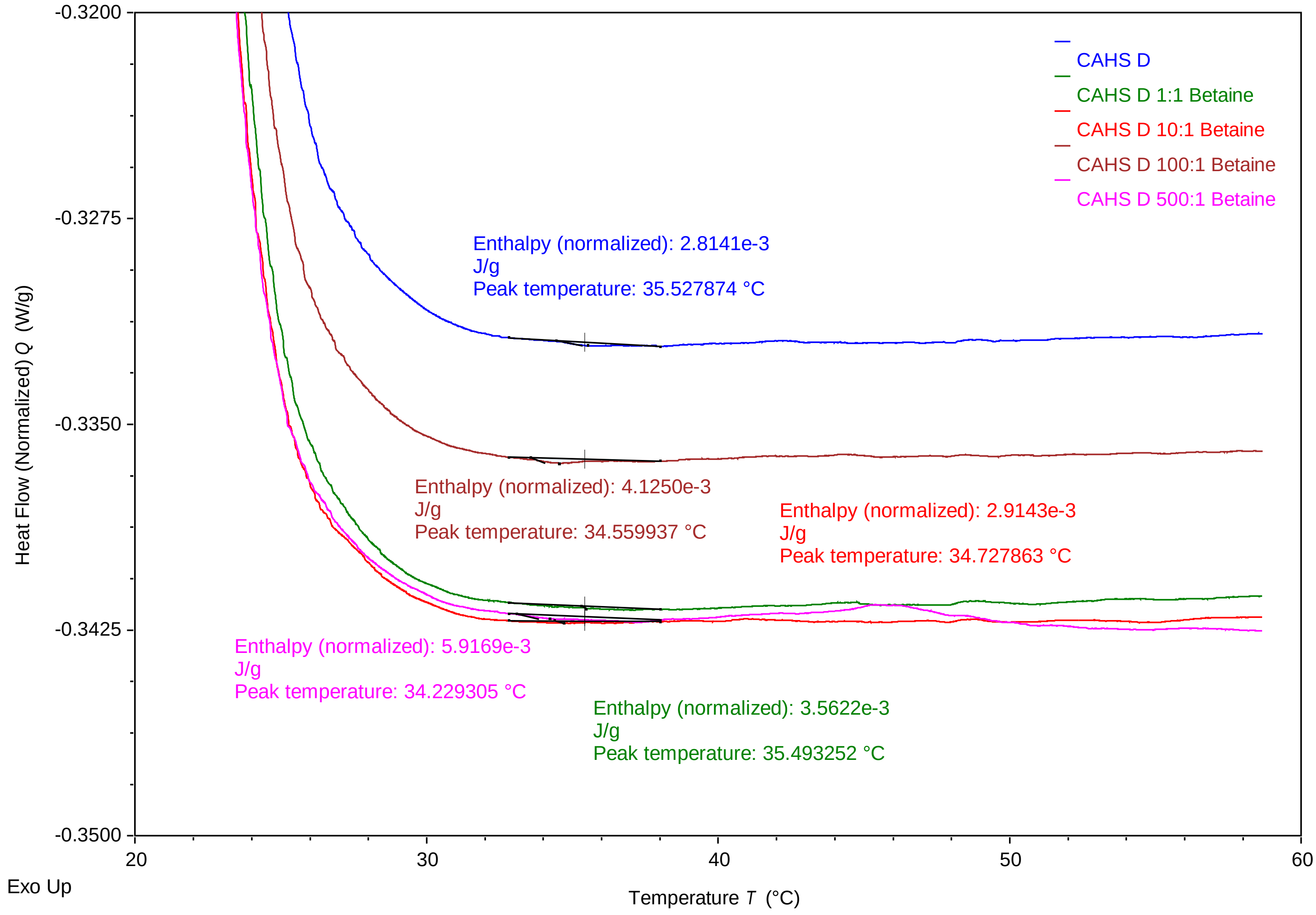


TA Instruments Trios V5.0.0.44616

### cahsd_betaine_6_1.pdf

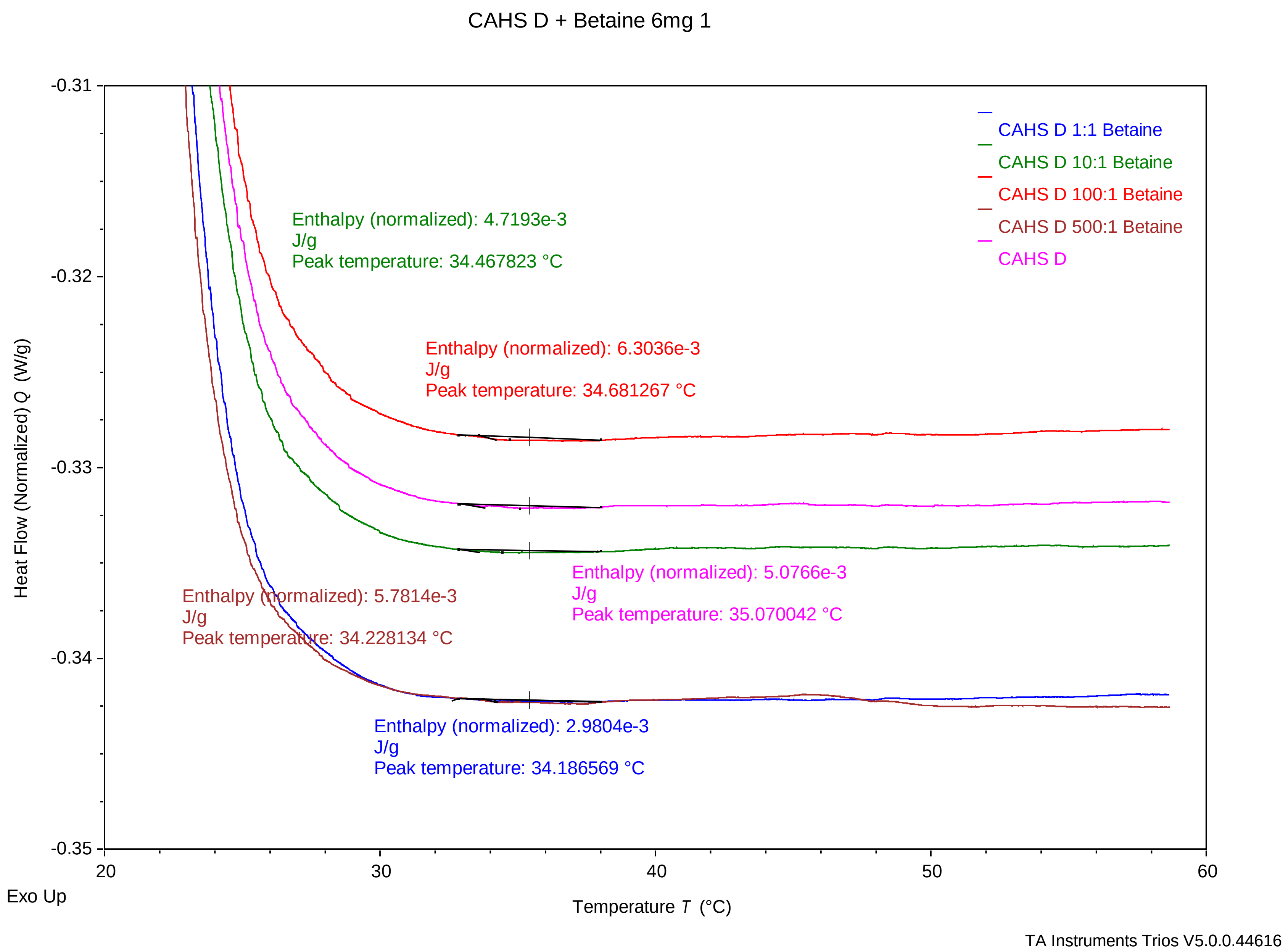

### CAHSD_Betaine_6_2.pdf

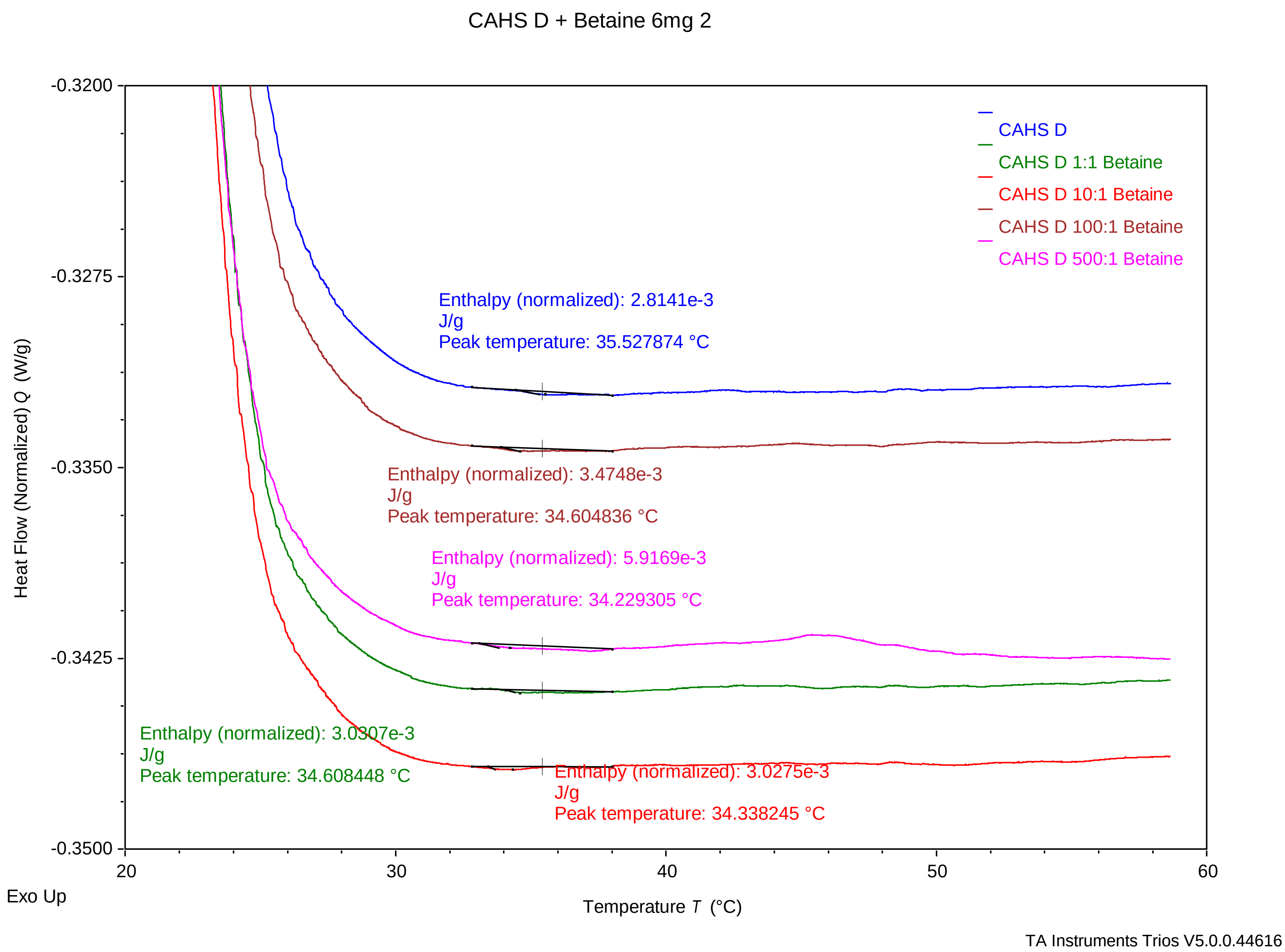

### CAHSD_Betaine_12_1.pdf

#### 12mg CAHS D + Betaine 1

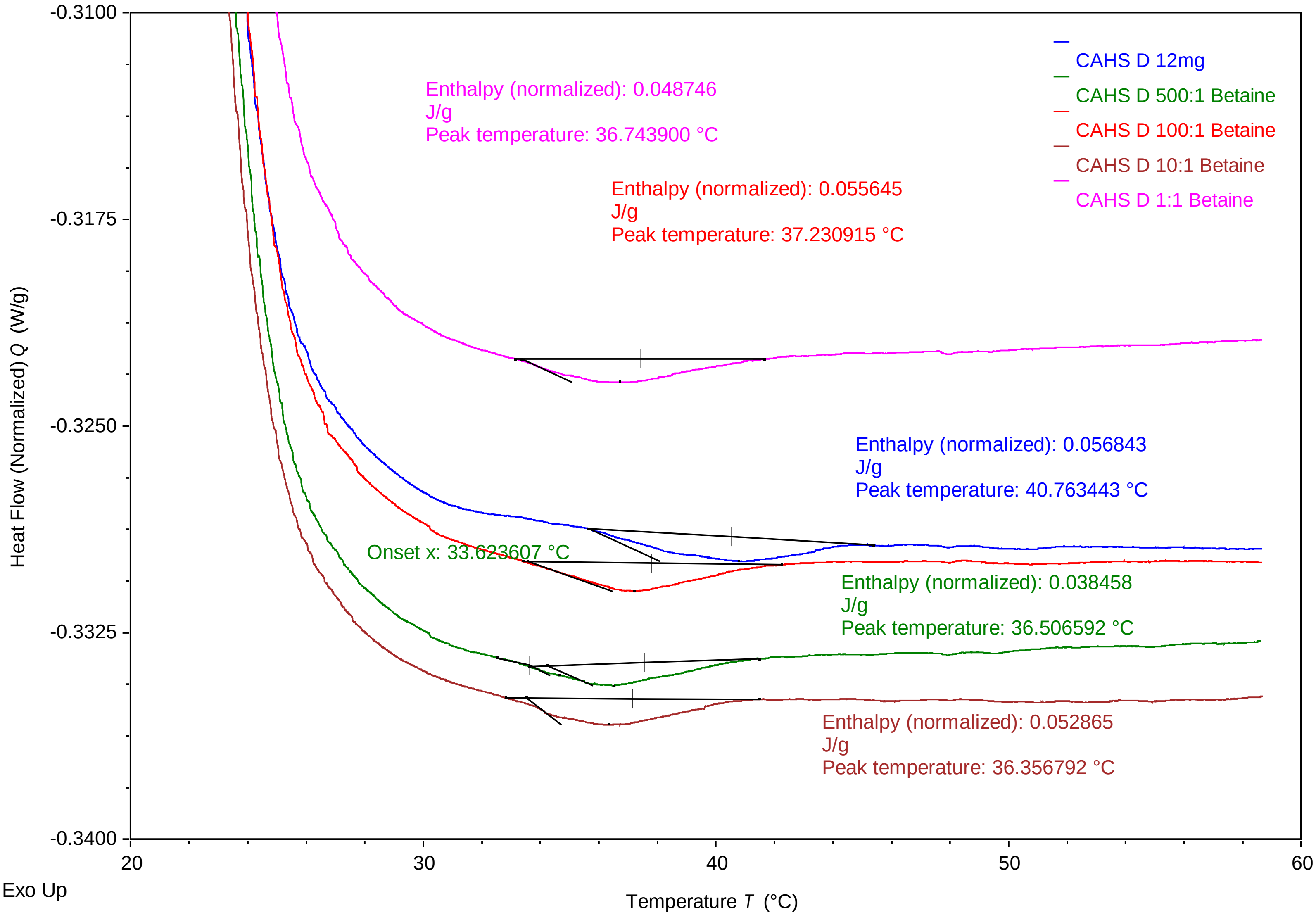


TA Instruments Trios V5.0.0.44616

### CAHSD_Betaine_12_2.pdf

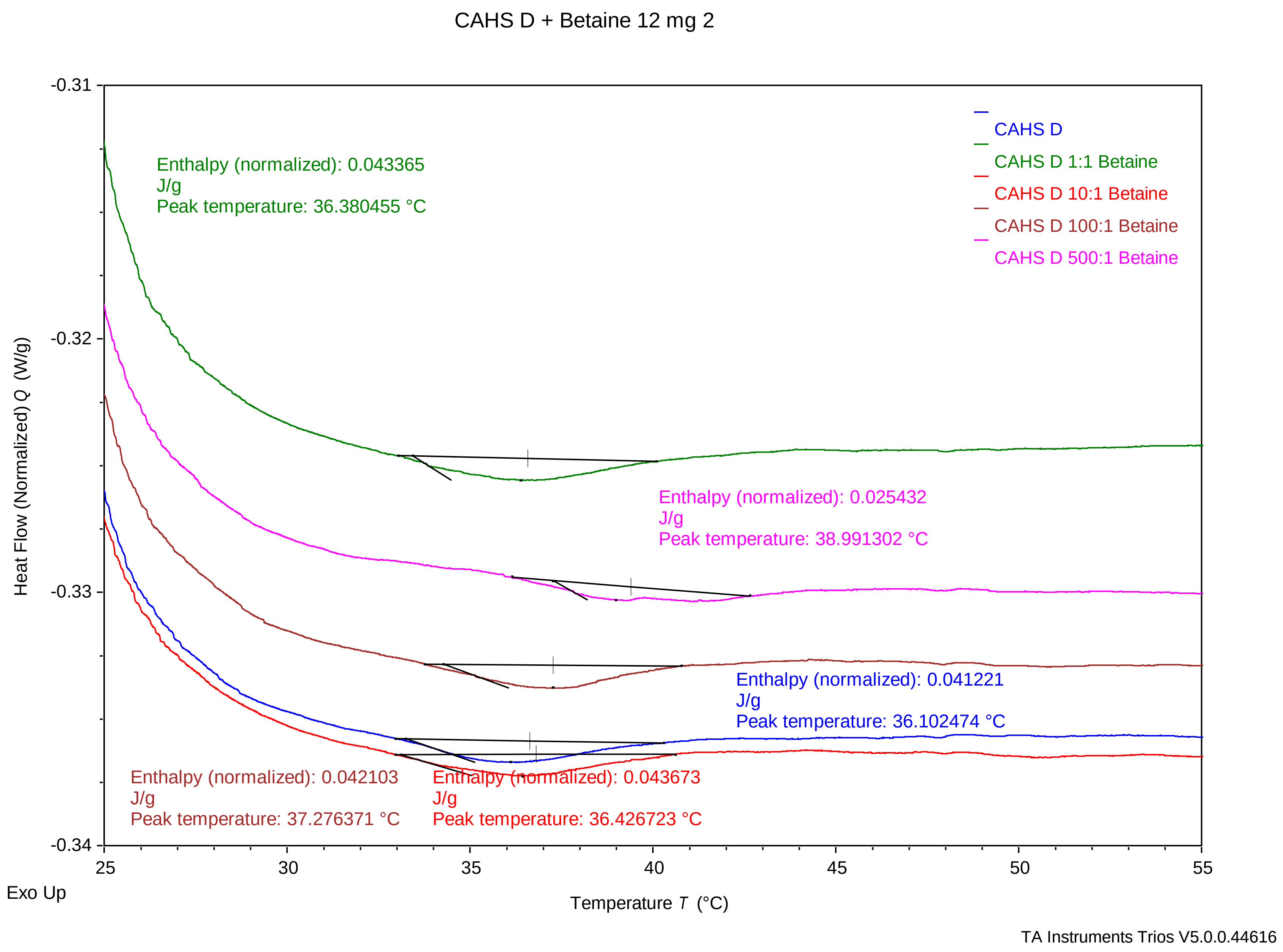

### CAHSD_Betaine_12_3.pdf

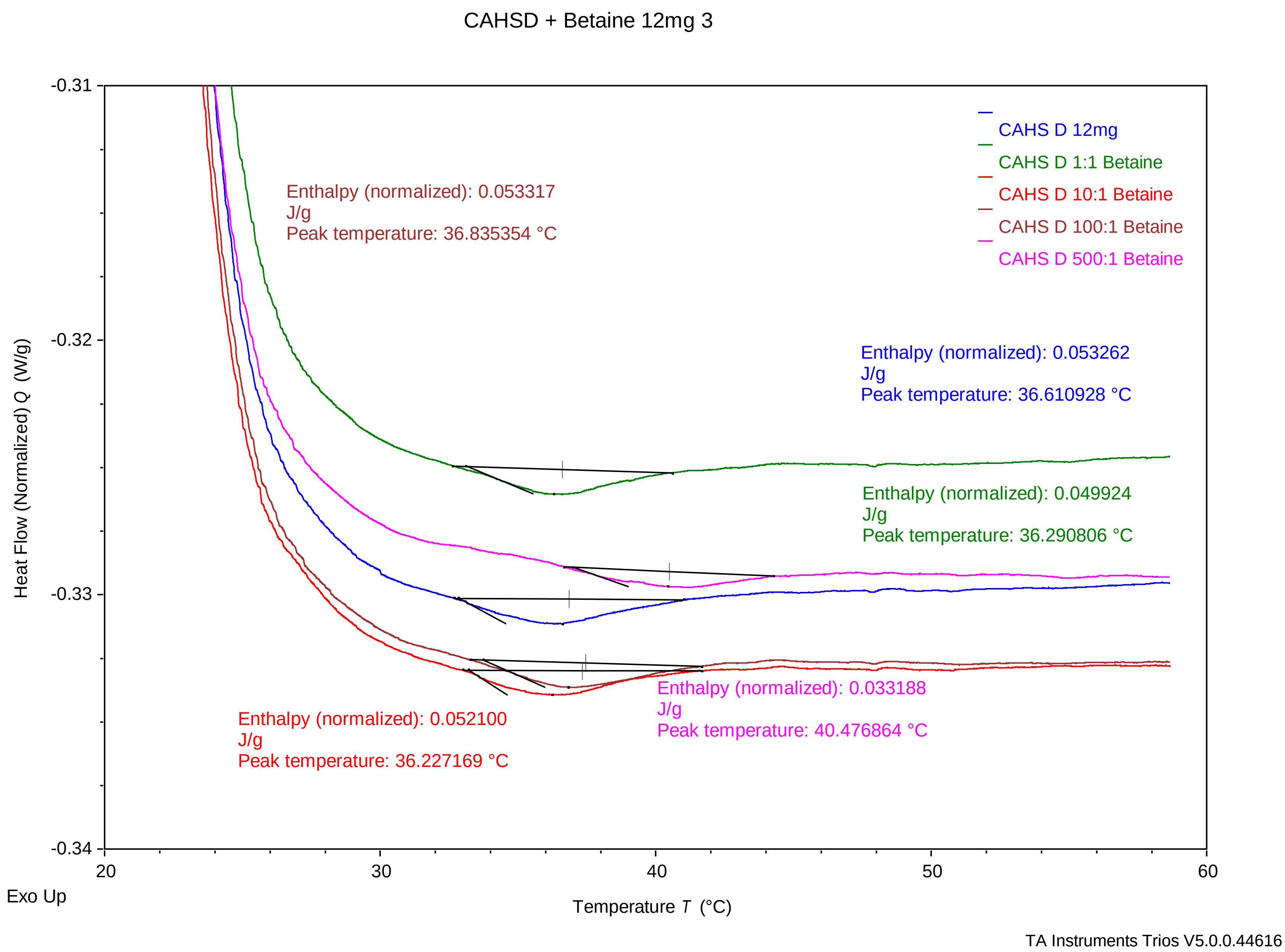

### CAHSD_Sucrose_6_1 .pdf

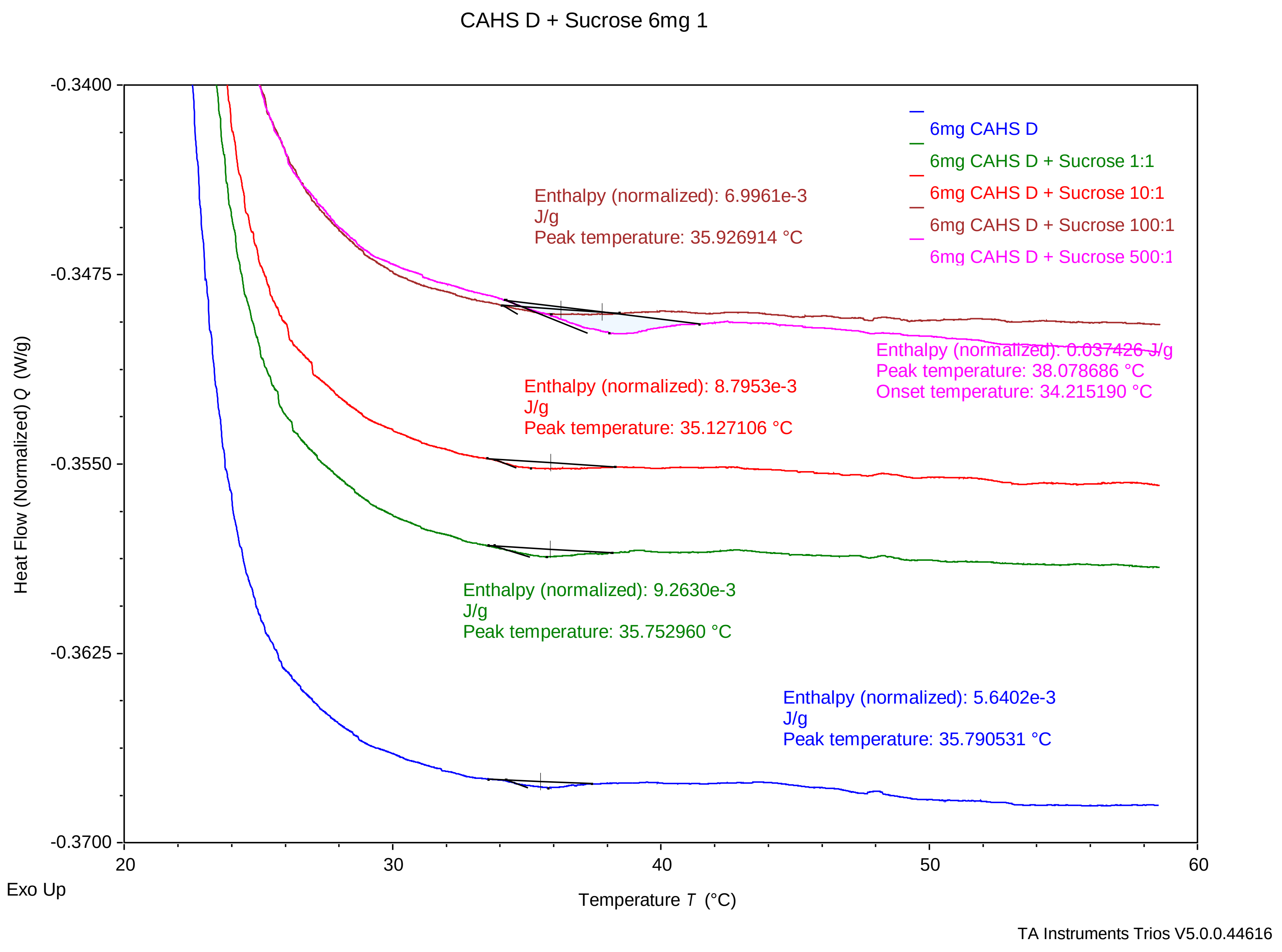

### CAHSD_Sucrose_6_2 .pdf

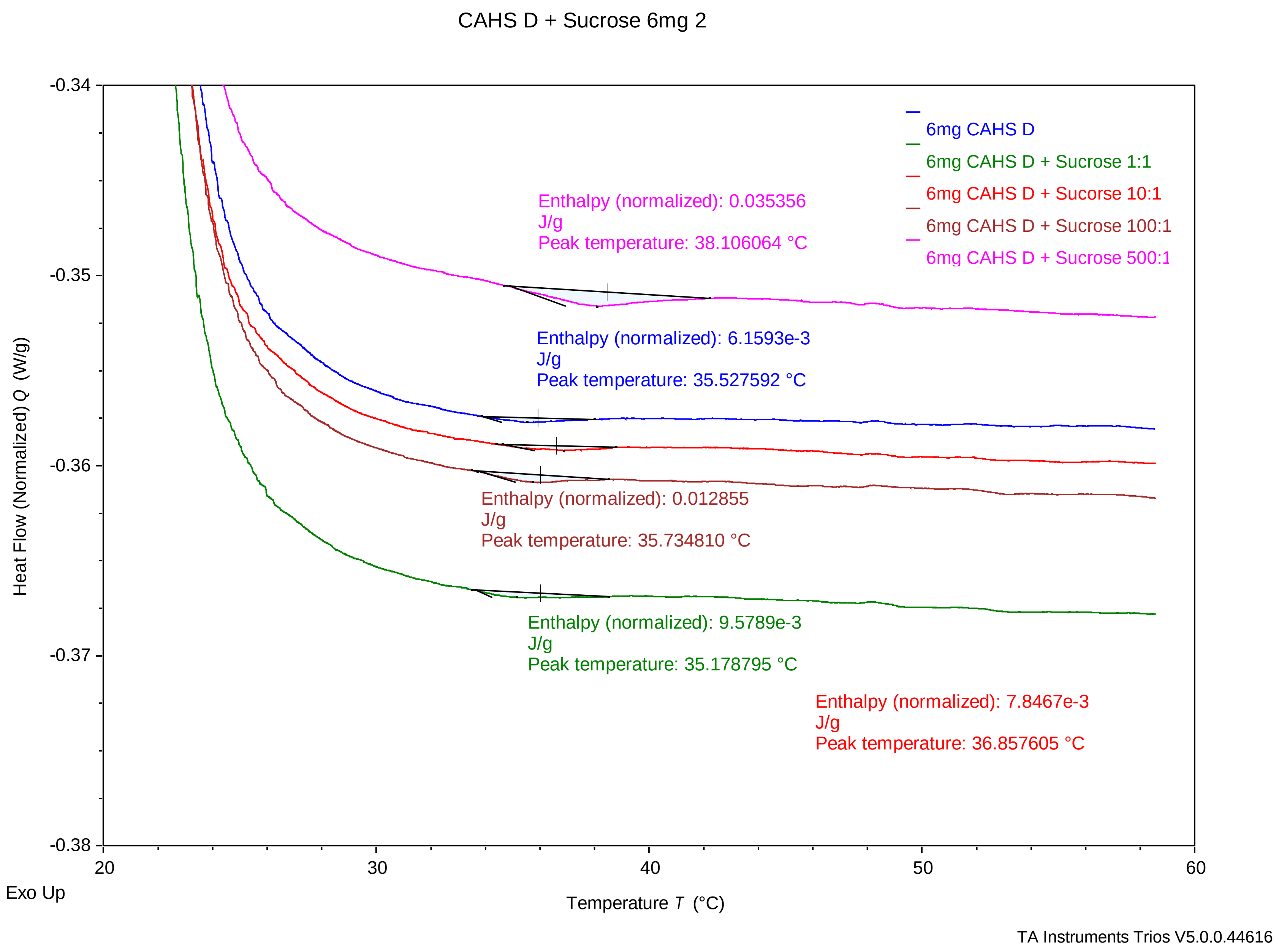

### CAHSD_Sucrose_6_3 .pdf

CAHS D + Sucrose 6mg 3

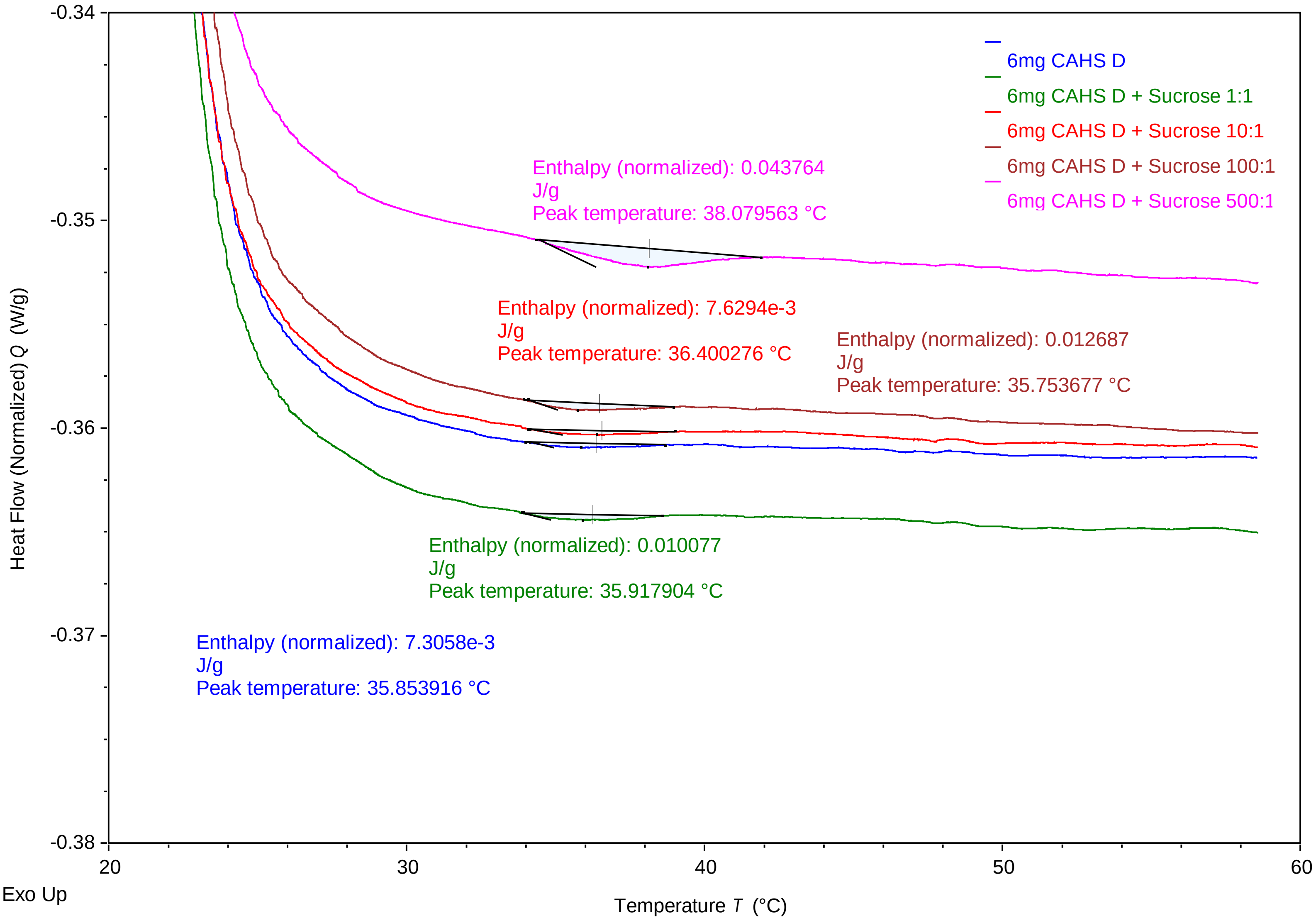


TA Instruments Trios V5.0.0.44616

### CAHSD_Sucrose_12_1.pdf

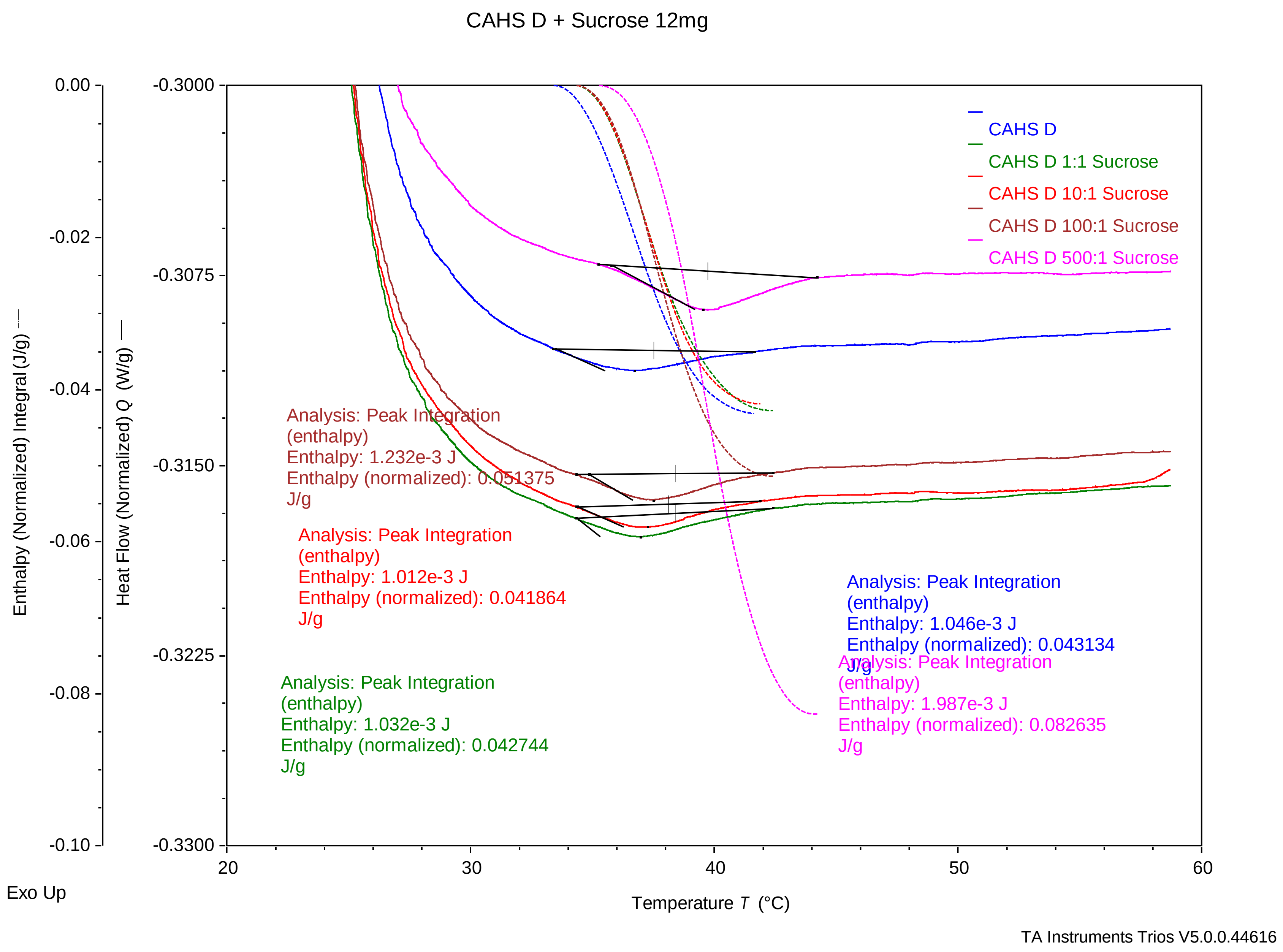

### CAHSD_Sucrose_12_2.pdf

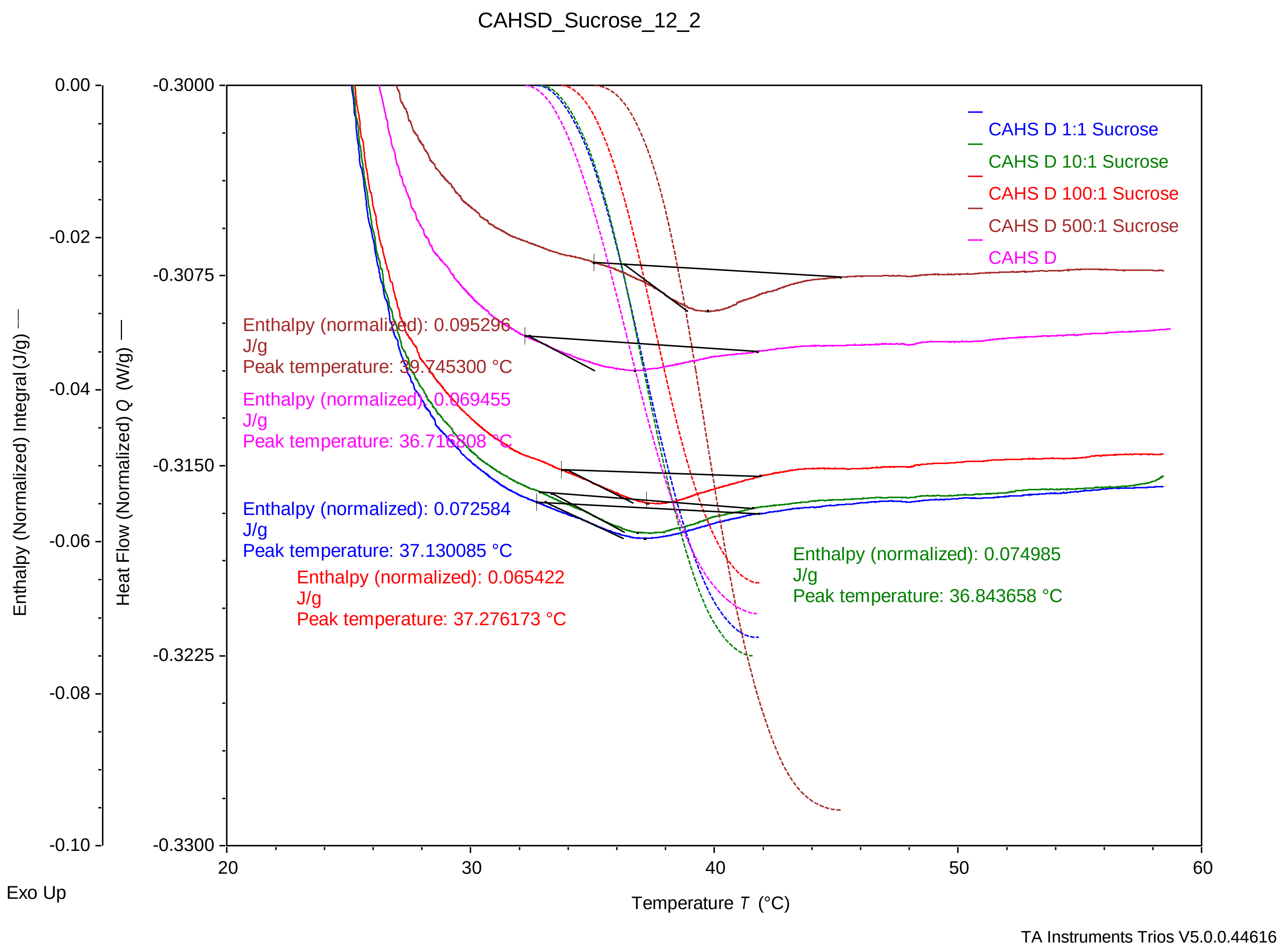

### CAHSD_Trehalose_6_1.pdf

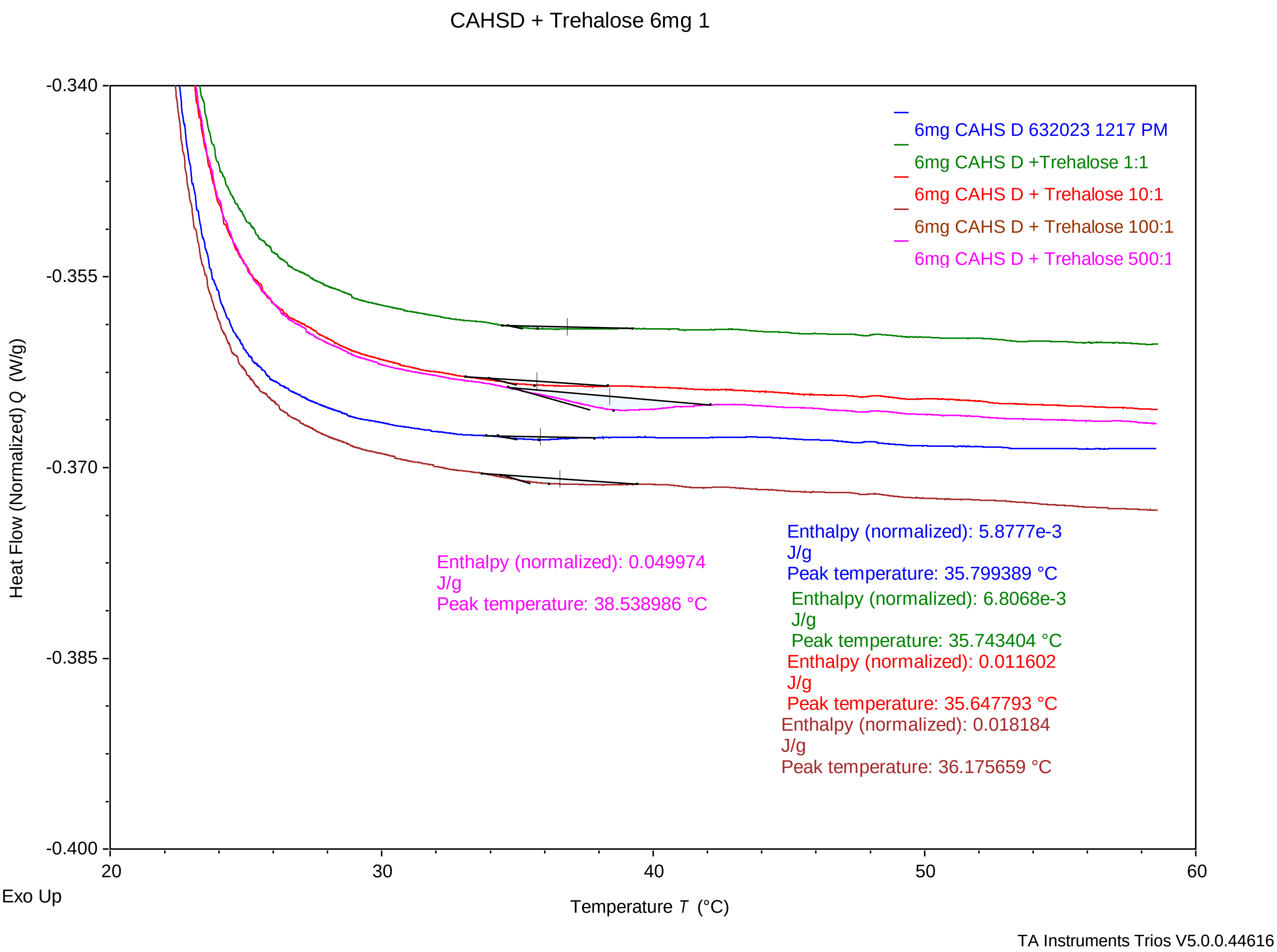

### CAHSD_Trehalose_6_2 .pdf

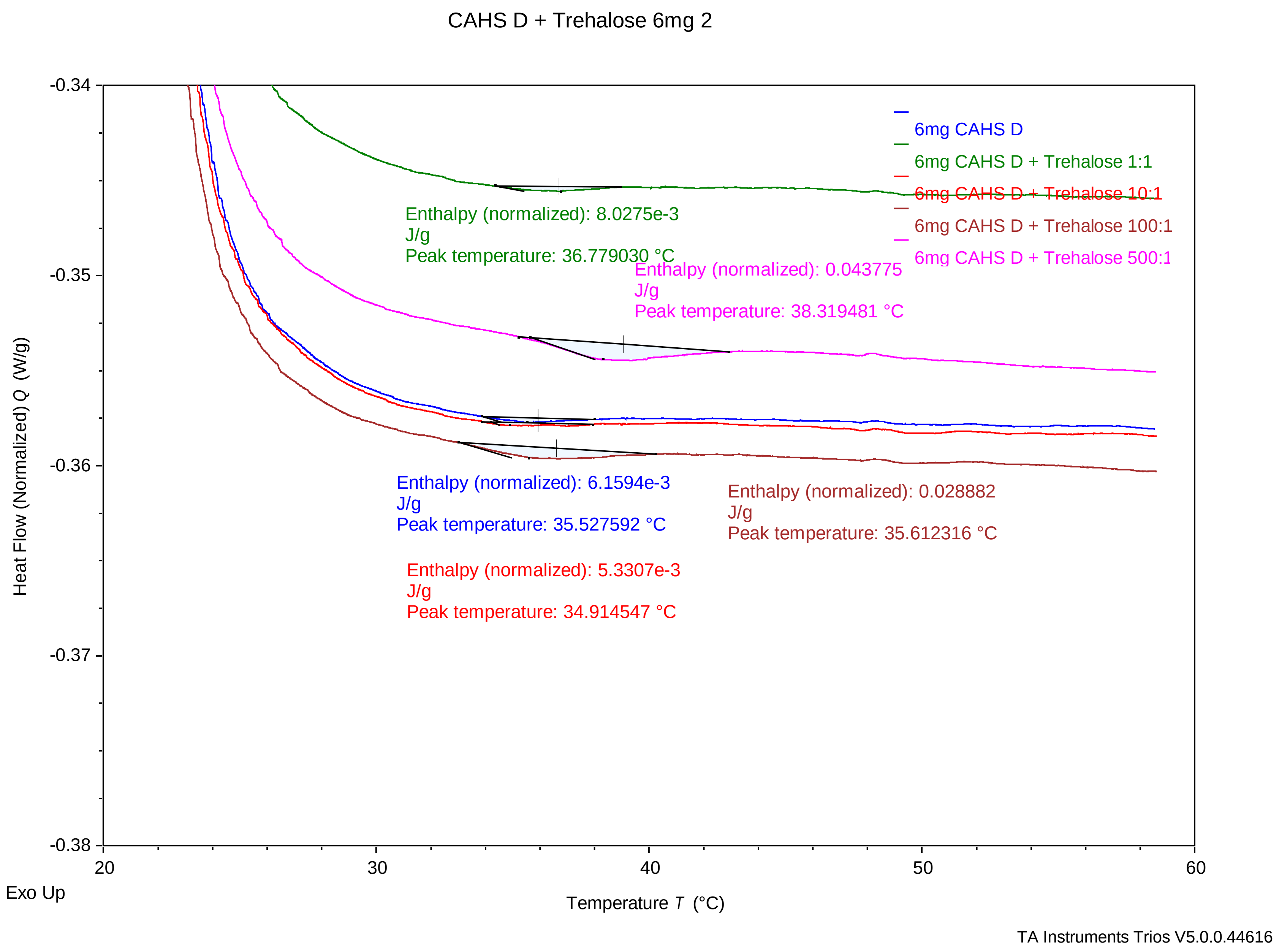

### CAHSD_Trehalose_6_3 .pdf

CAHS D + Trehalose 6mg 3

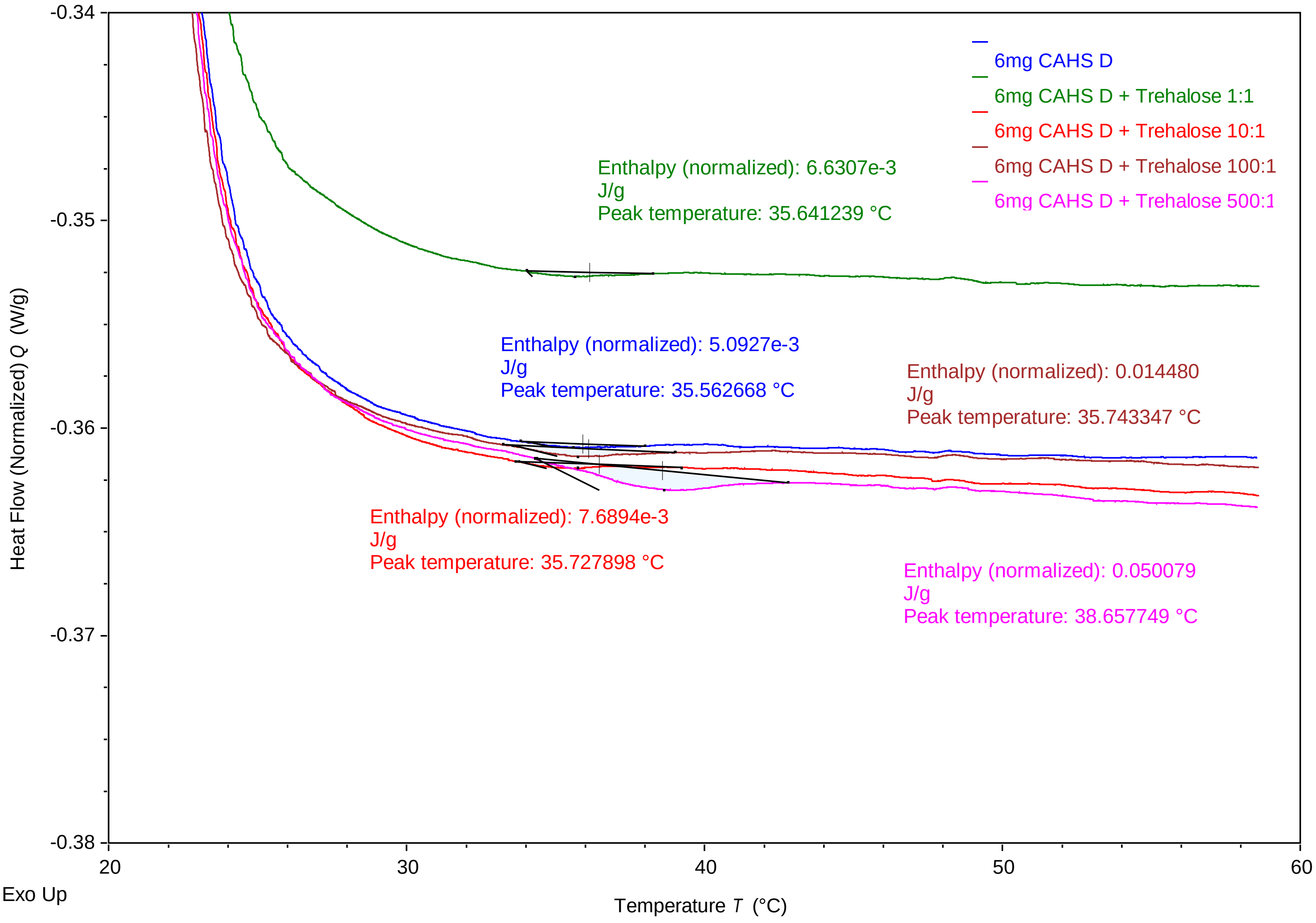


TA Instruments Trios V5.0.0.44616

### CAHSD_Trehalose_12_1 .pdf

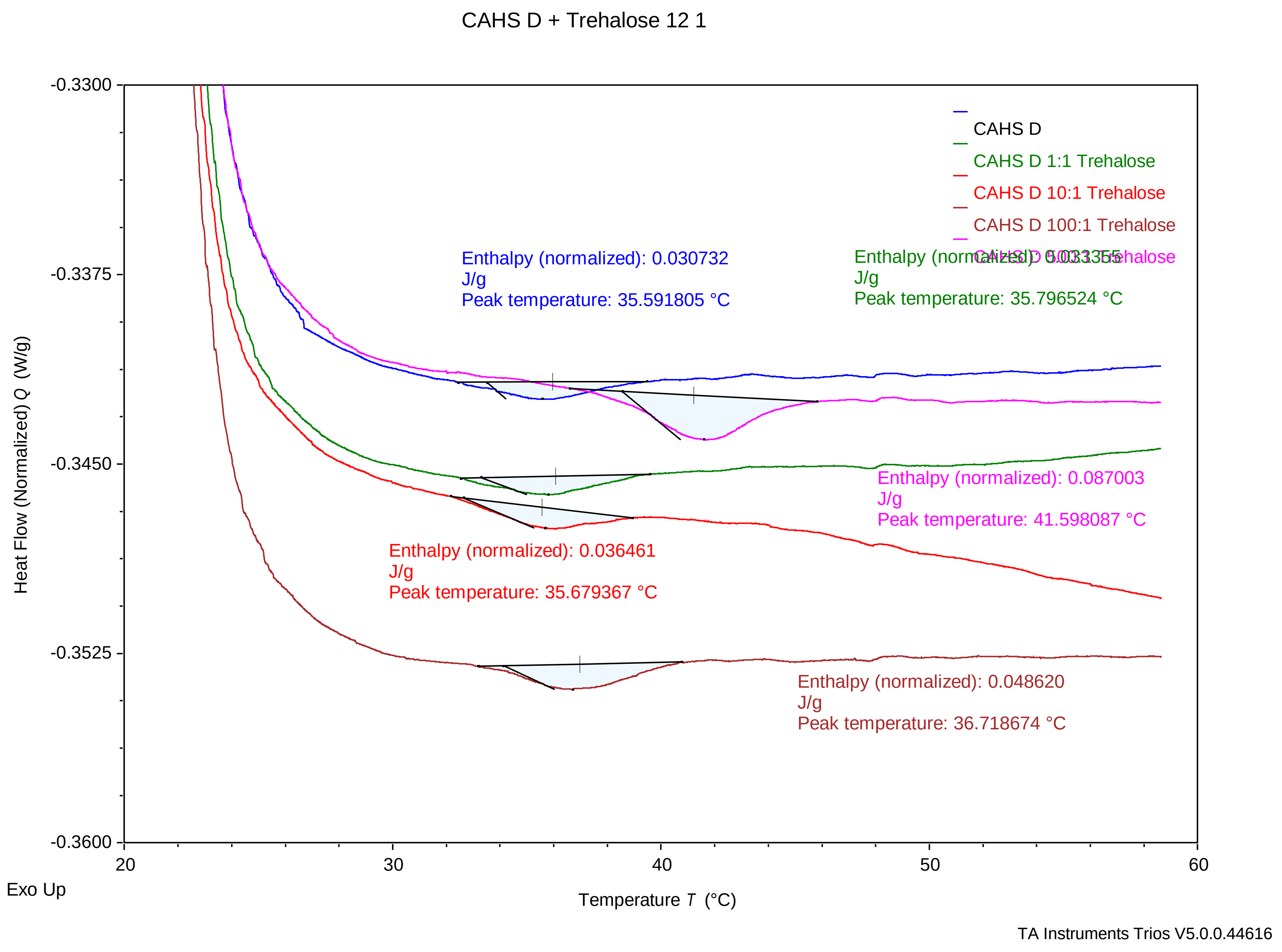

### CAHSD_Trehalose_12_2 .pdf

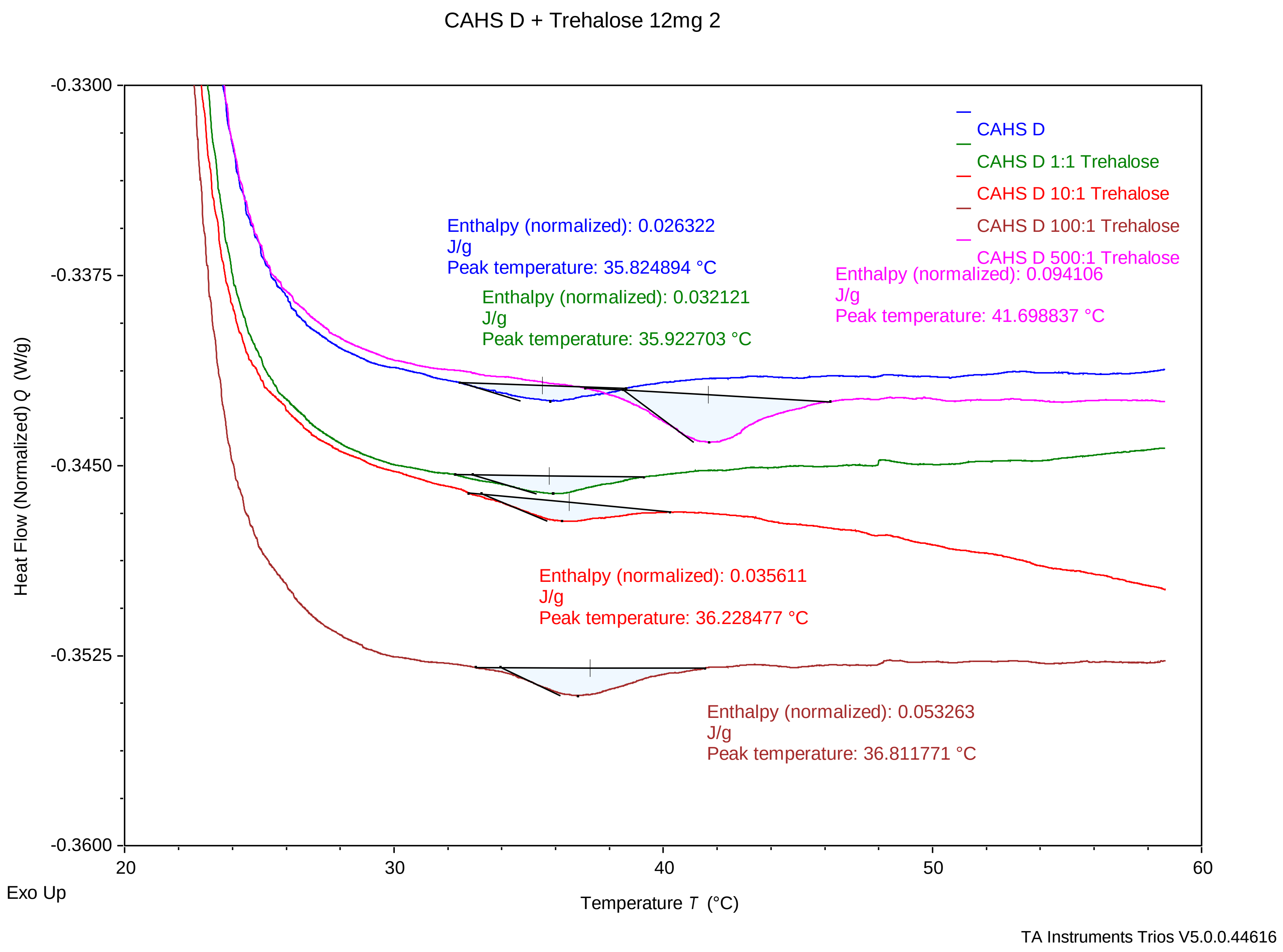

### CAHSD_Trehalose_12_3.pdf

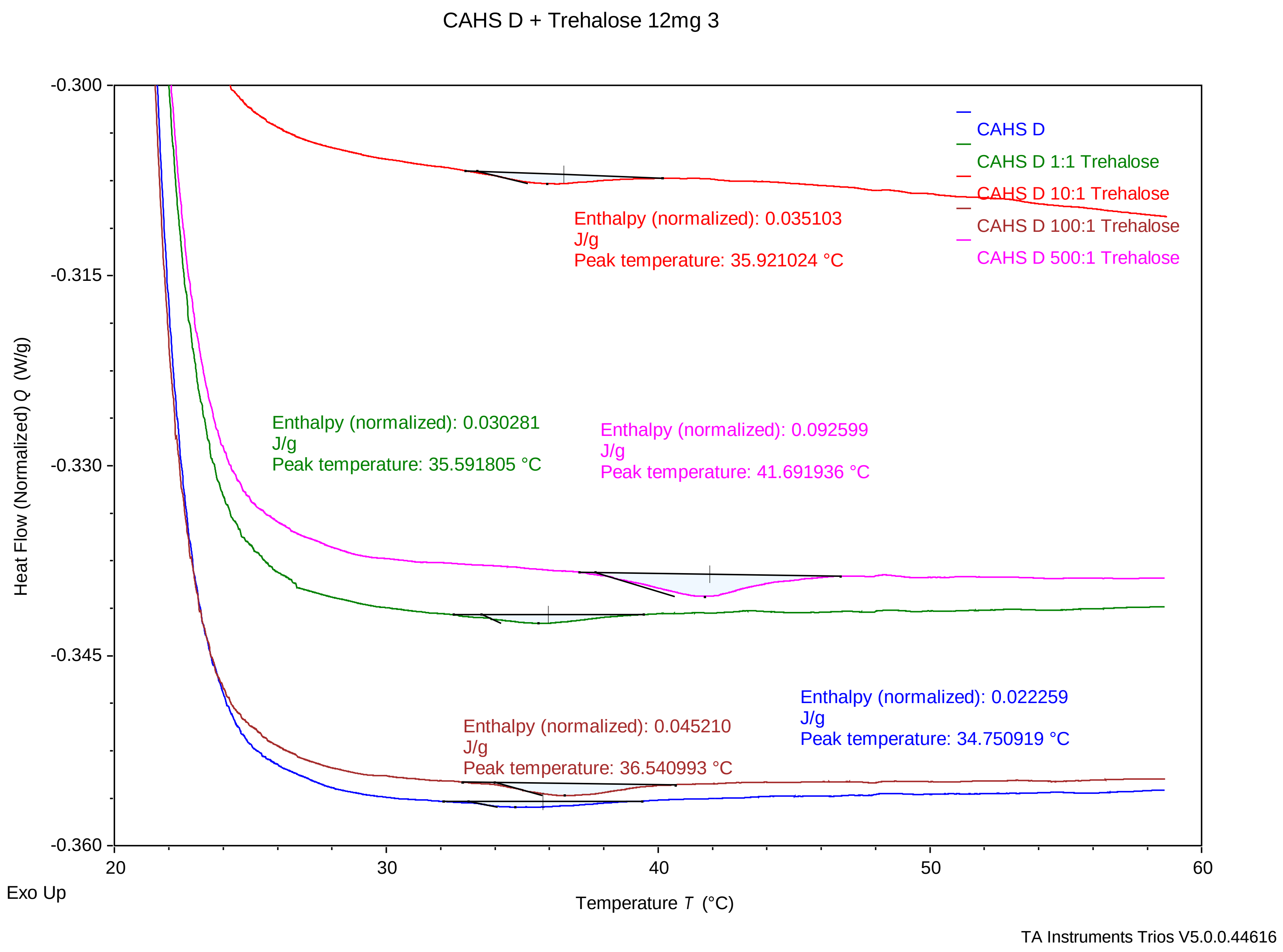
