## Supplementary material for "Disordered proteins interact with the chemical environment to tune their protective function during drying": File S3: 1C.R

```
#Set working directory----
setwd('~/Desktop/')

#Import necessary packages----
library(ggplot2)
library(dplyr)
library(readxl)
library(ggsignif)

#Import your dataset----
data <- read_xlsx('Example.xlsx')
#Change the factor order
data$Ratio <- factor(data$Sample,levels = c('Protein1',
                                            'Protein2',
                                            'Protein3'
))

#####make plot----
#create ggplot Environment----
ggplot(data = data, aes(x = Sample, y = Protect_add, fill = interaction(Sample, Calculation)))+
  #Make stacked bar graph---- 
##Make bottom layer first 
geom_col(data = data %>% filter(Protectant == 'Sugar'| Protectant == 'Mixed'), 
         position = position_dodge(width = .9), 
         color = 'black')+
  ##Make top layer w/ alpha adjustment
  geom_col(data = data %>% filter(Protectant == 'Protein'| Protectant == 'Mixed'), 
           position = position_dodge(width = .9), 
           alpha = .6, 
           color = 'black')+
  #Set y-axis scale
  scale_y_continuous(name = 'LDH Protection (%)',
                     breaks = seq(0,100,20),
                     limits = c(0,100))+  
  #Set graph labels
  labs(x = '',
       y = '%Protection',
       title = '')+
  #Make Black and white theme
  theme_bw()+
  #Adjust graph settings
  theme(
    plot.title = element_text(hjust = 0.05, size = 12, vjust = -8),
    axis.title = element_text(size =12),
    axis.text = element_text(size =10),
    legend.text=element_text(size=14), 
    legend.position = 'none', aspect.ratio = .5,
    panel.border = element_rect(colour = "black", fill=NA, size=1))+
  #add error bars based on standard deviation
  geom_errorbar(aes(ymin = Protect_add - Stdev, ymax = Protect_add + Stdev), width = .2, position = position_dodge(.9))+
  #custom assignment of bar color----
scale_fill_manual(values = c('honeydew', 'honeydew', 'honeydew',
                             'aquamarine4',   'aquamarine4', 'aquamarine4'))+
  #add significane values
  geom_signif(
    y_position = c(48,88,48),
    xmin = c(.75, 1.75, 2.75),
    xmax = c(1.25, 2.25, 3.25),
    annotation = c("NS", "***", "***"),
    tip_length = 0, color = "black")+
  #rename x-labels
  scale_x_discrete(breaks=c("Protein1", "Protein2","Protein3"),
                   labels=c("Protein1", "Protein2","Protein3"))
```
