## Supplementary material for "Disordered proteins interact with the chemical environment to tune their protective function during drying": File S3: 1D.R

```
#Set working directory----
setwd('~/Desktop/')

#Import necessary packages----
library(ggplot2)
library(dplyr)
library(readxl)
library(ggsignif)

#Import your dataset----
data <- read_xlsx('Figure_1_LEA_Synergy.xlsx', sheet = 'Peptide_Trehalose')
#Change the factor order
data$Sample <- factor(data$Sample,levels = c(
                                             'At11',
                                             'At22',
                                             'At44',
                                             'Aav11',
                                             'He11',
                                             'Av11',
                                             'At20'))


#####make plot----
#create ggplot Environment----
ggplot(data = data, aes(x = Sample, y = Protect_add, fill = interaction(Sample, Calculation)))+
  #Make stacked bar graph---- 
##Make bottom layer first 
geom_col(data = data %>% filter(Protectant == 'Sugar'| Protectant == 'Mixed'), 
         position = position_dodge(width = .9), 
         color = 'black')+
  ##Make top layer w/ alpha adjustment
  geom_col(data = data %>% filter(Protectant == 'Protein'| Protectant == 'Mixed'), 
           position = position_dodge(width = .9), 
           alpha = .6, 
           color = 'black')+
  #Set y-axis scale
  scale_y_continuous(name = ' LDH Protection (%)',
                     breaks = seq(0,100,20),
                     limits = c(0,90))+  
  #Set graph labels
  labs(x = 'LEA Motif',
       y = 'LDH Protection (%)',
       title = 'Trehalose: Peptide')+
  #Make Black and white theme
  theme_bw()+
  #Adjust graph settings
  theme(
    plot.title = element_text(hjust = 0.05, size = 12, vjust = -8),
    axis.title = element_text(size =12),
    axis.text = element_text(size =10),
    legend.text=element_text(size=14), 
    legend.position = 'none', aspect.ratio = .5,
    panel.border = element_rect(colour = "black", fill=NA, size=1))+
  #add error bars based on standard deviation
  geom_errorbar(aes(ymin = Protect_add - Stdev, ymax = Protect_add + Stdev), width = .2, position = position_dodge(.9))+
  #custom assignment of bar color----
scale_fill_manual(values = c( 'skyblue1','dodgerblue1', 'turquoise1', 'seagreen1', 'mediumpurple1','yellow1','coral1',
                                'skyblue3', 'dodgerblue3', 'turquoise3', 'seagreen3', 'mediumpurple3', 'yellow3','coral3'))+
  
  #add significane values
geom_signif(
  y_position = c(55, 55, 28, 28,73,52, 82),
  xmin = c(.75, 1.75, 2.75, 3.75,4.75,5.75, 6.75),
  xmax = c(1.25, 2.25, 3.25, 4.25, 5.25, 6.25, 7.25),
  annotation = c("NS", "**", "NS", "NS","NS", "**", "***"),
  tip_length = 0, color = "black")

#Export PDF 5.94 x 6.34 Landscape
```
