## Supplementary material for "Disordered proteins interact with the chemical environment to tune their protective function during drying": File S3: 1F.R

```
#Set working directory----
setwd('~/Desktop/')

#Import necessary packages----
library(ggplot2)
library(dplyr)
library(readxl)
library(ggsignif)

#Import your dataset----
data <- read_xlsx('Figure_1E_Peptide Synergy_with_Sugars.xlsx')
#Change the factor order
data$Peptide <- factor(data$Peptide,levels = c('At11',
                                               'At22',
                                               'At44',
                                               'Av11',
                                               'He11',
                                               'At20',
                                               'Aav11'))
#build ggplot environment
ggplot(data = data, aes(x = `%synergy_tre`, y = `%synergy_suc`, fill = Peptide))+
  geom_point(size = 6.5, shape = 21, color = 'black')+
  theme_bw()+
  theme(panel.grid = element_blank(),
        plot.title = element_text(hjust = 0.05, size = 14, vjust = -8),
        axis.title = element_text(size =14),
        axis.text.y = element_text(size =12),
        axis.text.x = element_text(size =12),
        legend.text=element_text(size=14), 
        legend.position = 'none', aspect.ratio = 1,
        panel.border = element_rect(colour = "black", fill=NA, size=1))+
  scale_color_manual(values = c(
    'skyblue3', 'dodgerblue3', 'turquoise3', 'yellow3', 'mediumpurple3', 'coral3','seagreen3'))+
  geom_abline(linetype = 'dashed',
              linewidth = 1)+
  geom_errorbar(aes(ymin = `%synergy_suc` - `%synergy_suc_err`,
                    ymax = `%synergy_suc` + `%synergy_suc_err`),
                width = 1, position = position_dodge(.9))+
  geom_errorbarh(aes(xmin = `%synergy_tre` - `%synergy_tre_err`,
                     xmax = `%synergy_tre` + `%synergy_tre_err`),
                 height = 1, position = position_dodge(.9))+
  labs(
    x = 'Synergy with Trehalose (%)',
    y = 'Synergy with Sucrose (%)'
  )+
  scale_y_continuous(
    breaks = seq(0,45,20),
    limits = c(-20,45))+
  xlim(-40,50)
```
