## Supplementary material for "Disordered proteins interact with the chemical environment to tune their protective function during drying": File S3: Figure_2B_FL_PD50.R

```
setwd('~/Documents/Research_Projects/LEA_CAHS_Oligomerization')

#call in packages
library(tidyverse)
library(dplyr)
library(purrr)
library(broom)
library(stringr)
library(ggplot2)
library(plotly)
library(readxl)
library(RColorBrewer)
library(patchwork)
library(grid)
library(ggsignif)
library(scales)

#FL Proteins
PD50_FL <- read_xlsx('Figure_1_PD50.xlsx', sheet = 'FL Proteins')
PD50_FL$Additive <- factor(PD50_FL$Additive, levels = c('CAHS D', 'HeLEA68614','AvLEA1C', 'BSA', 'AtLEA3-3', 'AavLEA1', 'AtLEA4-2', 'Trehalose','Sucrose'))
#make graph 
FL <-ggplot(data = PD50_FL, aes(x = Additive, y = PD50*100, fill = Additive))+
  geom_col(position = position_dodge(width = .9), color = 'black', size = 0.25)+
  theme_bw()+
  theme(
    plot.title = element_text(hjust = .5, size = 8),
    axis.title.y = element_text(size =8),
    axis.text.y = element_text(size =8),
    axis.title.x = element_text(size = 8),
    axis.text.x = element_text(size =8, angle = 45, hjust = 1),
    legend.text=element_text(size=14),
    legend.position = 'none', aspect.ratio = 1,
    legend.title = element_blank(),
    panel.border = element_rect(colour = "black", fill=NA, size=1))+
  labs(x = '',
       y = expression('PD50 (mM)'),
       title = '')+
  scale_fill_manual(values = c('#797979','#9337FF','#918F00','#CD91CD','#6BA5C8','#5ABA56','#CD5B45','#0433FF','#FF9200'))+
  geom_errorbar(aes(ymin = PD50 - STD, ymax = PD50 + STD), width = .2, position = position_dodge(.9))+
  scale_y_continuous(trans = 'log10', limits = c(1,10000),
                     breaks = c(1,10,100,1000,10000),
                     labels = c(0.01,0.1,1,10,100))+
  geom_errorbar(aes(ymin = PD50*100 - STD*100, ymax = PD50*100 + STD*100), width = .2, position = position_dodge(.9))
```
