## Supplementary material for "Disordered proteins interact with the chemical environment to tune their protective function during drying": File S3: Figure_2_All_CD_Graphs.R

```
#Set working directory----
setwd('~/Documents/LEA_CAHS_Oligomerization/')

#Import necessary packages----
library(ggplot2)
library(dplyr)
library(readxl)
library(ggsignif)
library(tidyverse)
library(ggsci)

#read in data----
data <- read_xlsx('Figure_2_CD_Data.xlsx')
#tidy up the data----
#filter out rows 262-288
data <- data[-c(262:288),]
#pivot columns 9-709 to wavelength
data <- data %>% pivot_longer(cols = 9:709,
                              names_to = 'Wavelength',
                              values_to = 'Absorbance')
#make a list of unique protein names
data_protein_list <- unique(data$`pep`)
#change absorbance values into numeric class
data$Wavelength <- as.numeric(data$Wavelength)
#select important groups
data <- data %>% dplyr::select(c('pep', 'sol', 'state', 'Wavelength', 'Absorbance'))
data$sol <- factor(data$sol, levels = c('None', 'Trehalose', 'Sucrose'))
#CAHSD----
  r <- data %>% filter(pep == 'CAHS_D') #filter out your protein
  r_d <- r %>% filter(state == 'Desiccated') #filter only desiccated
  r_h <- r %>% filter(state == 'Aqueous') #filter only hydrated samples
  r_h_buffer <- r_h %>% filter(sol == 'None') #filter only buffer
  r_d_buffer <- r_d %>% filter(sol == 'None') #filter only buffer
  r_h_Trehalose <- r_h %>% filter(sol == 'Trehalose') #filter only Trehalose
  r_d_Trehalose <- r_d %>% filter(sol == 'Trehalose') #filter only Trehalose
  r_h_Sucrose <- r_h %>% filter(sol == 'Sucrose') #filter only Sucrose
  r_d_Sucrose <- r_d %>% filter(sol == 'Sucrose') #filter only Sucrose
  #Calculate means for each condition
  r_h_buffer_mean <- r_h_buffer %>% group_by(state,sol, Wavelength) %>% summarise(mean = mean(Absorbance))
  r_d_buffer_mean <- r_d_buffer %>% group_by(state,sol,Wavelength) %>% summarise(mean = mean(Absorbance))
  r_h_Trehalose_mean <- r_h_Trehalose %>% group_by(state,sol,Wavelength) %>% summarise(mean = mean(Absorbance))
  r_d_Trehalose_mean <- r_d_Trehalose %>% group_by(state,sol,Wavelength) %>% summarise(mean = mean(Absorbance))
  r_h_Sucrose_mean <- r_h_Sucrose %>% group_by(state,sol,Wavelength) %>% summarise(mean = mean(Absorbance))
  r_d_Sucrose_mean <- r_d_Sucrose %>% group_by(state,sol,Wavelength) %>% summarise(mean = mean(Absorbance))
  
  #create plot - colors grouped by solution environment
  CAHSD_H <- ggplot(data = r_h_buffer_mean, aes(x = Wavelength, y = mean ,group = sol))+
    #create plot with all of hydrated buffer
    geom_line(data = r_h_buffer_mean, aes(x = Wavelength, y = mean, group = state, color = sol))+
    geom_line(data = r_h_Trehalose_mean, aes(x = Wavelength, y = mean, group = state, color = sol))+
    geom_line(data = r_h_Sucrose_mean, aes(x = Wavelength, y = mean, group = state, color = sol))+
    
    geom_line(data = r_h_buffer, aes(x = Wavelength, y = Absorbance, group = state, color = sol), alpha = .08)+
    geom_line(data = r_h_Trehalose, aes(x = Wavelength, y = Absorbance, group = state, color = sol), alpha = .08)+
    geom_line(data = r_h_Sucrose, aes(x = Wavelength, y = Absorbance, group = state, color = sol), alpha = .08)+
    
    #adjust graph paramaters
    theme_bw()+
    theme(panel.grid = element_blank(),
          plot.title = element_text(hjust = .9, vjust = -10, size = 8),
          axis.title = element_text(size =6),
          axis.text = element_text(size =6),
          legend.text=element_text(size=14), 
          legend.position = 'none', aspect.ratio = 1.25,
          panel.border = element_rect(colour = "black", fill=NA, size=1))+
    labs(x = 'Wavelength (nm)',
         y = '',
         title = 'CAHS_D')+
    #manually select for colors
    scale_color_manual(values = c('black', '#448B74', '#36628B'))+
    #set graph limits
    xlim(190,250)

  #desiccated samples
  CAHSD_D <- ggplot(data = r_h_buffer_mean, aes(x = Wavelength, y = mean ,group = sol))+
    #create plot with all of hydrated buffer
    geom_line(data = r_d_Sucrose_mean, aes(x = Wavelength, y = mean, group = state, color = sol))+
    geom_line(data = r_d_buffer_mean, aes(x = Wavelength, y = mean, group = state, color = sol))+
    geom_line(data = r_d_Trehalose_mean, aes(x = Wavelength, y = mean, group = state, color = sol))+
    
    geom_line(data = r_d_buffer, aes(x = Wavelength, y = Absorbance, group = state, color = sol), alpha = .08)+
    geom_line(data = r_d_Trehalose, aes(x = Wavelength, y = Absorbance, group = state, color = sol), alpha = .08)+
    geom_line(data = r_d_Sucrose, aes(x = Wavelength, y = Absorbance, group = state, color = sol), alpha = .08)+
    
    #adjust graph paramaters
    theme_bw()+
    theme(panel.grid = element_blank(),
          plot.title = element_text(hjust = .9, vjust = -10, size = 8),
          axis.title = element_text(size =6),
          axis.text = element_text(size =6),
          legend.text=element_text(size=14), 
          legend.position = 'none', aspect.ratio = 1.25,
          panel.border = element_rect(colour = "black", fill=NA, size=1))+
    labs(x = 'Wavelength (nm)',
         y = '',
         title = 'CAHS_D')+
    #manually select for colors
    scale_color_manual(values = c('black', '#448B74', '#36628B'))+
    #set graph limits
    xlim(190,250)

#AtLEA42----
r <- data %>% filter(pep == 'AtLEA4-2') #filter out your protein
r_d <- r %>% filter(state == 'Desiccated') #filter only desiccated
r_h <- r %>% filter(state == 'Aqueous') #filter only hydrated samples
r_h_buffer <- r_h %>% filter(sol == 'None') #filter only buffer
r_d_buffer <- r_d %>% filter(sol == 'None') #filter only buffer
r_h_Trehalose <- r_h %>% filter(sol == 'Trehalose') #filter only Trehalose
r_d_Trehalose <- r_d %>% filter(sol == 'Trehalose') #filter only Trehalose
r_h_Sucrose <- r_h %>% filter(sol == 'Sucrose') #filter only Sucrose
r_d_Sucrose <- r_d %>% filter(sol == 'Sucrose') #filter only Sucrose
#Calculate means for each condition
r_h_buffer_mean <- r_h_buffer %>% group_by(state,sol, Wavelength) %>% summarise(mean = mean(Absorbance))
r_d_buffer_mean <- r_d_buffer %>% group_by(state,sol,Wavelength) %>% summarise(mean = mean(Absorbance))
r_h_Trehalose_mean <- r_h_Trehalose %>% group_by(state,sol,Wavelength) %>% summarise(mean = mean(Absorbance))
r_d_Trehalose_mean <- r_d_Trehalose %>% group_by(state,sol,Wavelength) %>% summarise(mean = mean(Absorbance))
r_h_Sucrose_mean <- r_h_Sucrose %>% group_by(state,sol,Wavelength) %>% summarise(mean = mean(Absorbance))
r_d_Sucrose_mean <- r_d_Sucrose %>% group_by(state,sol,Wavelength) %>% summarise(mean = mean(Absorbance))

#create plot - colors grouped by solution environment
AtLEA42_H <- ggplot(data = r_h_buffer_mean, aes(x = Wavelength, y = mean ,group = sol))+
  #create plot with all of hydrated buffer
  geom_line(data = r_h_buffer_mean, aes(x = Wavelength, y = mean, group = state, color = sol))+
  geom_line(data = r_h_Trehalose_mean, aes(x = Wavelength, y = mean, group = state, color = sol))+
  geom_line(data = r_h_Sucrose_mean, aes(x = Wavelength, y = mean, group = state, color = sol))+
  
  geom_line(data = r_h_buffer, aes(x = Wavelength, y = Absorbance, group = state, color = sol), alpha = .08)+
  geom_line(data = r_h_Trehalose, aes(x = Wavelength, y = Absorbance, group = state, color = sol), alpha = .08)+
  geom_line(data = r_h_Sucrose, aes(x = Wavelength, y = Absorbance, group = state, color = sol), alpha = .08)+
  
  #adjust graph paramaters
  theme_bw()+
  theme(panel.grid = element_blank(),
        plot.title = element_text(hjust = .9, vjust = -10, size = 8),
        axis.title = element_text(size =6),
        axis.text = element_text(size =6),
        legend.text=element_text(size=14), 
        legend.position = 'none', aspect.ratio = 1.25,
        panel.border = element_rect(colour = "black", fill=NA, size=1))+
  labs(x = 'Wavelength (nm)',
       y = '',
       title = 'AtLEA4-2')+
  #manually select for colors
  scale_color_manual(values = c('black', '#448B74', '#36628B'))+
  #set graph limits
  xlim(190,250)

#desiccated samples
AtLEA42_D <- ggplot(data = r_h_buffer_mean, aes(x = Wavelength, y = mean ,group = sol))+
  #create plot with all of hydrated buffer
  geom_line(data = r_d_Sucrose_mean, aes(x = Wavelength, y = mean, group = state, color = sol))+
  geom_line(data = r_d_buffer_mean, aes(x = Wavelength, y = mean, group = state, color = sol))+
  geom_line(data = r_d_Trehalose_mean, aes(x = Wavelength, y = mean, group = state, color = sol))+
  
  geom_line(data = r_d_buffer, aes(x = Wavelength, y = Absorbance, group = state, color = sol), alpha = .08)+
  geom_line(data = r_d_Trehalose, aes(x = Wavelength, y = Absorbance, group = state, color = sol), alpha = .08)+
  geom_line(data = r_d_Sucrose, aes(x = Wavelength, y = Absorbance, group = state, color = sol), alpha = .08)+
  
  #adjust graph paramaters
  theme_bw()+
  theme(panel.grid = element_blank(),
        plot.title = element_text(hjust = .9, vjust = -10, size = 8),
        axis.title = element_text(size =6),
        axis.text = element_text(size =6),
        legend.text=element_text(size=14), 
        legend.position = 'none', aspect.ratio = 1.25,
        panel.border = element_rect(colour = "black", fill=NA, size=1))+
  labs(x = 'Wavelength (nm)',
       y = '',
       title = 'AtLEA4-2')+
  #manually select for colors
  scale_color_manual(values = c('black', '#448B74', '#36628B'))+
  #set graph limits
  xlim(190,250)
#HeLEA68614----
r <- data %>% filter(pep == 'HeLEA68614') #filter out your protein
r_d <- r %>% filter(state == 'Desiccated') #filter only desiccated
r_h <- r %>% filter(state == 'Aqueous') #filter only hydrated samples
r_h_buffer <- r_h %>% filter(sol == 'None') #filter only buffer
r_d_buffer <- r_d %>% filter(sol == 'None') #filter only buffer
r_h_Trehalose <- r_h %>% filter(sol == 'Trehalose') #filter only Trehalose
r_d_Trehalose <- r_d %>% filter(sol == 'Trehalose') #filter only Trehalose
r_h_Sucrose <- r_h %>% filter(sol == 'Sucrose') #filter only Sucrose
r_d_Sucrose <- r_d %>% filter(sol == 'Sucrose') #filter only Sucrose
#Calculate means for each condition
r_h_buffer_mean <- r_h_buffer %>% group_by(state,sol, Wavelength) %>% summarise(mean = mean(Absorbance))
r_d_buffer_mean <- r_d_buffer %>% group_by(state,sol,Wavelength) %>% summarise(mean = mean(Absorbance))
r_h_Trehalose_mean <- r_h_Trehalose %>% group_by(state,sol,Wavelength) %>% summarise(mean = mean(Absorbance))
r_d_Trehalose_mean <- r_d_Trehalose %>% group_by(state,sol,Wavelength) %>% summarise(mean = mean(Absorbance))
r_h_Sucrose_mean <- r_h_Sucrose %>% group_by(state,sol,Wavelength) %>% summarise(mean = mean(Absorbance))
r_d_Sucrose_mean <- r_d_Sucrose %>% group_by(state,sol,Wavelength) %>% summarise(mean = mean(Absorbance))

#create plot - colors grouped by solution environment
HeLEA68614_H <- ggplot(data = r_h_buffer_mean, aes(x = Wavelength, y = mean ,group = sol))+
  #create plot with all of hydrated buffer
  geom_line(data = r_h_buffer_mean, aes(x = Wavelength, y = mean, group = state, color = sol))+
  geom_line(data = r_h_Trehalose_mean, aes(x = Wavelength, y = mean, group = state, color = sol))+
  geom_line(data = r_h_Sucrose_mean, aes(x = Wavelength, y = mean, group = state, color = sol))+
  
  geom_line(data = r_h_buffer, aes(x = Wavelength, y = Absorbance, group = state, color = sol), alpha = .08)+
  geom_line(data = r_h_Trehalose, aes(x = Wavelength, y = Absorbance, group = state, color = sol), alpha = .08)+
  geom_line(data = r_h_Sucrose, aes(x = Wavelength, y = Absorbance, group = state, color = sol), alpha = .08)+
  
  #adjust graph paramaters
  theme_bw()+
  theme(panel.grid = element_blank(),
        plot.title = element_text(hjust = .9, vjust = -10, size = 8),
        axis.title = element_text(size =6),
        axis.text = element_text(size =6),
        legend.text=element_text(size=14), 
        legend.position = 'none', aspect.ratio = 1.25,
        panel.border = element_rect(colour = "black", fill=NA, size=1))+
  labs(x = 'Wavelength (nm)',
       y = '',
       title = 'HeLEA68614')+
  #manually select for colors
  scale_color_manual(values = c('black', '#448B74', '#36628B'))+
  #set graph limits
  xlim(190,250)

#desiccated samples
HeLEA68614_D <- ggplot(data = r_h_buffer_mean, aes(x = Wavelength, y = mean ,group = sol))+
  #create plot with all of hydrated buffer
  geom_line(data = r_d_Sucrose_mean, aes(x = Wavelength, y = mean, group = state, color = sol))+
  geom_line(data = r_d_buffer_mean, aes(x = Wavelength, y = mean, group = state, color = sol))+
  geom_line(data = r_d_Trehalose_mean, aes(x = Wavelength, y = mean, group = state, color = sol))+
  
  geom_line(data = r_d_buffer, aes(x = Wavelength, y = Absorbance, group = state, color = sol), alpha = .08)+
  geom_line(data = r_d_Trehalose, aes(x = Wavelength, y = Absorbance, group = state, color = sol), alpha = .08)+
  geom_line(data = r_d_Sucrose, aes(x = Wavelength, y = Absorbance, group = state, color = sol), alpha = .08)+
  
  #adjust graph paramaters
  theme_bw()+
  theme(panel.grid = element_blank(),
        plot.title = element_text(hjust = .9, vjust = -10, size = 8),
        axis.title = element_text(size =6),
        axis.text = element_text(size =6),
        legend.text=element_text(size=14), 
        legend.position = 'none', aspect.ratio = 1.25,
        panel.border = element_rect(colour = "black", fill=NA, size=1))+
  labs(x = 'Wavelength (nm)',
       y = '',
       title = 'HeLEA68614')+
  #manually select for colors
  scale_color_manual(values = c('black', '#448B74', '#36628B'))+
  #set graph limits
  xlim(190,250)
#AvLEA1C----
r <- data %>% filter(pep == 'AvLEA1C') #filter out your protein
r_d <- r %>% filter(state == 'Desiccated') #filter only desiccated
r_h <- r %>% filter(state == 'Aqueous') #filter only hydrated samples
r_h_buffer <- r_h %>% filter(sol == 'None') #filter only buffer
r_d_buffer <- r_d %>% filter(sol == 'None') #filter only buffer
r_h_Trehalose <- r_h %>% filter(sol == 'Trehalose') #filter only Trehalose
r_d_Trehalose <- r_d %>% filter(sol == 'Trehalose') #filter only Trehalose
r_h_Sucrose <- r_h %>% filter(sol == 'Sucrose') #filter only Sucrose
r_d_Sucrose <- r_d %>% filter(sol == 'Sucrose') #filter only Sucrose
#Calculate means for each condition
r_h_buffer_mean <- r_h_buffer %>% group_by(state,sol, Wavelength) %>% summarise(mean = mean(Absorbance))
r_d_buffer_mean <- r_d_buffer %>% group_by(state,sol,Wavelength) %>% summarise(mean = mean(Absorbance))
r_h_Trehalose_mean <- r_h_Trehalose %>% group_by(state,sol,Wavelength) %>% summarise(mean = mean(Absorbance))
r_d_Trehalose_mean <- r_d_Trehalose %>% group_by(state,sol,Wavelength) %>% summarise(mean = mean(Absorbance))
r_h_Sucrose_mean <- r_h_Sucrose %>% group_by(state,sol,Wavelength) %>% summarise(mean = mean(Absorbance))
r_d_Sucrose_mean <- r_d_Sucrose %>% group_by(state,sol,Wavelength) %>% summarise(mean = mean(Absorbance))

#create plot - colors grouped by solution environment
AvLEA1C_H <- ggplot(data = r_h_buffer_mean, aes(x = Wavelength, y = mean ,group = sol))+
  #create plot with all of hydrated buffer
  geom_line(data = r_h_buffer_mean, aes(x = Wavelength, y = mean, group = state, color = sol))+
  geom_line(data = r_h_Trehalose_mean, aes(x = Wavelength, y = mean, group = state, color = sol))+
  geom_line(data = r_h_Sucrose_mean, aes(x = Wavelength, y = mean, group = state, color = sol))+
  
  geom_line(data = r_h_buffer, aes(x = Wavelength, y = Absorbance, group = state, color = sol), alpha = .08)+
  geom_line(data = r_h_Trehalose, aes(x = Wavelength, y = Absorbance, group = state, color = sol), alpha = .08)+
  geom_line(data = r_h_Sucrose, aes(x = Wavelength, y = Absorbance, group = state, color = sol), alpha = .08)+
  
  #adjust graph paramaters
  theme_bw()+
  theme(panel.grid = element_blank(),
        plot.title = element_text(hjust = .9, vjust = -10, size = 8),
        axis.title = element_text(size =6),
        axis.text = element_text(size =6),
        legend.text=element_text(size=14), 
        legend.position = 'none', aspect.ratio = 1.25,
        panel.border = element_rect(colour = "black", fill=NA, size=1))+
  labs(x = 'Wavelength (nm)',
       y = '',
       title = 'AvLEA1C')+
  #manually select for colors
  scale_color_manual(values = c('black', '#448B74', '#36628B'))+
  #set graph limits
  xlim(190,250)

#desiccated samples
AvLEA1C_D <- ggplot(data = r_h_buffer_mean, aes(x = Wavelength, y = mean ,group = sol))+
  #create plot with all of hydrated buffer
  geom_line(data = r_d_Sucrose_mean, aes(x = Wavelength, y = mean, group = state, color = sol))+
  geom_line(data = r_d_buffer_mean, aes(x = Wavelength, y = mean, group = state, color = sol))+
  geom_line(data = r_d_Trehalose_mean, aes(x = Wavelength, y = mean, group = state, color = sol))+
  
  geom_line(data = r_d_buffer, aes(x = Wavelength, y = Absorbance, group = state, color = sol), alpha = .08)+
  geom_line(data = r_d_Trehalose, aes(x = Wavelength, y = Absorbance, group = state, color = sol), alpha = .08)+
  geom_line(data = r_d_Sucrose, aes(x = Wavelength, y = Absorbance, group = state, color = sol), alpha = .08)+
  
  #adjust graph paramaters
  theme_bw()+
  theme(panel.grid = element_blank(),
        plot.title = element_text(hjust = .9, vjust = -10, size = 8),
        axis.title = element_text(size =6),
        axis.text = element_text(size =6),
        legend.text=element_text(size=14), 
        legend.position = 'none', aspect.ratio = 1.25,
        panel.border = element_rect(colour = "black", fill=NA, size=1))+
  labs(x = 'Wavelength (nm)',
       y = '',
       title = 'AvLEA1C')+
  #manually select for colors
  scale_color_manual(values = c('black', '#448B74', '#36628B'))+
  #set graph limits
  xlim(190,250)
#AavLEA1----
r <- data %>% filter(pep == 'AavLEA1') #filter out your protein
r_d <- r %>% filter(state == 'Desiccated') #filter only desiccated
r_h <- r %>% filter(state == 'Aqueous') #filter only hydrated samples
r_h_buffer <- r_h %>% filter(sol == 'None') #filter only buffer
r_d_buffer <- r_d %>% filter(sol == 'None') #filter only buffer
r_h_Trehalose <- r_h %>% filter(sol == 'Trehalose') #filter only Trehalose
r_d_Trehalose <- r_d %>% filter(sol == 'Trehalose') #filter only Trehalose
r_h_Sucrose <- r_h %>% filter(sol == 'Sucrose') #filter only Sucrose
r_d_Sucrose <- r_d %>% filter(sol == 'Sucrose') #filter only Sucrose
#Calculate means for each condition
r_h_buffer_mean <- r_h_buffer %>% group_by(state,sol, Wavelength) %>% summarise(mean = mean(Absorbance))
r_d_buffer_mean <- r_d_buffer %>% group_by(state,sol,Wavelength) %>% summarise(mean = mean(Absorbance))
r_h_Trehalose_mean <- r_h_Trehalose %>% group_by(state,sol,Wavelength) %>% summarise(mean = mean(Absorbance))
r_d_Trehalose_mean <- r_d_Trehalose %>% group_by(state,sol,Wavelength) %>% summarise(mean = mean(Absorbance))
r_h_Sucrose_mean <- r_h_Sucrose %>% group_by(state,sol,Wavelength) %>% summarise(mean = mean(Absorbance))
r_d_Sucrose_mean <- r_d_Sucrose %>% group_by(state,sol,Wavelength) %>% summarise(mean = mean(Absorbance))

#create plot - colors grouped by solution environment
AavLEA1_H <- ggplot(data = r_h_buffer_mean, aes(x = Wavelength, y = mean ,group = sol))+
  #create plot with all of hydrated buffer
  geom_line(data = r_h_buffer_mean, aes(x = Wavelength, y = mean, group = state, color = sol))+
  geom_line(data = r_h_Trehalose_mean, aes(x = Wavelength, y = mean, group = state, color = sol))+
  geom_line(data = r_h_Sucrose_mean, aes(x = Wavelength, y = mean, group = state, color = sol))+
  
  geom_line(data = r_h_buffer, aes(x = Wavelength, y = Absorbance, group = state, color = sol), alpha = .08)+
  geom_line(data = r_h_Trehalose, aes(x = Wavelength, y = Absorbance, group = state, color = sol), alpha = .08)+
  geom_line(data = r_h_Sucrose, aes(x = Wavelength, y = Absorbance, group = state, color = sol), alpha = .08)+
  
  #adjust graph paramaters
  theme_bw()+
  theme(panel.grid = element_blank(),
        plot.title = element_text(hjust = .9, vjust = -10, size = 8),
        axis.title = element_text(size =6),
        axis.text = element_text(size =6),
        legend.text=element_text(size=14), 
        legend.position = 'none', aspect.ratio = 1.25,
        panel.border = element_rect(colour = "black", fill=NA, size=1))+
  labs(x = 'Wavelength (nm)',
       y = '',
       title = 'AavLEA1')+
  #manually select for colors
  scale_color_manual(values = c('black', '#448B74', '#36628B'))+
  #set graph limits
  xlim(190,250)

#desiccated samples
AavLEA1_D <- ggplot(data = r_h_buffer_mean, aes(x = Wavelength, y = mean ,group = sol))+
  #create plot with all of hydrated buffer
  geom_line(data = r_d_Sucrose_mean, aes(x = Wavelength, y = mean, group = state, color = sol))+
  geom_line(data = r_d_buffer_mean, aes(x = Wavelength, y = mean, group = state, color = sol))+
  geom_line(data = r_d_Trehalose_mean, aes(x = Wavelength, y = mean, group = state, color = sol))+
  
  geom_line(data = r_d_buffer, aes(x = Wavelength, y = Absorbance, group = state, color = sol), alpha = .08)+
  geom_line(data = r_d_Trehalose, aes(x = Wavelength, y = Absorbance, group = state, color = sol), alpha = .08)+
  geom_line(data = r_d_Sucrose, aes(x = Wavelength, y = Absorbance, group = state, color = sol), alpha = .08)+
  
  #adjust graph paramaters
  theme_bw()+
  theme(panel.grid = element_blank(),
        plot.title = element_text(hjust = .9, vjust = -10, size = 8),
        axis.title = element_text(size =6),
        axis.text = element_text(size =6),
        legend.text=element_text(size=14), 
        legend.position = 'none', aspect.ratio = 1.25,
        panel.border = element_rect(colour = "black", fill=NA, size=1))+
  labs(x = 'Wavelength (nm)',
       y = '',
       title = 'AavLEA1')+
  #manually select for colors
  scale_color_manual(values = c('black', '#448B74', '#36628B'))+
  #set graph limits
  xlim(190,250)
#AtLEA3-3----
r <- data %>% filter(pep == 'AtLEA3-3') #filter out your protein
r_d <- r %>% filter(state == 'Desiccated') #filter only desiccated
r_h <- r %>% filter(state == 'Aqueous') #filter only hydrated samples
r_h_buffer <- r_h %>% filter(sol == 'None') #filter only buffer
r_d_buffer <- r_d %>% filter(sol == 'None') #filter only buffer
r_h_Trehalose <- r_h %>% filter(sol == 'Trehalose') #filter only Trehalose
r_d_Trehalose <- r_d %>% filter(sol == 'Trehalose') #filter only Trehalose
r_h_Sucrose <- r_h %>% filter(sol == 'Sucrose') #filter only Sucrose
r_d_Sucrose <- r_d %>% filter(sol == 'Sucrose') #filter only Sucrose
#Calculate means for each condition
r_h_buffer_mean <- r_h_buffer %>% group_by(state,sol, Wavelength) %>% summarise(mean = mean(Absorbance))
r_d_buffer_mean <- r_d_buffer %>% group_by(state,sol,Wavelength) %>% summarise(mean = mean(Absorbance))
r_h_Trehalose_mean <- r_h_Trehalose %>% group_by(state,sol,Wavelength) %>% summarise(mean = mean(Absorbance))
r_d_Trehalose_mean <- r_d_Trehalose %>% group_by(state,sol,Wavelength) %>% summarise(mean = mean(Absorbance))
r_h_Sucrose_mean <- r_h_Sucrose %>% group_by(state,sol,Wavelength) %>% summarise(mean = mean(Absorbance))
r_d_Sucrose_mean <- r_d_Sucrose %>% group_by(state,sol,Wavelength) %>% summarise(mean = mean(Absorbance))

#create plot - colors grouped by solution environment
AtLEA33_H <- ggplot(data = r_h_buffer_mean, aes(x = Wavelength, y = mean ,group = sol))+
  #create plot with all of hydrated buffer
  geom_line(data = r_h_buffer_mean, aes(x = Wavelength, y = mean, group = state, color = sol))+
  geom_line(data = r_h_Trehalose_mean, aes(x = Wavelength, y = mean, group = state, color = sol))+
  geom_line(data = r_h_Sucrose_mean, aes(x = Wavelength, y = mean, group = state, color = sol))+
  
  geom_line(data = r_h_buffer, aes(x = Wavelength, y = Absorbance, group = state, color = sol), alpha = .08)+
  geom_line(data = r_h_Trehalose, aes(x = Wavelength, y = Absorbance, group = state, color = sol), alpha = .08)+
  geom_line(data = r_h_Sucrose, aes(x = Wavelength, y = Absorbance, group = state, color = sol), alpha = .08)+
  
  #adjust graph paramaters
  theme_bw()+
  theme(panel.grid = element_blank(),
        plot.title = element_text(hjust = .9, vjust = -10, size = 8),
        axis.title = element_text(size =6),
        axis.text = element_text(size =6),
        legend.text=element_text(size=14), 
        legend.position = 'none', aspect.ratio = 1.25,
        panel.border = element_rect(colour = "black", fill=NA, size=1))+
  labs(x = 'Wavelength (nm)',
       y = '',
       title = 'AtLEA3-3')+
  #manually select for colors
  scale_color_manual(values = c('black', '#448B74', '#36628B'))+
  #set graph limits
  xlim(190,250)

#desiccated samples
AtLEA33_D <- ggplot(data = r_h_buffer_mean, aes(x = Wavelength, y = mean ,group = sol))+
  #create plot with all of hydrated buffer
  geom_line(data = r_d_Sucrose_mean, aes(x = Wavelength, y = mean, group = state, color = sol))+
  geom_line(data = r_d_buffer_mean, aes(x = Wavelength, y = mean, group = state, color = sol))+
  geom_line(data = r_d_Trehalose_mean, aes(x = Wavelength, y = mean, group = state, color = sol))+
  
  geom_line(data = r_d_buffer, aes(x = Wavelength, y = Absorbance, group = state, color = sol), alpha = .08)+
  geom_line(data = r_d_Trehalose, aes(x = Wavelength, y = Absorbance, group = state, color = sol), alpha = .08)+
  geom_line(data = r_d_Sucrose, aes(x = Wavelength, y = Absorbance, group = state, color = sol), alpha = .08)+
  
  #adjust graph paramaters
  theme_bw()+
  theme(panel.grid = element_blank(),
        plot.title = element_text(hjust = .9, vjust = -10, size = 8),
        axis.title = element_text(size =6),
        axis.text = element_text(size =6),
        legend.text=element_text(size=14), 
        legend.position = 'none', aspect.ratio = 1.25,
        panel.border = element_rect(colour = "black", fill=NA, size=1))+
  labs(x = 'Wavelength (nm)',
       y = '',
       title = 'AtLEA3-3')+
  #manually select for colors
  scale_color_manual(values = c('black', '#448B74', '#36628B'))+
  #set graph limits
  xlim(190,250)
#AtLEA3-3----
r <- data %>% filter(pep == 'BSA') #filter out your protein
r_d <- r %>% filter(state == 'Desiccated') #filter only desiccated
r_h <- r %>% filter(state == 'Aqueous') #filter only hydrated samples
r_h_buffer <- r_h %>% filter(sol == 'None') #filter only buffer
r_d_buffer <- r_d %>% filter(sol == 'None') #filter only buffer
r_h_Trehalose <- r_h %>% filter(sol == 'Trehalose') #filter only Trehalose
r_d_Trehalose <- r_d %>% filter(sol == 'Trehalose') #filter only Trehalose
r_h_Sucrose <- r_h %>% filter(sol == 'Sucrose') #filter only Sucrose
r_d_Sucrose <- r_d %>% filter(sol == 'Sucrose') #filter only Sucrose
#Calculate means for each condition
r_h_buffer_mean <- r_h_buffer %>% group_by(state,sol, Wavelength) %>% summarise(mean = mean(Absorbance))
r_d_buffer_mean <- r_d_buffer %>% group_by(state,sol,Wavelength) %>% summarise(mean = mean(Absorbance))
r_h_Trehalose_mean <- r_h_Trehalose %>% group_by(state,sol,Wavelength) %>% summarise(mean = mean(Absorbance))
r_d_Trehalose_mean <- r_d_Trehalose %>% group_by(state,sol,Wavelength) %>% summarise(mean = mean(Absorbance))
r_h_Sucrose_mean <- r_h_Sucrose %>% group_by(state,sol,Wavelength) %>% summarise(mean = mean(Absorbance))
r_d_Sucrose_mean <- r_d_Sucrose %>% group_by(state,sol,Wavelength) %>% summarise(mean = mean(Absorbance))

#create plot - colors grouped by solution environment
BSA_H <- ggplot(data = r_h_buffer_mean, aes(x = Wavelength, y = mean ,group = sol))+
  #create plot with all of hydrated buffer
  geom_line(data = r_h_buffer_mean, aes(x = Wavelength, y = mean, group = state, color = sol))+
  geom_line(data = r_h_Trehalose_mean, aes(x = Wavelength, y = mean, group = state, color = sol))+
  geom_line(data = r_h_Sucrose_mean, aes(x = Wavelength, y = mean, group = state, color = sol))+
  
  geom_line(data = r_h_buffer, aes(x = Wavelength, y = Absorbance, group = state, color = sol), alpha = .08)+
  geom_line(data = r_h_Trehalose, aes(x = Wavelength, y = Absorbance, group = state, color = sol), alpha = .08)+
  geom_line(data = r_h_Sucrose, aes(x = Wavelength, y = Absorbance, group = state, color = sol), alpha = .08)+
  
  #adjust graph paramaters
  theme_bw()+
  theme(panel.grid = element_blank(),
        plot.title = element_text(hjust = .9, vjust = -10, size = 8),
        axis.title = element_text(size =6),
        axis.text = element_text(size =6),
        legend.text=element_text(size=14), 
        legend.position = 'none', aspect.ratio = 1.25,
        panel.border = element_rect(colour = "black", fill=NA, size=1))+
  labs(x = 'Wavelength (nm)',
       y = '',
       title = 'BSA')+
  #manually select for colors
  scale_color_manual(values = c('black', '#448B74', '#36628B'))+
  #set graph limits
  xlim(190,250)

#desiccated samples
BSA_D <- ggplot(data = r_h_buffer_mean, aes(x = Wavelength, y = mean ,group = sol))+
  #create plot with all of hydrated buffer
  geom_line(data = r_d_Sucrose_mean, aes(x = Wavelength, y = mean, group = state, color = sol))+
  geom_line(data = r_d_buffer_mean, aes(x = Wavelength, y = mean, group = state, color = sol))+
  geom_line(data = r_d_Trehalose_mean, aes(x = Wavelength, y = mean, group = state, color = sol))+
  
  geom_line(data = r_d_buffer, aes(x = Wavelength, y = Absorbance, group = state, color = sol), alpha = .08)+
  geom_line(data = r_d_Trehalose, aes(x = Wavelength, y = Absorbance, group = state, color = sol), alpha = .08)+
  geom_line(data = r_d_Sucrose, aes(x = Wavelength, y = Absorbance, group = state, color = sol), alpha = .08)+
  
  #adjust graph paramaters
  theme_bw()+
  theme(panel.grid = element_blank(),
        plot.title = element_text(hjust = .9, vjust = -10, size = 8),
        axis.title = element_text(size =6),
        axis.text = element_text(size =6),
        legend.text=element_text(size=14), 
        legend.position = 'none', aspect.ratio = 1.25,
        panel.border = element_rect(colour = "black", fill=NA, size=1))+
  labs(x = 'Wavelength (nm)',
       y = '',
       title = 'BSA')+
  #manually select for colors
  scale_color_manual(values = c('black', '#448B74', '#36628B'))+
  #set graph limits
  xlim(190,250)
#Patchwork---- 3x 5.1 export----
(AtLEA33_H + AavLEA1_H + HeLEA68614_H + AvLEA1C_H + AtLEA42_H + BSA_H + CAHSD_H) + plot_layout(ncol = 4)
(AtLEA33_D + AavLEA1_D + HeLEA68614_D + AvLEA1C_D + AtLEA42_D + BSA_D + CAHSD_D) + plot_layout(ncol = 4)
```
