## Supplementary material for "Disordered proteins interact with the chemical environment to tune their protective function during drying": File S3: Figure_2_CD_Barchart.R

```
#Set working directory----
setwd('~/Documents/Research_Projects/LEA_CAHS_Oligomerization/')

#Import necessary packages----
library(ggplot2)
library(dplyr)
library(readxl)
library(ggsignif)
library(tidyverse)
library(ggsci)

#Read in data----
data <- read_xlsx('Figure_2_CD_State_Bar.xlsx', sheet = 'Sheet2')

#Set factor levels----
data$Solvent <- factor(data$Solvent, levels = c(
  'Buffer',
  'Trehalose',
  'Sucrose'))

data$Condition <- factor(data$Condition, levels = c('Hydrated',
                                                    'Dehydrated'))

#CAHS_D----
  r <- data %>% filter(Protein == 'CAHS_D')
  CAHS_D_bar<-ggplot(data = r, aes(x = Condition, y = Ratio, fill = Solvent))+
    geom_bar(stat = 'identity', position = 'dodge', color ='black', size = .3)+
    theme_bw()+
    theme(panel.grid = element_blank(),
          plot.title = element_text(vjust = -10, size = 6, hjust = .05),
          axis.title = element_text(size =6),
          axis.text = element_text(size =6),
          legend.text=element_text(size=12), 
          legend.position = 'none', aspect.ratio = 1.25,
          panel.border = element_rect(colour = "black", fill=NA, size=1))+
    labs(x = '',
         y = '',
         title = 'CAHS_D')+
    #manually select for colors
    scale_fill_manual(values = c('black', '#36628B', '#448B74'))+

    #change y continuos scaling
    scale_y_continuous(limits = c(0,2.5))+
    #add error bars based on standard deviation
    geom_errorbar(aes(ymin = Ratio - Error, ymax = Ratio + Error), width = .2, position = position_dodge(.9))+
    scale_x_discrete(labels = c('Hydrated' = 'Hydrated',
                                'Dehydrated' = "Dry") )+
    geom_signif(
      y_position= c(2, 2), xmin=c(0.7, 1.7), xmax=c(1.3, 2.3),
      annotations =c("NS", "NS"), tip_length= 0 , textsize = 2)
  #AtLEA3-3----
  r <- data %>% filter(Protein == 'AtLEA3-3')
  AtLEA33_bar<-ggplot(data = r, aes(x = Condition, y = Ratio, fill = Solvent))+
    geom_bar(stat = 'identity', position = 'dodge', color ='black', size = .3)+
    theme_bw()+
    theme(panel.grid = element_blank(),
          plot.title = element_text(vjust = -10, size = 6, hjust = .05),
          axis.title = element_text(size =6),
          axis.text = element_text(size =6),
          legend.text=element_text(size=12), 
          legend.position = 'none', aspect.ratio = 1.25,
          panel.border = element_rect(colour = "black", fill=NA, size=1))+
    labs(x = '',
         y = '',
         title = 'AtLEA3-3')+
    #manually select for colors
    scale_fill_manual(values = c('black', '#36628B', '#448B74'))+
    
    #change y continuos scaling
    scale_y_continuous(limits = c(0,2.5))+
    #add error bars based on standard deviation
    geom_errorbar(aes(ymin = Ratio - Error, ymax = Ratio + Error), width = .2, position = position_dodge(.9))+
    scale_x_discrete(labels = c('Hydrated' = 'Hydrated',
                                'Dehydrated' = "Dry") )+
    geom_signif(
      y_position= c(2, 2), xmin=c(0.7, 1.7), xmax=c(1.3, 2.3),
      annotations =c("NS", "NS"), tip_length= 0 , textsize = 2)
  #AtLEA4-2----
  r <- data %>% filter(Protein == 'AtLEA4-2')
  AtLEA42_bar<-ggplot(data = r, aes(x = Condition, y = Ratio, fill = Solvent))+
    geom_bar(stat = 'identity', position = 'dodge', color ='black', size = .3)+
    theme_bw()+
    theme(panel.grid = element_blank(),
          plot.title = element_text(vjust = -10, size = 6, hjust = .05),
          axis.title = element_text(size =6),
          axis.text = element_text(size =6),
          legend.text=element_text(size=12), 
          legend.position = 'none', aspect.ratio = 1.25,
          panel.border = element_rect(colour = "black", fill=NA, size=1))+
    labs(x = '',
         y = '',
         title = 'AtLEA4-2')+
    #manually select for colors
    scale_fill_manual(values = c('black', '#36628B', '#448B74'))+
    
    #change y continuos scaling
    scale_y_continuous(limits = c(0,2.5))+
    #add error bars based on standard deviation
    geom_errorbar(aes(ymin = Ratio - Error, ymax = Ratio + Error), width = .2, position = position_dodge(.9))+
    scale_x_discrete(labels = c('Hydrated' = 'Hydrated',
                                'Dehydrated' = "Dry") )+
    geom_signif(
      y_position= c(2, 2), xmin=c(0.7, 1.7), xmax=c(1.3, 2.3),
      annotations =c("NS", "NS"), tip_length= 0 , textsize = 2)
  #AvLEA1C----
  r <- data %>% filter(Protein == 'AvLEA1C')
  AvLEA1C_bar<-ggplot(data = r, aes(x = Condition, y = Ratio, fill = Solvent))+
    geom_bar(stat = 'identity', position = 'dodge', color ='black', size = .3)+
    theme_bw()+
    theme(panel.grid = element_blank(),
          plot.title = element_text(vjust = -10, size = 6, hjust = .05),
          axis.title = element_text(size =6),
          axis.text = element_text(size =6),
          legend.text=element_text(size=12), 
          legend.position = 'none', aspect.ratio = 1.25,
          panel.border = element_rect(colour = "black", fill=NA, size=1))+
    labs(x = '',
         y = '',
         title = 'AvLEA1C')+
    #manually select for colors
    scale_fill_manual(values = c('black', '#36628B', '#448B74'))+
    
    #change y continuos scaling
    scale_y_continuous(limits = c(0,2.5))+
    #add error bars based on standard deviation
    geom_errorbar(aes(ymin = Ratio - Error, ymax = Ratio + Error), width = .2, position = position_dodge(.9))+
    scale_x_discrete(labels = c('Hydrated' = 'Hydrated',
                                'Dehydrated' = "Dry") )+
    geom_signif(
      y_position= c(2, 2), xmin=c(0.7, 1.7), xmax=c(1.3, 2.3),
      annotations =c("NS", "NS"), tip_length= 0 , textsize = 2)
  #AavLEA1----
  r <- data %>% filter(Protein == 'AavLEA1')
  AavLEA1_bar<-ggplot(data = r, aes(x = Condition, y = Ratio, fill = Solvent))+
    geom_bar(stat = 'identity', position = 'dodge', color ='black', size = .3)+
    theme_bw()+
    theme(panel.grid = element_blank(),
          plot.title = element_text(vjust = -10, size = 6, hjust = .05),
          axis.title = element_text(size =6),
          axis.text = element_text(size =6),
          legend.text=element_text(size=12), 
          legend.position = 'none', aspect.ratio = 1.25,
          panel.border = element_rect(colour = "black", fill=NA, size=1))+
    labs(x = '',
         y = '',
         title = 'AavLEA1')+
    #manually select for colors
    scale_fill_manual(values = c('black', '#36628B', '#448B74'))+
    
    #change y continuos scaling
    scale_y_continuous(limits = c(0,2.5))+
    #add error bars based on standard deviation
    geom_errorbar(aes(ymin = Ratio - Error, ymax = Ratio + Error), width = .2, position = position_dodge(.9))+
    scale_x_discrete(labels = c('Hydrated' = 'Hydrated',
                                'Dehydrated' = "Dry") )+
    geom_signif(
      y_position= c(2, 2), xmin=c(0.7, 1.7), xmax=c(1.3, 2.3),
      annotations =c("NS", "NS"), tip_length= 0 , textsize = 2)
  #HeLEA68614----
  r <- data %>% filter(Protein == 'HeLEA68614')
  HeLEA68614_bar<-ggplot(data = r, aes(x = Condition, y = Ratio, fill = Solvent))+
    geom_bar(stat = 'identity', position = 'dodge', color ='black', size = .3)+
    theme_bw()+
    theme(panel.grid = element_blank(),
          plot.title = element_text(vjust = -10, size = 6, hjust = .05),
          axis.title = element_text(size =6),
          axis.text = element_text(size =6),
          legend.text=element_text(size=12), 
          legend.position = 'none', aspect.ratio = 1.25,
          panel.border = element_rect(colour = "black", fill=NA, size=1))+
    labs(x = '',
         y = '',
         title = 'HeLEA68614')+
    #manually select for colors
    scale_fill_manual(values = c('black', '#36628B', '#448B74'))+
    
    #change y continuos scaling
    scale_y_continuous(limits = c(0,2.5))+
    #add error bars based on standard deviation
    geom_errorbar(aes(ymin = Ratio - Error, ymax = Ratio + Error), width = .2, position = position_dodge(.9))+
    scale_x_discrete(labels = c('Hydrated' = 'Hydrated',
                                'Dehydrated' = "Dry") )+
    geom_signif(
      y_position= c(2, 2), xmin=c(0.7, 1.7), xmax=c(1.3, 2.3),
      annotations =c("NS", "NS"), tip_length= 0 , textsize = 2)
  #BSA----
  r <- data %>% filter(Protein == 'BSA')
  BSA_bar<-ggplot(data = r, aes(x = Condition, y = Ratio, fill = Solvent))+
    geom_bar(stat = 'identity', position = 'dodge', color ='black', size = .3)+
    theme_bw()+
    theme(panel.grid = element_blank(),
          plot.title = element_text(vjust = -10, size = 6, hjust = .05),
          axis.title = element_text(size =6),
          axis.text = element_text(size =6),
          legend.text=element_text(size=12), 
          legend.position = 'none', aspect.ratio = 1.25,
          panel.border = element_rect(colour = "black", fill=NA, size=1))+
    labs(x = '',
         y = '',
         title = 'BSA')+
    #manually select for colors
    scale_fill_manual(values = c('black', '#36628B', '#448B74'))+
    
    #change y continuos scaling
    scale_y_continuous(limits = c(0,2.5))+
    #add error bars based on standard deviation
    geom_errorbar(aes(ymin = Ratio - Error, ymax = Ratio + Error), width = .2, position = position_dodge(.9))+
    scale_x_discrete(labels = c('Hydrated' = 'Hydrated',
                                'Dehydrated' = "Dry") )+
    geom_signif(
      y_position= c(2, 2), xmin=c(0.7, 1.7), xmax=c(1.3, 2.3),
      annotations =c("NS", "NS"), tip_length= 0 , textsize = 2)
  
(AtLEA33_bar+HeLEA68614_bar+AavLEA1_bar+AvLEA1C_bar+AtLEA42_bar+BSA_bar+CAHS_D_bar)+plot_layout(ncol = 4)
```
