## Supplementary material for "Disordered proteins interact with the chemical environment to tune their protective function during drying": File S3: SAXS Data Viz.R

```
####set working directory####
setwd('C:/Users/Vincent/OneDrive/Documents/Boothby Lab Work/Team CAHSD/Oligomerization Project')
#import necessary packages
library(ggplot2)
library(dplyr)
library(tidyverse)
library(readxl)
library(patchwork)
#read in data
SAXS <- read_excel("Oligomerization SAXS Detailed Guinier.xlsx")

Proteins <- c("BSA", "AavLEA", "AtLEA3-3", "HeLEA", "AvLEA", "AtLEA4-2", "CAHSD")

Rg_list <- c()
MW_list <- c()

for(i in 1:length(Proteins))
{
  protein = Proteins[i]
  
  CT_SAXS <- subset(SAXS, Protein == protein)
  CT_SAXS$Name <- paste(substring(CT_SAXS$CoSolute, 1, 1), " ", CT_SAXS$Ratio, ":1", sep = "")
  CT_SAXS$Name[1] <- "Tris"
  
  if(protein == "CAHSD")
  {
    ymax = 400
    vals = c("Tris", "S 1.6:1", "S 16:1", "S 160:1", "T 1.6:1", "T 16:1", "T 160:1")
    Colors <- c("#cccccc", "#8cb8de", "#6393bf", "#36628B", "#9cdbc7", "#63B89C", "#448B74")
    aspect = 0.8
  }else
  {
    ymax = 150
    vals = c("Tris","S 20:1","S 200:1", "T 20:1", "T 200:1")
    Colors <- c("#cccccc", "#6393bf", "#36628B", "#63B89C", "#448B74")
    aspect = 1
  }
  
  Rg <-ggplot(data = CT_SAXS, aes(x = fct_inorder(Name), y = Rg, fill = fct_inorder(Name)))+
    geom_bar(stat = 'identity', color = 'black')+
    geom_errorbar(aes(ymin = Rg - `Rg_+-`, ymax = Rg + `Rg_+-`), width = .2, position = position_dodge(.9))+
    scale_x_discrete(breaks = vals) +
    theme_bw() +
    theme(
      plot.title = element_text(hjust = 0.065, size = 8, vjust = -12),
      axis.title = element_text(size = 8),
      axis.text.y = element_text(size = 8),
      axis.text.x = element_text(size = 8, angle = 70, hjust = 1, vjust = 1),
      legend.text=element_text(size=14),
      legend.position = 'none', aspect.ratio = aspect,
      legend.title = element_blank(),
      panel.border = element_rect(colour = "black", fill=NA, size=1)) +
    labs(x = '', y = 'Radius of Gyration (Å)', title = protein) +
    scale_fill_manual(values = Colors) +
    ylim(0,80)
  
  Rg_list[[i]] <- Rg
  
  MW<- ggplot(data = CT_SAXS, aes(x = fct_inorder(Name), y = Bayes, fill = fct_inorder(Name)))+
    geom_bar(stat = 'identity', color = 'black')+
    geom_errorbar(aes(ymin = `Bayes_Low`, ymax = `Bayes_High`), width = .2, position = position_dodge(.9))+
    scale_x_discrete(breaks= vals) +
    theme_bw() +
    theme(
      plot.title = element_text(hjust = 0.05, size = 14, vjust = -8),
      axis.title = element_text(size = 8),
      axis.text.y = element_text(size =8),
      axis.text.x = element_text(size = 8, angle = 70, hjust = 1, vjust = 1),
      legend.text=element_text(size=14),
      legend.position = 'none', aspect.ratio = aspect,
      legend.title = element_blank(),
      panel.border = element_rect(colour = "black", fill=NA, size=1)) +
    labs(x = '', y = 'Predicted Molecular Weight (kDa)', title = '') +
    scale_fill_manual(values = Colors) +
    ylim(0,ymax)
  
  MW_list[[i]] <- MW
}

(Rg_list[[1]] | MW_list [[1]] | Rg_list[[2]] | MW_list [[2]]) / (Rg_list[[3]] | MW_list [[3]] | Rg_list[[4]] | MW_list [[4]]) / (Rg_list[[5]] | MW_list [[5]] | Rg_list[[6]] | MW_list [[6]]) / (Rg_list[[7]] | MW_list [[7]])
```
