## Supplementary material for "Disordered proteins interact with the chemical environment to tune their protective function during drying": File S3: Figure_5.R

```
setwd('~/Documents/Research_Projects/LEA_CAHS_Oligomerization/Submission')

#call in packages
library(tidyverse)
library(dplyr)
library(purrr)
library(broom)
library(stringr)
library(ggplot2)
library(plotly)
library(readxl)
library(RColorBrewer)
library(patchwork)

#read in data
T6 <- read_xlsx('Figure_5_DSC_Trehalose_Sucrose.xlsx', sheet = 'Trehalose')
colnames(T6)<- c('Temperature', 'Heat_Flow', 'Sample')
DSC_min <- T6 %>% 
  group_by(Sample)%>%
  summarise_at(vars(Heat_Flow),
               list(min = min))
#create df with discrete heat_flow values
Sample <- c('CT0', 'CT1', 'CT10', 'CT100', 'CT500')
adj_heatflow <- c(-.324, -.33, -.336, -.342, -.348)
adjusted_heatflow <- data.frame(Sample, adj_heatflow)
#combine dataframe
Adjusted <- inner_join(DSC_min, adjusted_heatflow)
#find difference
Adjusted <- Adjusted %>% mutate(Diff= adj_heatflow - min)
#select and merge with DSC 
Adjusted <- Adjusted %>% select(c(Sample, Diff))
#merge
Adjusted_DSC <- inner_join(T6, Adjusted)
Adjusted_DSC <- Adjusted_DSC %>% mutate(adjusted_heatflow = Heat_Flow + Diff)
#import data for enthalpy selection
Trehalose_markers <- read_xlsx('6mg_DSC_CTS.xlsx', sheet = 'Trehalose_points')
DSC_T6<-ggplot(data = Adjusted_DSC, aes(x = Temperature, y = adjusted_heatflow))+
  geom_line(aes(color = Sample))+
  geom_point(data = Trehalose_markers, aes(x = Temperature, y = adjusted_heatflow), shape = 3, size = 1)+
  ylim(-.35,-.31)+
  theme_bw()+
  theme(axis.text = element_text(size = 6),
        plot.title = element_text(hjust = .5, size = 6, vjust = -10),
        axis.title.y = element_text(size = 6),
        axis.title.x = element_text(size = 6),
        legend.position = 'none', aspect.ratio = 1,
        panel.border = element_rect(colour = "black", fill=NA, size=.5),
        panel.grid = element_blank(),
        axis.text.y = element_blank(),
        axis.ticks.y = element_blank())+
  labs(color = '',
       y = 'Heat flow (W/g)',
       title = 'CAHS D',
       x = 'Temperature (\u00B0C)')+
  xlim(20,55)+
  
  scale_color_manual(values = c('black', '#B0E0E6','#8CB8DE','#6393BF','#36628B'))

#read in data
S6 <- read_xlsx('Figure_5_DSC_Trehalose_Sucrose.xlsx', sheet = 'Sucrose')
colnames(S6)<- c('Temperature', 'Heat_Flow', 'Sample')
DSC_min <- S6 %>% 
  group_by(Sample)%>%
  summarise_at(vars(Heat_Flow),
               list(min = min))
#create df with discrete heat_flow values
Sample <- c('CS0', 'CS1', 'CS10', 'CS100', 'CS500')
adj_heatflow <- c(-.324, -.33, -.336, -.342, -.348)
adjusted_heatflow <- data.frame(Sample, adj_heatflow)
#combine dataframe
Adjusted <- inner_join(DSC_min, adjusted_heatflow)
#find difference
Adjusted <- Adjusted %>% mutate(Diff= adj_heatflow - min)
#select and merge with DSC 
Adjusted <- Adjusted %>% select(c(Sample, Diff))
#merge
Adjusted_DSC <- inner_join(S6, Adjusted)
Adjusted_DSC <- Adjusted_DSC %>% mutate(adjusted_heatflow = Heat_Flow + Diff)
#import data for enthalpy selection
Sucrose_markers <- read_xlsx('6mg_DSC_CTS.xlsx', sheet = 'Sucrose_points')

DSC_S6<-ggplot(data = Adjusted_DSC, aes(x = Temperature, y = adjusted_heatflow))+
  geom_line(aes(color = Sample))+
  geom_point(data = Sucrose_markers, aes(x = Temperature, y = adjusted_heatflow), shape = 3, size = 1)+
  
  ylim(-.35,-.31)+
  theme_bw()+
  theme(axis.text = element_text(size = 6),
        plot.title = element_text(hjust = 0.5, size = 6, vjust = -10),
        axis.title.y = element_text(size = 6),
        axis.title.x = element_text(size = 6),
        legend.position = 'none', aspect.ratio = 1,
        panel.border = element_rect(colour = "black", fill=NA, size=.5),
        panel.grid = element_blank(),
        axis.text.y = element_blank(),
        axis.ticks.y = element_blank())+
  labs(color = '',
       y = 'Heat flow (W/g)',
       title = 'CAHS D',
       x = 'Temperature (\u00B0C)')+
  xlim(20,55)+
  
  scale_color_manual(values = c('black', '#CAE7D3','#9CDBC7','#63B89C','#448B74'))
#Trehalose Enthalpy----
Treh_E <- read_xlsx('Figure_5_DSC_Trehalose_Sucrose.xlsx', sheet = 'CAHSD_Trehalose_6')
Treh_E <- Treh_E %>% filter(Sample %in% c('CT1', 'CT10', 'CT100', 'CT500'))
TE<-ggplot(data = Treh_E, aes(x = Sample, y = Avg_C_Enthalpy*1000, fill = Sample))+
  geom_bar(position = position_dodge(), stat = 'identity', color = 'black', size = .3)+
  theme_bw()+
  theme(axis.text = element_text(size = 6),
        plot.title = element_text(hjust = 0.5, size = 6, vjust = -10),
        axis.title.y = element_text(size = 6),
        axis.title.x = element_text(size = 6),
        legend.position = 'none', aspect.ratio = 1,
        panel.border = element_rect(colour = "black", fill=NA, size=.5))+
  labs(
    x = 'Molar Ratio (Trehalose:CAHS D)',
    y = expression(Delta*'Enthalpy (mJ/g)'),
    title = 'CAHS D')+
  scale_fill_manual(values = c('#B0E0E6','#8CB8DE','#6393BF','#36628B'))+
  scale_x_discrete(breaks=c("CT1","CT10", "CT100", "CT500"),
                   labels=c("1:1","10:1", "100:1", "500:1"))+
  geom_errorbar(aes(ymin = Avg_C_Enthalpy*1000 - Stdev_C_Enthalpy*1000, ymax = Avg_C_Enthalpy*1000 + Stdev_C_Enthalpy*1000), width = .2, position = position_dodge(.9), size = .3)+
  ylim(-5,50)

#Sucrose Enthalpy----
Sucr_E <- read_xlsx('Figure_5_DSC_Trehalose_Sucrose.xlsx', sheet = 'CAHSD_Sucrose_6')
Sucr_E <- Sucr_E %>% filter(Sample %in% c('CS1', 'CS10', 'CS100', 'CS500'))
SE<-ggplot(data = Sucr_E, aes(x = Sample, y = Avg_C_Enthalpy*1000, fill = Sample))+
  geom_bar(position = position_dodge(), stat = 'identity', color = 'black', size = .3)+
  theme_bw()+
  theme(axis.text = element_text(size = 6),
        plot.title = element_text(hjust = 0.5, size = 6, vjust = -10),
        axis.title.y = element_text(size = 6),
        axis.title.x = element_text(size = 6),
        legend.position = 'none', aspect.ratio = 1,
        panel.border = element_rect(colour = "black", fill=NA, size=.5))+
  labs(
    x = 'Molar Ratio (Trehalose:CAHS D)',
    y = expression(Delta*'Enthalpy (mJ/g)'),
    title = 'CAHS D')+
  scale_fill_manual(values = c( '#CAE7D3','#9CDBC7','#63B89C','#448B74'))+
  scale_x_discrete(breaks=c("CS1","CS10", "CS100", "CS500"),
                   labels=c("1:1","10:1", "100:1", "500:1"))+
  geom_errorbar(aes(ymin = Avg_C_Enthalpy*1000 - Stdev_C_Enthalpy*1000, ymax = Avg_C_Enthalpy*1000 + Stdev_C_Enthalpy*1000), width = .2, position = position_dodge(.9), size = .3)+
  ylim(-5,50)
#AavLEA1 Trehalose----
#read in data
AavLEA1T <- read_xlsx('Figure_5_DSC.xlsx', sheet = 'AavLEA1_Trehalose')
colnames(AavLEA1T)<- c('Temperature', 'Heat_Flow', 'Sample')
DSC_min <- AavLEA1T %>% 
  group_by(Sample)%>%
  summarise_at(vars(Heat_Flow),
               list(min = min))
#create df with discrete heat_flow values
Sample <- c('T0', 'T1', 'T10', 'T100', 'T500')
adj_heatflow <- c(-.324, -.33, -.336, -.342, -.348)
adjusted_heatflow <- data.frame(Sample, adj_heatflow)
#combine dataframe
Adjusted <- inner_join(DSC_min, adjusted_heatflow)
#find difference
Adjusted <- Adjusted %>% mutate(Diff= adj_heatflow - min)
#select and merge with DSC 
Adjusted <- Adjusted %>% select(c(Sample, Diff))
#merge
Adjusted_DSC <- inner_join(AavLEA1T, Adjusted)
Adjusted_DSC <- Adjusted_DSC %>% mutate(adjusted_heatflow = Heat_Flow + Diff)
#import data for enthalpy selection
DSC_AavLEA1T<-ggplot(data = Adjusted_DSC, aes(x = Temperature, y = adjusted_heatflow))+
  geom_line(aes(color = Sample))+
  ylim(-.35,-.31)+
  theme_bw()+
  theme(axis.text = element_text(size = 6),
        plot.title = element_text(hjust = .5, size = 6, vjust = -10),
        axis.title.y = element_text(size = 6),
        axis.title.x = element_text(size = 6),
        legend.position = 'none', aspect.ratio = 1,
        panel.border = element_rect(colour = "black", fill=NA, size=.5),
        panel.grid = element_blank(),
        axis.text.y = element_blank(),
        axis.ticks.y = element_blank())+
  labs(color = '',
       y = 'Heat flow (W/g)',
       title = 'AavLEA1',
       x = 'Temperature (\u00B0C)')+
  xlim(20,55)+
  
  scale_color_manual(values = c('black', '#B0E0E6','#8CB8DE','#6393BF','#36628B'))
#AavLEA1 Sucrose----
#read in data
AavLEA1S <- read_xlsx('Figure_5_DSC.xlsx', sheet = 'AavLEA1_Sucrose')
colnames(AavLEA1S)<- c('Temperature', 'Heat_Flow', 'Sample')
DSC_min <- AavLEA1S %>% 
  group_by(Sample)%>%
  summarise_at(vars(Heat_Flow),
               list(min = min))
#create df with discrete heat_flow values
Sample <- c('S0', 'S1', 'S10', 'S100', 'S500')
adj_heatflow <- c(-.324, -.333, -.336, -.342, -.348)
adjusted_heatflow <- data.frame(Sample, adj_heatflow)
#combine dataframe
Adjusted <- inner_join(DSC_min, adjusted_heatflow)
#find difference
Adjusted <- Adjusted %>% mutate(Diff= adj_heatflow - min)
#select and merge with DSC 
Adjusted <- Adjusted %>% select(c(Sample, Diff))
#merge
Adjusted_DSC <- inner_join(AavLEA1S, Adjusted)
Adjusted_DSC <- Adjusted_DSC %>% mutate(adjusted_heatflow = Heat_Flow + Diff)
#import data for enthalpy selection
DSC_AavLEA1S<-ggplot(data = Adjusted_DSC, aes(x = Temperature, y = adjusted_heatflow))+
  geom_line(aes(color = Sample))+
  ylim(-.35,-.31)+
  theme_bw()+
  theme(axis.text = element_text(size = 6),
        plot.title = element_text(hjust = .5, size = 6, vjust = -10),
        axis.title.y = element_text(size = 6),
        axis.title.x = element_text(size = 6),
        legend.position = 'none', aspect.ratio = 1,
        panel.border = element_rect(colour = "black", fill=NA, size=.5),
        panel.grid = element_blank(),
        axis.text.y = element_blank(),
        axis.ticks.y = element_blank())+
  labs(color = '',
       y = 'Heat flow (W/g)',
       title = 'AavLEA1',
       x = 'Temperature (\u00B0C)')+
  xlim(20,55)+
  
  scale_color_manual(values = c('black', '#CAE7D3','#9CDBC7','#63B89C','#448B74'))

#AtLEA3-3 Trehalose----
#read in data
AavLEA1T <- read_xlsx('Figure_5_DSC.xlsx', sheet = 'AtLEA3-3_Trehalose')
colnames(AavLEA1T)<- c('Temperature', 'Heat_Flow', 'Sample')
DSC_min <- AavLEA1T %>% 
  group_by(Sample)%>%
  summarise_at(vars(Heat_Flow),
               list(min = min))
#create df with discrete heat_flow values
Sample <- c('T0', 'T1', 'T10', 'T100', 'T500')
adj_heatflow <- c(-.324, -.33, -.339, -.338, -.352)
adjusted_heatflow <- data.frame(Sample, adj_heatflow)
#combine dataframe
Adjusted <- inner_join(DSC_min, adjusted_heatflow)
#find difference
Adjusted <- Adjusted %>% mutate(Diff= adj_heatflow - min)
#select and merge with DSC 
Adjusted <- Adjusted %>% select(c(Sample, Diff))
#merge
Adjusted_DSC <- inner_join(AavLEA1T, Adjusted)
Adjusted_DSC <- Adjusted_DSC %>% mutate(adjusted_heatflow = Heat_Flow + Diff)
#import data for enthalpy selection
DSC_AtLEA33T<-ggplot(data = Adjusted_DSC, aes(x = Temperature, y = adjusted_heatflow))+
  geom_line(aes(color = Sample))+
  ylim(-.35,-.31)+
  theme_bw()+
  theme(axis.text = element_text(size = 6),
        plot.title = element_text(hjust = .5, size = 6, vjust = -10),
        axis.title.y = element_text(size = 6),
        axis.title.x = element_text(size = 6),
        legend.position = 'none', aspect.ratio = 1,
        panel.border = element_rect(colour = "black", fill=NA, size=.5),
        panel.grid = element_blank(),
        axis.text.y = element_blank(),
        axis.ticks.y = element_blank())+
  labs(color = '',
       y = 'Heat flow (W/g)',
       title = 'AtLEA3-3',
       x = 'Temperature (\u00B0C)')+
  xlim(20,55)+
  
  scale_color_manual(values = c('black', '#B0E0E6','#8CB8DE','#6393BF','#36628B'))
#AtLEA3-3 Sucrose----
#read in data
AavLEA1S <- read_xlsx('Figure_5_DSC.xlsx', sheet = 'AtLEA3-3_Sucrose')
colnames(AavLEA1S)<- c('Temperature', 'Heat_Flow', 'Sample')
DSC_min <- AavLEA1S %>% 
  group_by(Sample)%>%
  summarise_at(vars(Heat_Flow),
               list(min = min))
#create df with discrete heat_flow values
Sample <- c('S0', 'S1', 'S10', 'S100', 'S500')
adj_heatflow <- c(-.324, -.33, -.336, -.342, -.468)
adjusted_heatflow <- data.frame(Sample, adj_heatflow)
#combine dataframe
Adjusted <- inner_join(DSC_min, adjusted_heatflow)
#find difference
Adjusted <- Adjusted %>% mutate(Diff= adj_heatflow - min)
#select and merge with DSC 
Adjusted <- Adjusted %>% select(c(Sample, Diff))
#merge
Adjusted_DSC <- inner_join(AavLEA1S, Adjusted)
Adjusted_DSC <- Adjusted_DSC %>% mutate(adjusted_heatflow = Heat_Flow + Diff)
#import data for enthalpy selection
DSC_AtLEA33S<-ggplot(data = Adjusted_DSC, aes(x = Temperature, y = adjusted_heatflow))+
  geom_line(aes(color = Sample))+
  ylim(-.35,-.31)+
  theme_bw()+
  theme(axis.text = element_text(size = 6),
        plot.title = element_text(hjust = .5, size = 6, vjust = -10),
        axis.title.y = element_text(size = 6),
        axis.title.x = element_text(size = 6),
        legend.position = 'none', aspect.ratio = 1,
        panel.border = element_rect(colour = "black", fill=NA, size=.5),
        panel.grid = element_blank(),
        axis.text.y = element_blank(),
        axis.ticks.y = element_blank())+
  labs(color = '',
       y = 'Heat flow (W/g)',
       title = 'AtLEA3-3',
       x = 'Temperature (\u00B0C)')+
  xlim(20,55)+
  
  scale_color_manual(values = c('black', '#CAE7D3','#9CDBC7','#63B89C','#448B74'))
#AtLEA4-2 Trehalose----
#read in data
AavLEA1T <- read_xlsx('Figure_5_DSC.xlsx', sheet = 'AtLEA4-2_Trehalose')
colnames(AavLEA1T)<- c('Temperature', 'Heat_Flow', 'Sample')
DSC_min <- AavLEA1T %>% 
  group_by(Sample)%>%
  summarise_at(vars(Heat_Flow),
               list(min = min))
#create df with discrete heat_flow values
Sample <- c('T0', 'T1', 'T10', 'T100', 'T500')
adj_heatflow <- c(-.324, -.336, -.336, -.342, -.348)
adjusted_heatflow <- data.frame(Sample, adj_heatflow)
#combine dataframe
Adjusted <- inner_join(DSC_min, adjusted_heatflow)
#find difference
Adjusted <- Adjusted %>% mutate(Diff= adj_heatflow - min)
#select and merge with DSC 
Adjusted <- Adjusted %>% select(c(Sample, Diff))
#merge
Adjusted_DSC <- inner_join(AavLEA1T, Adjusted)
Adjusted_DSC <- Adjusted_DSC %>% mutate(adjusted_heatflow = Heat_Flow + Diff)
#import data for enthalpy selection
DSC_AtLEA42T<-ggplot(data = Adjusted_DSC, aes(x = Temperature, y = adjusted_heatflow))+
  geom_line(aes(color = Sample))+
  ylim(-.35,-.31)+
  theme_bw()+
  theme(axis.text = element_text(size = 6),
        plot.title = element_text(hjust = .5, size = 6, vjust = -10),
        axis.title.y = element_text(size = 6),
        axis.title.x = element_text(size = 6),
        legend.position = 'none', aspect.ratio = 1,
        panel.border = element_rect(colour = "black", fill=NA, size=.5),
        panel.grid = element_blank(),
        axis.text.y = element_blank(),
        axis.ticks.y = element_blank())+
  labs(color = '',
       y = 'Heat flow (W/g)',
       title = 'AtLEA4-2',
       x = 'Temperature (\u00B0C)')+
  xlim(20,55)+
  
  scale_color_manual(values = c('black', '#B0E0E6','#8CB8DE','#6393BF','#36628B'))

#AtLEA4-2 Sucrose----
#read in data
AavLEA1S <- read_xlsx('Figure_5_DSC.xlsx', sheet = 'AtLEA4-2_Sucrose')
colnames(AavLEA1S)<- c('Temperature', 'Heat_Flow', 'Sample')
DSC_min <- AavLEA1S %>% 
  group_by(Sample)%>%
  summarise_at(vars(Heat_Flow),
               list(min = min))
#create df with discrete heat_flow values
Sample <- c('CS0', 'CS1', 'CS10', 'CS100', 'CS500')
adj_heatflow <- c(-.324, -.336, -.336, -.342, -.348)
adjusted_heatflow <- data.frame(Sample, adj_heatflow)
#combine dataframe
Adjusted <- inner_join(DSC_min, adjusted_heatflow)
#find difference
Adjusted <- Adjusted %>% mutate(Diff= adj_heatflow - min)
#select and merge with DSC 
Adjusted <- Adjusted %>% select(c(Sample, Diff))
#merge
Adjusted_DSC <- inner_join(AavLEA1S, Adjusted)
Adjusted_DSC <- Adjusted_DSC %>% mutate(adjusted_heatflow = Heat_Flow + Diff)
#import data for enthalpy selection
DSC_AtLEA42S<-ggplot(data = Adjusted_DSC, aes(x = Temperature, y = adjusted_heatflow))+
  geom_line(aes(color = Sample))+
  theme_bw()+
  theme(axis.text = element_text(size = 6),
        plot.title = element_text(hjust = .5, size = 6, vjust = -10),
        axis.title.y = element_text(size = 6),
        axis.title.x = element_text(size = 6),
        legend.position = 'none', aspect.ratio = 1,
        panel.border = element_rect(colour = "black", fill=NA, size=.5),
        panel.grid = element_blank(),
        axis.text.y = element_blank(),
        axis.ticks.y = element_blank())+
  labs(color = '',
       y = 'Heat flow (W/g)',
       title = 'AtLEA4-2',
       x = 'Temperature (\u00B0C)')+
  xlim(20,55)+
  ylim(-.35,-.31)+
  
  scale_color_manual(values = c('black', '#CAE7D3','#9CDBC7','#63B89C','#448B74'))
#AvLEA1C Trehalose----
#read in data
AavLEA1T <- read_xlsx('Figure_5_DSC.xlsx', sheet = 'AvLEA1C_Trehalose')
colnames(AavLEA1T)<- c('Temperature', 'Heat_Flow', 'Sample')
DSC_min <- AavLEA1T %>% 
  group_by(Sample)%>%
  summarise_at(vars(Heat_Flow),
               list(min = min))
#create df with discrete heat_flow values
Sample <- c('T0', 'T1', 'T10', 'T100', 'T500')
adj_heatflow <- c(-.324, -.336, -.336, -.342, -.352)
adjusted_heatflow <- data.frame(Sample, adj_heatflow)
#combine dataframe
Adjusted <- inner_join(DSC_min, adjusted_heatflow)
#find difference
Adjusted <- Adjusted %>% mutate(Diff= adj_heatflow - min)
#select and merge with DSC 
Adjusted <- Adjusted %>% select(c(Sample, Diff))
#merge
Adjusted_DSC <- inner_join(AavLEA1T, Adjusted)
Adjusted_DSC <- Adjusted_DSC %>% mutate(adjusted_heatflow = Heat_Flow + Diff)
#import data for enthalpy selection
DSC_AvLEA1CT<-ggplot(data = Adjusted_DSC, aes(x = Temperature, y = adjusted_heatflow))+
  geom_line(aes(color = Sample))+
  ylim(-.35,-.31)+
  theme_bw()+
  theme(axis.text = element_text(size = 6),
        plot.title = element_text(hjust = .5, size = 6, vjust = -10),
        axis.title.y = element_text(size = 6),
        axis.title.x = element_text(size = 6),
        legend.position = 'none', aspect.ratio = 1,
        panel.border = element_rect(colour = "black", fill=NA, size=.5),
        panel.grid = element_blank(),
        axis.text.y = element_blank(),
        axis.ticks.y = element_blank())+
  labs(color = '',
       y = 'Heat flow (W/g)',
       title = 'AvLEA1C',
       x = 'Temperature (\u00B0C)')+
  xlim(20,55)+
  
  scale_color_manual(values = c('black', '#B0E0E6','#8CB8DE','#6393BF','#36628B'))
#AvLEA1C Sucrose----
#read in data
AavLEA1S <- read_xlsx('Figure_5_DSC.xlsx', sheet = 'AvLEA1C_Sucrose')
colnames(AavLEA1S)<- c('Temperature', 'Heat_Flow', 'Sample')
DSC_min <- AavLEA1S %>% 
  group_by(Sample)%>%
  summarise_at(vars(Heat_Flow),
               list(min = min))
#create df with discrete heat_flow values
Sample <- c('S0', 'S1', 'S10', 'S100', 'S500')
adj_heatflow <- c(-.324, -.33, -.336, -.342, -.348)
adjusted_heatflow <- data.frame(Sample, adj_heatflow)
#combine dataframe
Adjusted <- inner_join(DSC_min, adjusted_heatflow)
#find difference
Adjusted <- Adjusted %>% mutate(Diff= adj_heatflow - min)
#select and merge with DSC 
Adjusted <- Adjusted %>% select(c(Sample, Diff))
#merge
Adjusted_DSC <- inner_join(AavLEA1S, Adjusted)
Adjusted_DSC <- Adjusted_DSC %>% mutate(adjusted_heatflow = Heat_Flow + Diff)
#import data for enthalpy selection
DSC_AvLEA1CS<-ggplot(data = Adjusted_DSC, aes(x = Temperature, y = adjusted_heatflow))+
  geom_line(aes(color = Sample))+
  theme_bw()+
  theme(axis.text = element_text(size = 6),
        plot.title = element_text(hjust = .5, size = 6, vjust = -10),
        axis.title.y = element_text(size = 6),
        axis.title.x = element_text(size = 6),
        legend.position = 'none', aspect.ratio = 1,
        panel.border = element_rect(colour = "black", fill=NA, size=.5),
        panel.grid = element_blank(),
        axis.text.y = element_blank(),
        axis.ticks.y = element_blank())+
  labs(color = '',
       y = 'Heat flow (W/g)',
       title = 'AvLEA1C',
       x = 'Temperature (\u00B0C)')+
  xlim(20,55)+
  ylim(-.35,-.31)+
  
  scale_color_manual(values = c('black', '#CAE7D3','#9CDBC7','#63B89C','#448B74'))
#BSA Trehalose----
#read in data
AavLEA1T <- read_xlsx('Figure_5_DSC.xlsx', sheet = 'BSA_Trehalose')
colnames(AavLEA1T)<- c('Temperature', 'Heat_Flow', 'Sample')
DSC_min <- AavLEA1T %>% 
  group_by(Sample)%>%
  summarise_at(vars(Heat_Flow),
               list(min = min))
#create df with discrete heat_flow values
Sample <- c('T0', 'T1', 'T10', 'T100', 'T500')
adj_heatflow <- c(-.324, -.33, -.336, -.342, -.348)
adjusted_heatflow <- data.frame(Sample, adj_heatflow)
#combine dataframe
Adjusted <- inner_join(DSC_min, adjusted_heatflow)
#find difference
Adjusted <- Adjusted %>% mutate(Diff= adj_heatflow - min)
#select and merge with DSC 
Adjusted <- Adjusted %>% select(c(Sample, Diff))
#merge
Adjusted_DSC <- inner_join(AavLEA1T, Adjusted)
Adjusted_DSC <- Adjusted_DSC %>% mutate(adjusted_heatflow = Heat_Flow + Diff)
#import data for enthalpy selection
DSC_BSAT<-ggplot(data = Adjusted_DSC, aes(x = Temperature, y = adjusted_heatflow))+
  geom_line(aes(color = Sample))+
  ylim(-.35,-.31)+
  theme_bw()+
  theme(axis.text = element_text(size = 6),
        plot.title = element_text(hjust = .5, size = 6, vjust = -10),
        axis.title.y = element_text(size = 6),
        axis.title.x = element_text(size = 6),
        legend.position = 'none', aspect.ratio = 1,
        panel.border = element_rect(colour = "black", fill=NA, size=.5),
        panel.grid = element_blank(),
        axis.text.y = element_blank(),
        axis.ticks.y = element_blank())+
  labs(color = '',
       y = 'Heat flow (W/g)',
       title = 'BSA',
       x = 'Temperature (\u00B0C)')+
  xlim(20,55)+
  
  scale_color_manual(values = c('black', '#B0E0E6','#8CB8DE','#6393BF','#36628B'))
#BSA Sucrose----
#read in data
AavLEA1S <- read_xlsx('Figure_5_DSC.xlsx', sheet = 'BSA_Sucrose')
colnames(AavLEA1S)<- c('Temperature', 'Heat_Flow', 'Sample')
DSC_min <- AavLEA1S %>% 
  group_by(Sample)%>%
  summarise_at(vars(Heat_Flow),
               list(min = min))
#create df with discrete heat_flow values
Sample <- c('S0', 'S1', 'S10', 'S100', 'S500')
adj_heatflow <- c(-.324, -.33, -.336, -.342, -.348)
adjusted_heatflow <- data.frame(Sample, adj_heatflow)
#combine dataframe
Adjusted <- inner_join(DSC_min, adjusted_heatflow)
#find difference
Adjusted <- Adjusted %>% mutate(Diff= adj_heatflow - min)
#select and merge with DSC 
Adjusted <- Adjusted %>% select(c(Sample, Diff))
#merge
Adjusted_DSC <- inner_join(AavLEA1S, Adjusted)
Adjusted_DSC <- Adjusted_DSC %>% mutate(adjusted_heatflow = Heat_Flow + Diff)
#import data for enthalpy selection
DSC_BSAS<-ggplot(data = Adjusted_DSC, aes(x = Temperature, y = adjusted_heatflow))+
  geom_line(aes(color = Sample))+
  theme_bw()+
  theme(axis.text = element_text(size = 6),
        plot.title = element_text(hjust = .5, size = 6, vjust = -10),
        axis.title.y = element_text(size = 6),
        axis.title.x = element_text(size = 6),
        legend.position = 'none', aspect.ratio = 1,
        panel.border = element_rect(colour = "black", fill=NA, size=.5),
        panel.grid = element_blank(),
        axis.text.y = element_blank(),
        axis.ticks.y = element_blank())+
  labs(color = '',
       y = 'Heat flow (W/g)',
       title = 'BSA',
       x = 'Temperature (\u00B0C)')+
  xlim(20,55)+
  ylim(-.35,-.31)+
  
  scale_color_manual(values = c('black', '#CAE7D3','#9CDBC7','#63B89C','#448B74'))

#HeLEA68614 Trehalose----
#read in data
AavLEA1T <- read_xlsx('Figure_5_DSC.xlsx', sheet = 'HeLEA68614_Trehalose')
colnames(AavLEA1T)<- c('Temperature', 'Heat_Flow', 'Sample')
DSC_min <- AavLEA1T %>% 
  group_by(Sample)%>%
  summarise_at(vars(Heat_Flow),
               list(min = min))
#create df with discrete heat_flow values
Sample <- c('T0', 'T1', 'T10', 'T100', 'T500')
adj_heatflow <- c(-.324, -.33, -.336, -.342, -.348)
adjusted_heatflow <- data.frame(Sample, adj_heatflow)
#combine dataframe
Adjusted <- inner_join(DSC_min, adjusted_heatflow)
#find difference
Adjusted <- Adjusted %>% mutate(Diff= adj_heatflow - min)
#select and merge with DSC 
Adjusted <- Adjusted %>% select(c(Sample, Diff))
#merge
Adjusted_DSC <- inner_join(AavLEA1T, Adjusted)
Adjusted_DSC <- Adjusted_DSC %>% mutate(adjusted_heatflow = Heat_Flow + Diff)
#import data for enthalpy selection
DSC_HET<-ggplot(data = Adjusted_DSC, aes(x = Temperature, y = adjusted_heatflow))+
  geom_line(aes(color = Sample))+
  ylim(-.35,-.31)+
  theme_bw()+
  theme(axis.text = element_text(size = 6),
        plot.title = element_text(hjust = .5, size = 6, vjust = -10),
        axis.title.y = element_text(size = 6),
        axis.title.x = element_text(size = 6),
        legend.position = 'none', aspect.ratio = 1,
        panel.border = element_rect(colour = "black", fill=NA, size=.5),
        panel.grid = element_blank(),
        axis.text.y = element_blank(),
        axis.ticks.y = element_blank())+
  labs(color = '',
       y = 'Heat flow (W/g)',
       title = 'HeLEA68614',
       x = 'Temperature (\u00B0C)')+
  xlim(20,55)+
  
  scale_color_manual(values = c('black', '#B0E0E6','#8CB8DE','#6393BF','#36628B'))

#HeLEA68614 Sucrose----
#read in data
AavLEA1S <- read_xlsx('Figure_5_DSC.xlsx', sheet = 'HeLEA68614_Sucrose')
colnames(AavLEA1S)<- c('Temperature', 'Heat_Flow', 'Sample')
DSC_min <- AavLEA1S %>% 
  group_by(Sample)%>%
  summarise_at(vars(Heat_Flow),
               list(min = min))
#create df with discrete heat_flow values
Sample <- c('S0', 'S1', 'S10', 'S100', 'S500')
adj_heatflow <- c(-.324, -.33, -.336, -.342, -.348)
adjusted_heatflow <- data.frame(Sample, adj_heatflow)
#combine dataframe
Adjusted <- inner_join(DSC_min, adjusted_heatflow)
#find difference
Adjusted <- Adjusted %>% mutate(Diff= adj_heatflow - min)
#select and merge with DSC 
Adjusted <- Adjusted %>% select(c(Sample, Diff))
#merge
Adjusted_DSC <- inner_join(AavLEA1S, Adjusted)
Adjusted_DSC <- Adjusted_DSC %>% mutate(adjusted_heatflow = Heat_Flow + Diff)
#import data for enthalpy selection
DSC_HES<-ggplot(data = Adjusted_DSC, aes(x = Temperature, y = adjusted_heatflow))+
  geom_line(aes(color = Sample))+
  theme_bw()+
  theme(axis.text = element_text(size = 6),
        plot.title = element_text(hjust = .5, size = 6, vjust = -10),
        axis.title.y = element_text(size = 6),
        axis.title.x = element_text(size = 6),
        legend.position = 'none', aspect.ratio = 1,
        panel.border = element_rect(colour = "black", fill=NA, size=.5),
        panel.grid = element_blank(),
        axis.text.y = element_blank(),
        axis.ticks.y = element_blank())+
  labs(color = '',
       y = 'Heat flow (W/g)',
       title = 'HeLEA68614',
       x = 'Temperature (\u00B0C)')+
  xlim(20,55)+
  ylim(-.35,-.31)+
  
  scale_color_manual(values = c('black', '#CAE7D3','#9CDBC7','#63B89C','#448B74'))

#Trehalose----
#read in data
AavLEA1T <- read_xlsx('Figure_5_DSC.xlsx', sheet = 'Trehalose')
colnames(AavLEA1T)<- c('Temperature', 'Heat_Flow', 'Sample')
DSC_min <- AavLEA1T %>% 
  group_by(Sample)%>%
  summarise_at(vars(Heat_Flow),
               list(min = min))
#create df with discrete heat_flow values
Sample <- c('T1', 'T10', 'T100', 'T500')
adj_heatflow <- c(-.324, -.33, -.336, -.342)
adjusted_heatflow <- data.frame(Sample, adj_heatflow)
#combine dataframe
Adjusted <- inner_join(DSC_min, adjusted_heatflow)
#find difference
Adjusted <- Adjusted %>% mutate(Diff= adj_heatflow - min)
#select and merge with DSC 
Adjusted <- Adjusted %>% select(c(Sample, Diff))
#merge
Adjusted_DSC <- inner_join(AavLEA1T, Adjusted)
Adjusted_DSC <- Adjusted_DSC %>% mutate(adjusted_heatflow = Heat_Flow + Diff)
#import data for enthalpy selection
DSC_Trehalose<-ggplot(data = Adjusted_DSC, aes(x = Temperature, y = adjusted_heatflow))+
  geom_line(aes(color = Sample))+
  ylim(-.35,-.31)+
  theme_bw()+
  theme(axis.text = element_text(size = 6),
        plot.title = element_text(hjust = .5, size = 6, vjust = -10),
        axis.title.y = element_text(size = 6),
        axis.title.x = element_text(size = 6),
        legend.position = 'none', aspect.ratio = 1,
        panel.border = element_rect(colour = "black", fill=NA, size=.5),
        panel.grid = element_blank(),
        axis.text.y = element_blank(),
        axis.ticks.y = element_blank())+
  labs(color = '',
       y = 'Heat flow (W/g)',
       title = 'Trehalose',
       x = 'Temperature (\u00B0C)')+
  xlim(20,55)+
  
  scale_color_manual(values = c('#B0E0E6','#8CB8DE','#6393BF','#36628B'))

#Sucrose----
#read in data
AavLEA1S <- read_xlsx('Figure_5_DSC.xlsx', sheet = 'Sucrose')
colnames(AavLEA1S)<- c('Temperature', 'Heat_Flow', 'Sample')
DSC_min <- AavLEA1S %>% 
  group_by(Sample)%>%
  summarise_at(vars(Heat_Flow),
               list(min = min))
#create df with discrete heat_flow values
Sample <- c('S1', 'S10', 'S100', 'S500')
adj_heatflow <- c(-.324, -.333, -.336, -.342)
adjusted_heatflow <- data.frame(Sample, adj_heatflow)
#combine dataframe
Adjusted <- inner_join(DSC_min, adjusted_heatflow)
#find difference
Adjusted <- Adjusted %>% mutate(Diff= adj_heatflow - min)
#select and merge with DSC 
Adjusted <- Adjusted %>% select(c(Sample, Diff))
#merge
Adjusted_DSC <- inner_join(AavLEA1S, Adjusted)
Adjusted_DSC <- Adjusted_DSC %>% mutate(adjusted_heatflow = Heat_Flow + Diff)
#import data for enthalpy selection
DSC_Sucrose<-ggplot(data = Adjusted_DSC, aes(x = Temperature, y = adjusted_heatflow))+
  geom_line(aes(color = Sample))+
  theme_bw()+
  theme(axis.text = element_text(size = 6),
        plot.title = element_text(hjust = .5, size = 6, vjust = -10),
        axis.title.y = element_text(size = 6),
        axis.title.x = element_text(size = 6),
        legend.position = 'none', aspect.ratio = 1,
        panel.border = element_rect(colour = "black", fill=NA, size=.5),
        panel.grid = element_blank(),
        axis.text.y = element_blank(),
        axis.ticks.y = element_blank())+
  labs(color = '',
       y = 'Heat flow (W/g)',
       title = 'Sucrose',
       x = 'Temperature (\u00B0C)')+
  xlim(20,55)+
  ylim(-.35,-.31)+
  
  scale_color_manual(values = c('#CAE7D3','#9CDBC7','#63B89C','#448B74'))

DSC_T6 + 
DSC_S6 +
SE+
TE+
DSC_AavLEA1T+
DSC_AavLEA1S+
DSC_AtLEA33T+
  DSC_AtLEA33S+
DSC_AtLEA42T+
  DSC_AtLEA42S+
  DSC_AvLEA1CT+
  DSC_AvLEA1CS+
  DSC_BSAT+
  DSC_BSAS+
  DSC_HET+
  DSC_HES+
  DSC_Trehalose+
  DSC_Sucrose+
plot_layout(ncol = 4)
#Export 7x8
```
