## Supplementary material for "Disordered proteins interact with the chemical environment to tune their protective function during drying": File S3: Figure_6.R

```
setwd('~/Documents/Research_Projects/LEA_CAHS_Oligomerization/Submission')

#call in packages
library(tidyverse)
library(dplyr)
library(purrr)
library(broom)
library(stringr)
library(ggplot2)
library(plotly)
library(readxl)
library(RColorBrewer)
library(patchwork)
library(grid)

Figure_6B <- read_xlsx('Figure_6B_TFE Barplot Data.xlsx')
Figure_6B <- Figure_5B %>% filter(Protein == 'CAHS D')
Figure_6B$Cosolute<-factor(Figure_6B$Cosolute, levels = c('Trehalose', 'Sucrose', 'Betaine'))
Fig6B <-ggplot(data = Figure_6B, aes(x = Cosolute, y = -`dG Dimerization`, fill = Cosolute))+
  geom_bar(position = position_dodge(), stat = 'identity', color = 'black', size = .3)+
  theme_bw()+
  theme(axis.text = element_text(size = 6),
        plot.title = element_text(hjust = 0.95, size = 6, vjust = -10),
        axis.title.y = element_text(size = 6),
        axis.title.x = element_text(size = 6),
        legend.position = 'none', aspect.ratio = 1,
        panel.border = element_rect(colour = "black", fill=NA, size=.5))+
  labs(title = '',
       x = 'Osmolyte',
       y = expression(t*Delta*Delta*"G"^dimer*" "*(cal/moll/M)))+
  scale_fill_manual(values = c('#36628B', '#448B74','#a63a2e'))+
  geom_errorbar(aes(ymin = -`dG Dimerization` - `dG Dimerization Error`, ymax = -`dG Dimerization` + `dG Dimerization Error`), width = .2, position = position_dodge(.9), size = .3)+
  ylim(-5000,2500)
#Figure 5C
#read in data
B6 <- read_xlsx('Figure_6C_Betaine_DSC.xlsx', sheet = 'Betaine')
colnames(B6)<- c('Temperature', 'Heat_Flow', 'Sample')
DSC_min <- B6 %>% 
  group_by(Sample)%>%
  summarise_at(vars(Heat_Flow),
               list(min = min))
#create df with discrete heat_flow values
Sample <- c('CB0', 'CB1', 'CB10', 'CB100', 'CB500')
adj_heatflow <- c(-.324, -.33, -.336, -.342, -.348)
adjusted_heatflow <- data.frame(Sample, adj_heatflow)
#combine dataframe
Adjusted <- inner_join(DSC_min, adjusted_heatflow)
#find difference
Adjusted <- Adjusted %>% mutate(Diff= adj_heatflow - min)
#select and merge with DSC 
Adjusted <- Adjusted %>% select(c(Sample, Diff))
#merge
Adjusted_DSC <- inner_join(B6, Adjusted)
Adjusted_DSC <- Adjusted_DSC %>% mutate(adjusted_heatflow = Heat_Flow + Diff)
Betaine_markers <- read_xlsx('Figure_6C_Betaine_DSC.xlsx', sheet = 'Betaine_points')
#Figure
Fig6C<-ggplot(data = Adjusted_DSC, aes(x = Temperature, y = adjusted_heatflow))+
  geom_line(aes(color = Sample))+
  geom_point(data = Betaine_markers, aes(x = Temperature, y = adjusted_heatflow), shape = 3, size = 1)+
  ylim(-.35,-.31)+
  theme_bw()+
  theme(axis.text = element_text(size = 6),
        plot.title = element_text(hjust = 0.95, size = 6, vjust = -10),
        axis.title.y = element_text(size = 6),
        axis.title.x = element_text(size = 6),
        legend.position = 'none', aspect.ratio = 1,
        panel.border = element_rect(colour = "black", fill=NA, size=.5),
        panel.grid = element_blank(),,
        axis.text.y = element_blank(),
        axis.ticks.y = element_blank())+
  labs(color = '',
       y = 'Heat flow (W/g)',
       title = '6 mg/mL',
       x = 'Temperature (\u00B0C)')+
  xlim(20,55)+
  
  scale_color_manual(values = c( 'black', '#cb7c6e','#bf6758','#b35142','#a63a2e'))
#Figure 6D
df <- read_xlsx('Figure_6D_Betaine_6mg_Enthalpy.xlsx', sheet = 'CAHSD_Betaine_6')
#find mean of different trials
df <- df %>% filter(Sample %in% c('CB1', 'CB10', 'CB100', 'CB500'))
Fig6D <-ggplot(data = df, aes(x = Sample, y = Avg_C_Enthalpy*1000, fill = Sample))+
  geom_bar(position = position_dodge(), stat = 'identity', color = 'black', size = .3)+
  theme_bw()+
  theme(axis.text = element_text(size = 6),
        plot.title = element_text(hjust = 0.95, size = 6, vjust = -10),
        axis.title.y = element_text(size = 6),
        axis.title.x = element_text(size = 6),
        legend.position = 'none', aspect.ratio = 1,
        panel.border = element_rect(colour = "black", fill=NA, size=.5))+
  labs(title = '6 mg/mL',
       x = 'Molar Ratio (Betaine:CAHS D)',
       y = expression(Delta*'Enthalpy (mJ/g)'))+
  scale_fill_manual(values = c( '#cb7c6e','#bf6758','#b35142','#a63a2e'))+
  scale_x_discrete(breaks=c("CB1","CB10", "CB100", "CB500"),
                   labels=c("1:1","10:1", "100:1", "500:1"))+
  geom_errorbar(aes(ymin = Avg_C_Enthalpy*1000 - Stdev_C_Enthalpy*1000, ymax = Avg_C_Enthalpy*1000 + Stdev_C_Enthalpy*1000), width = .2, position = position_dodge(.9), size = .3)+
  ylim(-2.5,5)

#12mg Betaine Enthalpy----
B12_E <- read_xlsx('Figure_6F_Betaine_12_Enthalpy.xlsx', sheet = 'CAHSD_Betaine_12')
B12_E <- B12_E %>% filter(Sample %in% c('CB1', 'CB10', 'CB100', 'CB500'))
Fig6F<-ggplot(data = B12_E, aes(x = Sample, y = Avg_C_Enthalpy*1000, fill = Sample))+
  geom_bar(position = position_dodge(), stat = 'identity', color = 'black', size = .3)+
  theme_bw()+
  theme(axis.text = element_text(size = 6),
        plot.title = element_text(hjust = 0.95, size = 6, vjust = -10),
        axis.title.y = element_text(size = 6),
        axis.title.x = element_text(size = 6),
        legend.position = 'none', aspect.ratio = 1,
        panel.border = element_rect(colour = "black", fill=NA, size=.5))+
  labs(
    x = 'Molar Ratio (Betaine:CAHS D)',
    y = expression(Delta*'Enthalpy (mJ/g)'),
    title = '12 mg/mL')+
  scale_fill_manual(values = c( '#cb7c6e','#bf6758','#bf6758','#a63a2e'))+
  scale_x_discrete(breaks=c("CB1","CB10", "CB100", "CB500"),
                   labels=c("1:1","10:1", "100:1", "500:1"))+
  geom_errorbar(aes(ymin = Avg_C_Enthalpy*1000 - Stdev_C_Enthalpy*1000, ymax = Avg_C_Enthalpy*1000 + Stdev_C_Enthalpy*1000), width = .2, position = position_dodge(.9), size = .3)+
  ylim(-25,25)

#12Betaine----
#read in data
B12 <- read_xlsx('Figure_6E_Betaine_DSC.xlsx', sheet = 'Betaine')
colnames(B12)<- c('Temperature', 'Heat_Flow', 'Sample')
DSC_min <- B12 %>% 
  group_by(Sample)%>%
  summarise_at(vars(Heat_Flow),
               list(min = min))
#create df with discrete heat_flow values
Sample <- c('CB0', 'CB1', 'CB10', 'CB100', 'CB500')
adj_heatflow <- c(-.324, -.33, -.336, -.342, -.348)
adjusted_heatflow <- data.frame(Sample, adj_heatflow)
#combine dataframe
Adjusted <- inner_join(DSC_min, adjusted_heatflow)
#find difference
Adjusted <- Adjusted %>% mutate(Diff= adj_heatflow - min)
#select and merge with DSC 
Adjusted <- Adjusted %>% select(c(Sample, Diff))
#merge
Adjusted_DSC <- inner_join(B12, Adjusted)
Adjusted_DSC <- Adjusted_DSC %>% mutate(adjusted_heatflow = Heat_Flow + Diff)

Betaine_12Markers <- read_xlsx('Figure_6E_Betaine_DSC.xlsx', sheet = 'Betaine_points')

Fig6E<-ggplot(data = Adjusted_DSC, aes(x = Temperature, y = adjusted_heatflow))+
  geom_line(aes(color = Sample))+
  geom_point(data = Betaine_12Markers, aes(x = Temperature, y = adjusted_heatflow), shape = 3, size = 1)+
  theme_bw()+
  theme(axis.text = element_text(size = 6),
        plot.title = element_text(hjust = 0.95, size = 6, vjust = -10),
        axis.title.y = element_text(size = 6),
        axis.title.x = element_text(size = 6),
        legend.position = 'none', aspect.ratio = 1,
        panel.border = element_rect(colour = "black", fill=NA, size=.5),
        panel.grid = element_blank(),
        axis.text.y = element_blank(),
        axis.ticks.y = element_blank())+
  labs(color = '',
       y = 'Heat flow (W/g)',
       title = '12 mg/mL',
       x = 'Temperature (\u00B0C)')+
  xlim(20,55)+
  ylim(-.35,-.31)+
  
  scale_color_manual(values = c( 'black', '#cb7c6e','#bf6758','#b35142','#a63a2e'))+
  annotation_custom(segmentsGrob(c(0.3, -0.1), c(-0.085, 0.28), 
                                 c(1, -0.1), c(-0.085, 1), gp = gpar(lwd = 2),
                                 arrow = arrow(length = unit(2.5, 'mm'))))
#Betaine SAXS
setwd('~/Documents/Research_Projects/LEA_CAHS_Oligomerization/Submission/Figure_6G')
library (tidyverse)
library (patchwork)
library(readxl)


col_list <- c("black", "#36628B", "#448B74", "#A63A2E")

Tris <- read_csv("Tris.csv")
Trehalose <- read_csv("Trehalose.csv")
Sucrose <- read_csv("Sucrose.csv")
Betaine <- read_csv("Betaine.csv")

Fig6G <- ggplot() +
  geom_line(data = Tris, aes(x = (`Q`), y = log(`I(Q)`)), color = col_list[1], size = .75) +
  geom_line(data = Trehalose, aes(x = (`Q`), y = log(`I(Q)`)), color = col_list[2], size = .75) +
  geom_line(data = Sucrose, aes(x = (`Q`), y = log(`I(Q)`)), color = col_list[3], size = .75) +
  geom_line(data = Betaine, aes(x = (`Q`), y = log(`I(Q)`)), color = col_list[4], size = .75) +
  theme_bw()+
  theme(
    plot.title = element_text(hjust = 0.1, size = 10, vjust = -11),
    axis.title = element_text(size = 6),
    axis.text.y = element_text(size = 6),
    axis.text.x = element_text(size = 6),
    legend.text=element_text(size=14),
    legend.position = 'none', aspect.ratio = 1,
    legend.title = element_blank(),
    panel.border = element_rect(colour = "black", fill=NA, size=.5)) +
  labs(x = expression('q' * ' ' * (Å ^ -1)), y = 'ln(I(q))', title = '') +
  xlim(0.015, 0.1) +
  ylim (0, 5)
#Betaine Synergy

#Import necessary packages
library(ggplot2)
library(dplyr)
library(readxl)
library(ggsignif)
library(patchwork)
setwd('~/Documents/Research_Projects/LEA_CAHS_Oligomerization/Submission')
#Import your dataset
data <- read_xlsx('Figure_6H_CAHSD_Betaine_Synergy.xlsx', sheet = 'R')
data$Ratio <- as.factor(data$Ratio)
data <- data %>% filter(Ratio == '1' | Ratio == '10'| Ratio == '100')

#####make plot
#create ggplot Environment
Fig6H<-ggplot(data = data, aes(x = Ratio, y = Protect_add, fill = interaction(Ratio, Calculation)))+
  #Make stacked bar graph
  ##Make bottom layer first 
  geom_col(data = data %>% filter(Protectant == 'Sugar'| Protectant == 'Mixed'), 
           position = position_dodge(width = .9), 
           color = 'black', size = .2)+
  ##Make top layer w/ alpha adjustment
  geom_col(data = data %>% filter(Protectant == 'Protein'| Protectant == 'Mixed'), 
           position = position_dodge(width = .9), 
           alpha = .6, 
           color = 'black', size = .2)+
  #Set y-axis scale
  scale_y_continuous(name = '%Protection',
                     breaks = seq(0,100,20),
                     limits = c(0,100))+  
  #Set graph labels
  labs(x = 'Molar Ratio (Betaine:CAHS D)',
       y = 'LDH Protection (%)')+
  #Make Black and white theme
  theme_bw()+
  #Adjust graph settings
  theme(
    plot.title = element_text(hjust = 0.95, size = 6, vjust = -10),
    axis.title = element_text(size =6),
    axis.text.y = element_text(size =6),
    axis.text.x = element_text(size =6),
    legend.text=element_text(size=12), 
    legend.position = 'none', aspect.ratio = 1,
    panel.border = element_rect(colour = "black", fill=NA, size=.5))+
  #add error bars based on standard deviation
  geom_errorbar(aes(ymin = Protect_add - Stdev, ymax = Protect_add + Stdev), width = .3, position = position_dodge(.9))+
  #custom assignment of bar color
  scale_fill_manual(values = c('lavenderblush1','lavenderblush1', 'lavenderblush1', 
                               '#A63A2E', '#A63A2E', '#A63A2E'))+
  #add significane values
  geom_signif(
    y_position = c(45, 43, 43),
    xmin = c(.75, 1.75, 2.75),
    xmax = c(1.25, 2.25, 3.25),
    annotation = c("NS", "NS", "***"),
    tip_length = 0, 
    color = "black",
    textsize = 2)+
  
  #add custom x labels
  scale_x_discrete(labels = c('1' = '1:1',
                              '10' = '10:1',
                              '100' = '100:1'))

Fig6B+Fig6B + Fig6C  +Fig6D  + Fig6E + Fig6F+ Fig6G + Fig6H + Fig6E +plot_layout(ncol = 3) + plot_annotation(tag_levels = list(c('A)','B)','C)','D)','E)','F)','G)','H)','I)'))) & theme(plot.tag = element_text(size = 8))
#Export 6x7
```
