## Supplementary material for "Disordered proteins interact with the chemical environment to tune their protective function during drying": File S3: G_Dimerization Correlation Plot.R

```
library(tidyverse)
library(readxl)
library(ggpmisc)
setwd("C:/Users/Vincent/OneDrive/Documents/Boothby Lab Work/Team CAHSD/Oligomerization Project")

prot_list <- c("BSA", "AtLEA3-3", "AavLEA", "HeLEA", "AvLEA", "AtLEA4-2", "CAHS D")
cor_list <- c()
r2_list <- c()

for(i in 1:length(prot_list))
{
  protein = prot_list[i]
  
  file <- read_excel("Synergy Correlation Values.xlsx")
  file <- subset(file, Protein == protein)
  
  if(protein == "CAHS D")
  {
    cols <- c("#A63A2E", "#448B74", "#36628B")
  }else{
    cols <- c("#448B74", "#36628B")
  }
  
  file$`TFE` = file$`Adjusted dG Dimerization`
  file$`Metric` = file$`Synergy`
  
  plot <- ggplot(data = file) +
    geom_point(aes(x = `Metric`, y = `TFE`, fill = `Cosolute`), pch = 21, size = 4) +
    geom_smooth(method = 'lm', aes(x = `Metric`, y = `TFE`), se = F, color = "black", size = 1) +
    theme_bw()+
    theme(
      plot.title = element_text(hjust = 0.05, size = 14, vjust = -8),
      axis.title = element_text(size =14),
      axis.text.y = element_text(size =10),
      axis.text.x = element_text(size = 10, angle = 70, vjust = 1, hjust=1),
      legend.text=element_text(size=14),
      legend.position = 'none', aspect.ratio = 1,
      legend.title = element_blank(),
      panel.border = element_rect(colour = "black", fill=NA, size=1)) +
    scale_fill_manual(values = cols) +
    labs(x = 'Placeholder', y = 'Placeholder (Edit in Illustrator)', title = protein) +
    ylim(min(file$`TFE`) - 1, max(file$`TFE`) + 5) +
    xlim(min(file$`Metric` - 0.2), max(file$`Metric`) + 0.2)
  
  cor_list[i] <- cor.test(file$`TFE`, file$`Metric`, method = "pearson")$p.value
  reg_1 <- lm(file$`TFE` ~ file$`Metric`)
  r2_list[i] <- summary(reg_1)$r.squared
  
  print(plot)
}
```
