## Supplementary material for "Disordered proteins interact with the chemical environment to tune their protective function during drying": File S3: SAXS Curve Comparison Zoom.R

```
library (tidyverse)
library (patchwork)
library(readxl)

setwd("C:/Users/Vincent/OneDrive/Documents/csv SAXS Dataset/500")

col_list <- c("#acacac", "#36628B", "#448B74", "#A63A2E")

Tris <- read_csv("Tris.csv")
Trehalose <- read_csv("Trehalose.csv")
Sucrose <- read_csv("Sucrose.csv")
Betaine <- read_csv("Betaine.csv")

Tris_adjusted = Tris
  Tris_adjusted$`I(Q)` = Tris_adjusted$`I(Q)` - Tris$`I(Q)`
Trehalose_adjusted = Trehalose
  Trehalose_adjusted$`I(Q)` = Trehalose_adjusted$`I(Q)` - Tris$`I(Q)`
Sucrose_adjusted = Sucrose
 Sucrose_adjusted$`I(Q)` = Sucrose_adjusted$`I(Q)` - Tris$`I(Q)`
Betaine_adjusted = Betaine
  Betaine_adjusted$`I(Q)` = Betaine_adjusted$`I(Q)` - Tris$`I(Q)`

Full <- ggplot() +
  geom_line(data = Tris, aes(x = (`Q`), y = log(`I(Q)`)), color = col_list[1], size = 1) +
  geom_line(data = Trehalose, aes(x = (`Q`), y = log(`I(Q)`)), color = col_list[2], size = 1) +
  geom_line(data = Sucrose, aes(x = (`Q`), y = log(`I(Q)`)), color = col_list[3], size = 1) +
  geom_line(data = Betaine, aes(x = (`Q`), y = log(`I(Q)`)), color = col_list[4], size = 1) +
  theme_bw()+
  theme(
    plot.title = element_text(hjust = 0.1, size = 10, vjust = -11),
    axis.title = element_text(size = 12),
    axis.text.y = element_text(size = 12),
    axis.text.x = element_text(size = 12),
    legend.text=element_text(size=14),
    legend.position = 'none', aspect.ratio = 1,
    legend.title = element_blank(),
    panel.border = element_rect(colour = "black", fill=NA, size=1)) +
  labs(x = expression('q' * ' ' * (Å ^ -1)), y = 'ln(I(q))', title = '') +
  xlim(0, 0.1) +
  ylim (0, 5)

Full

Full <- ggplot() +
  geom_line(data = Tris, aes(x = (`Q`), y = log(`I(Q)`)), color = col_list[1], size = 1) +
  geom_line(data = Trehalose, aes(x = (`Q`), y = log(`I(Q)`)), color = col_list[2], size = 1) +
  geom_line(data = Sucrose, aes(x = (`Q`), y = log(`I(Q)`)), color = col_list[3], size = 1) +
  theme_bw()+
  theme(
    plot.title = element_text(hjust = 0.1, size = 10, vjust = -11),
    axis.title = element_text(size = 12),
    axis.text.y = element_text(size = 12),
    axis.text.x = element_text(size = 12),
    legend.text=element_text(size=14),
    legend.position = 'none', aspect.ratio = 1,
    legend.title = element_blank(),
    panel.border = element_rect(colour = "black", fill=NA, size=1)) +
  labs(x = expression('q' * ' ' * (Å ^ -1)), y = 'ln(I(q))', title = '') +
  xlim(0.015, 0.41) +
  ylim (-2.5, 5)

Full

Full <- ggplot() +
  geom_line(data = Trehalose_adjusted, aes(x = (`Q`), y = `I(Q)`), color = col_list[2], size = 1) +
  geom_line(data = Sucrose_adjusted, aes(x = (`Q`), y = `I(Q)`), color = col_list[3], size = 1) +
  geom_line(data = Betaine_adjusted, aes(x = (`Q`), y = `I(Q)`), color = col_list[4], size = 1) +
  theme_bw()+
  theme(
    plot.title = element_text(hjust = 0.1, size = 10, vjust = -11),
    axis.title = element_text(size = 12),
    axis.text.y = element_text(size = 12),
    axis.text.x = element_text(size = 12),
    legend.text=element_text(size=14),
    legend.position = 'none', aspect.ratio = 1,
    legend.title = element_blank(),
    panel.border = element_rect(colour = "black", fill=NA, size=1)) +
  labs(x = expression('q' * ' ' * (Å ^ -1)), y = 'I(q)', title = '') +
  xlim(0.015, 0.41) +
  ylim (-5, 50)

Full

Guinier <- ggplot() +
  geom_line(data = Tris, aes(x = (`Q` * `Q`), y = log(`I(Q)`)), color = col_list[1], size = 1) +
  geom_line(data = Trehalose, aes(x = (`Q` * `Q`), y = log(`I(Q)`)), color = col_list[2], size = 1) +
  geom_line(data = Sucrose, aes(x = (`Q` * `Q`), y = log(`I(Q)`)), color = col_list[3], size = 1) +
  geom_line(data = Betaine, aes(x = (`Q` * `Q`), y = log(`I(Q)`)), color = col_list[4], size = 1) +
  theme_bw()+
  theme(
    plot.title = element_text(hjust = 0.1, size = 10, vjust = -11),
    axis.title = element_text(size = 12),
    axis.text.y = element_text(size = 12),
    axis.text.x = element_text(size = 12),
    legend.text=element_text(size=14),
    legend.position = 'none', aspect.ratio = 1,
    legend.title = element_blank(),
    panel.border = element_rect(colour = "black", fill=NA, size=1)) +
  labs(x = expression('q' * ' ' * (Å ^ -1)), y = 'ln(I(q))', title = '') +
  xlim(0, 0.0015) +
  ylim (0, 5)

Guinier
```
