## Supplementary material for "Disordered proteins interact with the chemical environment to tune their protective function during drying": File S3: TFE Barplot.R

```
library(tidyverse)
library(readxl)
library(ggpmisc)

names <- c("Trehalose", "Sucrose", "Betaine")
prot_list <- c("AtLEA3-3", "AavLEA", "HeLEA", "AvLEA", "AtLEA4-2", "CAHS D")

setwd("C:/Users/Vincent/OneDrive/Documents/Boothby Lab Work/Team CAHSD/Oligomerization Project")
file <- read_excel("TFE Barplot Data.xlsx")

for(i in 1:length(prot_list))
{
  if(prot_list[i] == "CAHS D")
  {
    cols <- c("#36628B", "#448B74", "#A63A2E")
  }
  else
  {
    cols <- c("#36628B", "#448B74")
  }
  
  temp <- subset(file, Protein == prot_list[i])
  
  plot <- ggplot(data = temp) +
    geom_bar(position = position_dodge(), stat = 'identity', color = 'black', aes(x = ordered(Cosolute, levels = names), y = `dG Dimerization`), fill = cols) +
    theme_bw()+
    theme(
      plot.title = element_text(hjust = 0.05, size = 14, vjust = -8),
      axis.title = element_text(size =14),
      axis.text.y = element_text(size =10),
      axis.text.x = element_text(size = 10),
      legend.text=element_text(size=14),
      legend.position = 'none', aspect.ratio = 1,
      legend.title = element_blank(),
      panel.border = element_rect(colour = "black", fill=NA, size=1)) +
    geom_errorbar(aes(x = `Cosolute`, ymin = `dG Dimerization` - `dG Dimerization Error`, ymax = `dG Dimerization` + `dG Dimerization Error`), width = 0.2, size = 0.8) +
    scale_fill_manual(values = cols) +
    labs(x = '', y = 'Placeholder (Edit in Illustrator)', title = paste("Dimerization TFE - ", prot_list[i])) +
    ylim(NA, max(temp$`dG Dimerization`) + 1500)
  
  print(plot)
}
```
