## Supplementary material for "Disordered proteins interact with the chemical environment to tune their protective function during drying": File S3: Fig. S3F.R

```
#Set working directory----
setwd('~/Downloads/')

#Import necessary packages----
library(ggplot2)
library(dplyr)
library(readxl)
library(ggsignif)

#Import your dataset----
data <- read_xlsx('Kapp.xlsx')
#Change the factor order
data$prot <- factor(data$prot,levels = c('AtLEA3-3',
                                         'AavLEA1',
                                         'HeLEA68614',
                                         'AvLEA1C',
                                         'AtLEA4-2',
                                         'CAHS D',
                                         'BSA'))
#build ggplot environment
ggplot(data = data, aes(x = `Kapp_tre`, y = `Kapp_suc`, fill = prot))+
  geom_point(size = 6.5, shape = 21, color = 'black')+
  theme_bw()+
  theme(panel.grid = element_blank(),
        plot.title = element_text(hjust = 0.05, size = 14, vjust = -8),
        axis.title = element_text(size =14),
        axis.text.y = element_text(size =12),
        axis.text.x = element_text(size =12),
        legend.text=element_text(size=14), 
        legend.position = 'none', aspect.ratio = 1,
        panel.border = element_rect(colour = "black", fill=NA, size=1))+
  xlim(-0.5,6)+
  ylim(-0.5,6)+
  scale_color_manual(values = c(
    'skyblue3', 'seagreen3', 'mediumpurple3', 'yellow3', 'coral3', 'azure4', 'thistle3'))+
  geom_abline(linetype = 'dashed',
              linewidth = 1)+
  geom_errorbar(aes(ymin = `Kapp_suc` - `Kapp_suc_err`,
                    ymax = `Kapp_suc` + `Kapp_suc_err`),
                width = .075, position = position_dodge(.9))+
  geom_errorbarh(aes(xmin = `Kapp_tre` - `Kapp_tre_err`,
                     xmax = `Kapp_tre` + `Kapp_tre_err`),
                 height = .075, position = position_dodge(.9))+
  labs(
    x = 'Synergy with Trehalose',
    y = 'Synergy with Sucrose'
  )
```
