## Supplementary material for "Disordered proteins interact with the chemical environment to tune their protective function during drying": File S3: S4.R

```
#Hydrated----
#Set working directory---
setwd('~/Documents/Research_Projects/LEA_CAHS_Oligomerization/')

#Import necessary packages----
library(ggplot2)
library(dplyr)
library(readxl)
library(ggsignif)
library(patchwork)

#Import your dataset----
data <- read_xlsx('Figure_2_CD_Compare.xlsx', sheet = 'Hydrated')
#Change the factor order
data$prot <- factor(data$prot,levels = c('AtLEA3-3',
                                         'AavLEA1',
                                         'HeLEA68614',
                                         'AvLEA1C',
                                         'AtLEA4-2',
                                         'BSA',
                                         'CAHS_D'))
#build ggplot environment
Hydrated <- ggplot(data = data, aes(x = `tre`, y = `suc`, fill = prot))+
  geom_point(size = 5, shape = 21, color = 'black')+
  theme_bw()+
  theme(panel.grid = element_blank(),
        plot.title = element_text(hjust = 0.05, size = 14, vjust = -8),
        axis.title = element_text(size =14),
        axis.text.y = element_text(size =14),
        axis.text.x = element_text(size =14),
        legend.text=element_text(size=14), 
        legend.position = 'none', aspect.ratio = 1,
        panel.border = element_rect(colour = "black", fill=NA, size=1))+
  scale_y_continuous(name = '222/210 Sucrose',
                     breaks = seq(.5,1,.2),
                     limits = c(.45,1))+  
  scale_x_continuous(name = '222/210 Trehalose',
                     breaks = seq(.5,1,.2),
                     limits = c(.45,1))+  
  
  scale_fill_manual(values = c(
    'skyblue3', 'seagreen3', 'mediumpurple3', 'yellow3', 'coral3', 'thistle3', 'blue'))+
  geom_abline(linetype = 'dashed',
              linewidth = 1)+
  geom_errorbar(aes(ymin = `suc` - `suc_err`,
                    ymax = `suc` + `suc_err`),
                width = .005, position = position_dodge(.9))+
  geom_errorbarh(aes(xmin = `tre` - `tre_err`,
                     xmax = `tre` + `tre_err`),
                 height = .005, position = position_dodge(.9))+
  labs(
    x = '%Synergy with Trehalose',
    y = '%Synergy with Sucrose',
    title = 'Hydrated'
  )

#Desiccated----
#Set working directory---
setwd('~/Documents/Research_Projects/LEA_CAHS_Oligomerization/')

#Import necessary packages----
library(ggplot2)
library(dplyr)
library(readxl)
library(ggsignif)

#Import your dataset----
data <- read_xlsx('Figure_2_CD_Compare.xlsx', sheet = 'Dry')
#Change the factor order
data$prot <- factor(data$prot,levels = c('AtLEA3-3',
                                         'AavLEA1',
                                         'HeLEA68614',
                                         'AvLEA1C',
                                         'AtLEA4-2',
                                         'BSA',
                                         'CAHS_D'))
#build ggplot environment
Dry <- ggplot(data = data, aes(x = `tre`, y = `suc`, fill = prot))+
  geom_point(size = 5, shape = 21, color = 'black')+
  theme_bw()+
  theme(panel.grid = element_blank(),
        plot.title = element_text(hjust = 0.05, size = 14, vjust = -8),
        axis.title = element_text(size =14),
        axis.text.y = element_text(size =14),
        axis.text.x = element_text(size =14),
        legend.text=element_text(size=14), 
        legend.position = 'none', aspect.ratio = 1,
        panel.border = element_rect(colour = "black", fill=NA, size=1))+
  scale_y_continuous(name = '222/210 Sucrose',
                     breaks = seq(.8,2.2,.4),
                     limits = c(.8,2.2))+  
  scale_x_continuous(name = '222/210 Trehalose',
                     breaks = seq(.8,2.2,.4),
                     limits = c(.8,2.2))+  
  
  scale_fill_manual(values = c(
    'skyblue3', 'seagreen3', 'mediumpurple3', 'yellow3', 'coral3', 'thistle3', 'blue'))+
  geom_abline(linetype = 'dashed',
              linewidth = 1)+
  geom_errorbar(aes(ymin = `suc` - `suc_err`,
                    ymax = `suc` + `suc_err`),
                width = .005, position = position_dodge(.9))+
  geom_errorbarh(aes(xmin = `tre` - `tre_err`,
                     xmax = `tre` + `tre_err`),
                 height = .005, position = position_dodge(.9))+
  labs(
    x = '%Synergy with Trehalose',
    y = '%Synergy with Sucrose',
    title = 'Desiccated'
  )

Hydrated | Dry
```
