## Supplementary material for "Disordered proteins interact with the chemical environment to tune their protective function during drying": File S3: CAHS D Oligomerization SAXS Viz.R

```
library (tidyverse)
library (patchwork)
library(readxl)

prot_list <- c("BSA", "AavLEA", "AtLEA3-3", "AvLEA", "HeLEA", "AtLEA4-2", "CAHSD")
Full_list <- c()
Guinier_list <- c()
Kratky_list <- c()
ymax_list <- c(5, 2, 2, 2, 5, 5, 5)
rmarg_list <- c(0.25, 0.01, 0.25, 0.01, 0.25, 0.01, 0.25, 0.01, 0.25, 0.01)

for(i in 1:1)
{
  num = 7
  protein = prot_list[num]
  setwd(paste("C:/Users/Vincent/OneDrive/Documents/csv SAXS Dataset/", protein, sep = ""))
  
  if(protein == "CAHSD")
  {
    file_list <- c("Tris - 4.csv", "1T - 4.csv", "10T - 4.csv", "100T - 4.csv", "1S - 4.csv", "10S - 4.csv", "100S - 4.csv")
    fit_list <- c("Tris Fit - 4.csv", "1T Fit - 4.csv", "10T Fit - 4.csv", "100T Fit - 4.csv", "1S Fit - 4.csv", "10S Fit - 4.csv", "100S Fit - 4.csv")
    col_list <- c("#cccccc", "#8cb8de", "#6393bf", "#36628B", "#9cdbc7", "#63B89C", "#448B74")
  }else
  {
    file_list <- c("Tris - 4.csv", "10T - 4.csv", "100T - 4.csv", "10S - 4.csv", "100S - 4.csv")
    fit_list <- c("Tris Fit - 4.csv", "10T Fit - 4.csv", "100T Fit - 4.csv", "10S Fit - 4.csv", "100S Fit - 4.csv")
    col_list <- c("#cccccc", "#6393bf", "#36628B", "#63B89C", "#448B74")
  }
  
  vals <- read_excel("C:/Users/Vincent/OneDrive/Documents/Boothby Lab Work/Team CAHSD/Oligomerization Project/Oligomerization SAXS Detailed Guinier.xlsx")
  vals <- subset(vals, Protein == protein)
  
  Full <- ggplot() +
    geom_line(data = read_csv(file_list[1]), aes(x = (`Q` * `Q`), y = log(`I(Q)`)), color = col_list[1], pch = 16, size = 1.0) +
    geom_line(data = read_csv(file_list[2]), aes(x = (`Q` * `Q`), y = log(`I(Q)`)), color = col_list[2], pch = 16, size = 1.0) +
    geom_line(data = read_csv(file_list[3]), aes(x = (`Q` * `Q`), y = log(`I(Q)`)), color = col_list[3], pch = 16, size = 1.0) +
    geom_line(data = read_csv(file_list[4]), aes(x = (`Q` * `Q`), y = log(`I(Q)`)), color = col_list[4], pch = 16, size = 1.0) +
    geom_line(data = read_csv(file_list[5]), aes(x = (`Q` * `Q`), y = log(`I(Q)`)), color = col_list[5], pch = 16, size = 1.0) +
    geom_line(data = read_csv(file_list[6]), aes(x = (`Q` * `Q`), y = log(`I(Q)`)), color = col_list[6], pch = 16, size = 1.0) +
    geom_line(data = read_csv(file_list[7]), aes(x = (`Q` * `Q`), y = log(`I(Q)`)), color = col_list[7], pch = 16, size = 1.0) +
    theme_bw()+
    theme(
      plot.title = element_text(hjust = 0.1, size = 10, vjust = -11),
      axis.title = element_text(size = 8),
      axis.text.y = element_text(size = 8),
      axis.text.x = element_text(size = 8),
      legend.text=element_text(size=14),
      legend.position = 'none', aspect.ratio = 1,
      legend.title = element_blank(),
      plot.margin = unit(c(0.001, 1, 0.001, 0.001), "inches"),
      panel.border = element_rect(colour = "black", fill=NA, size=1)) +
    labs(x = expression('q'^2 * ' ' * (Å ^ -1)), y = 'ln(I(q))', title = paste(protein, sep = "")) +
    ylim(-5, 8)
  
  Guinier <- ggplot() +
    geom_point(data = read_csv(file_list[1]), aes(x = (`Q` * `Q`), y = log(`I(Q)`)), color = col_list[1], pch = 16, size = 1.5) +
    geom_line(data = read_csv(fit_list[1]), aes(x = `q**2_fit`, y = `ln(I(q))_fit`), color = "black", size = 1.5) +
    geom_point(data = read_csv(file_list[2]), aes(x = (`Q` * `Q`), y = log(`I(Q)`)), color = col_list[2], pch = 16, size = 1.5) +
    geom_line(data = read_csv(fit_list[2]), aes(x = `q**2_fit`, y = `ln(I(q))_fit`), color = "black", size = 1.5) +
    geom_point(data = read_csv(file_list[3]), aes(x = (`Q` * `Q`), y = log(`I(Q)`)), color = col_list[3], pch = 16, size = 1.5) +
    geom_line(data = read_csv(fit_list[3]), aes(x = `q**2_fit`, y = `ln(I(q))_fit`), color = "black", size = 1.5) +
    geom_point(data = read_csv(file_list[4]), aes(x = (`Q` * `Q`), y = log(`I(Q)`)), color = col_list[4], pch = 16, size = 1.5) +
    geom_line(data = read_csv(fit_list[4]), aes(x = `q**2_fit`, y = `ln(I(q))_fit`), color = "black", size = 1.5) +
    geom_point(data = read_csv(file_list[5]), aes(x = (`Q` * `Q`), y = log(`I(Q)`)), color = col_list[5], pch = 16, size = 1.5) +
    geom_line(data = read_csv(fit_list[5]), aes(x = `q**2_fit`, y = `ln(I(q))_fit`), color = "black", size = 1.5) +
    geom_point(data = read_csv(file_list[6]), aes(x = (`Q` * `Q`), y = log(`I(Q)`)), color = col_list[6], pch = 16, size = 1.5) +
    geom_line(data = read_csv(fit_list[6]), aes(x = `q**2_fit`, y = `ln(I(q))_fit`), color = "black", size = 1.5) +
    geom_point(data = read_csv(file_list[7]), aes(x = (`Q` * `Q`), y = log(`I(Q)`)), color = col_list[7], pch = 16, size = 1.5) +
    geom_line(data = read_csv(fit_list[7]), aes(x = `q**2_fit`, y = `ln(I(q))_fit`), color = "black", size = 1.5) +
    theme_bw()+
    theme(
      plot.title = element_text(hjust = 0.05, size = 10, vjust = -8),
      axis.title = element_blank(),
      axis.text.y = element_text(size = 12),
      axis.text.x = element_text(size = 12),
      legend.text = element_blank(),
      legend.position = 'none', aspect.ratio = 1,
      legend.title = element_blank(),
      plot.margin = unit(c(0.01, 0.01, 0.01, 0.01), "inches"),
      panel.border = element_rect(colour = "black", fill=NA, size=1)) +
    xlim(0, 0.0015) +
    ylim(0, ymax_list[i])
  
  Kratky <- ggplot() +
    geom_point(data = read_csv(file_list[1]), aes(x = `Q`, y = `I(Q)` * `Q` * `Q`), color = col_list[1], pch = 16, size = 1.5) +
    geom_point(data = read_csv(file_list[2]), aes(x = `Q`, y = `I(Q)` * `Q` * `Q`), color = col_list[2], pch = 16, size = 1.5) +
    geom_point(data = read_csv(file_list[3]), aes(x = `Q`, y = `I(Q)` * `Q` * `Q`), color = col_list[3], pch = 16, size = 1.5) +
    geom_point(data = read_csv(file_list[4]), aes(x = `Q`, y = `I(Q)` * `Q` * `Q`), color = col_list[4], pch = 16, size = 1.5) +
    geom_point(data = read_csv(file_list[5]), aes(x = `Q`, y = `I(Q)` * `Q` * `Q`), color = col_list[5], pch = 16, size = 1.5) +
    geom_point(data = read_csv(file_list[6]), aes(x = `Q`, y = `I(Q)` * `Q` * `Q`), color = col_list[6], pch = 16, size = 1.5) +
    geom_point(data = read_csv(file_list[7]), aes(x = `Q`, y = `I(Q)` * `Q` * `Q`), color = col_list[7], pch = 16, size = 1.5) +
    theme_bw()+
    theme(
      plot.title = element_text(hjust = 0.05, size = 10, vjust = -8),
      axis.title = element_text(size = 8),
      axis.text.y = element_text(size = 8),
      axis.text.x = element_text(size = 8),
      legend.text=element_text(size=14),
      legend.position = 'none', aspect.ratio = 1,
      legend.title = element_blank(),
      plot.margin = unit(c(0.001, rmarg_list[i], 0.001, 0.01), "inches"),
      panel.border = element_rect(colour = "black", fill=NA, size=1)) +
    labs(x = expression('q' * ' ' * (Å ^ -1)), y = expression('I(q)' * '*' *'q'^2), title = "")
  
  Full_list[[i]] <- Full 
  Guinier_list[[i]] <- Guinier
  Kratky_list[[i]] <- Kratky
}

Guinier_list[[1]]

Full_list[[1]]

(Full_list[[1]] | Kratky_list[[1]] | Full_list[[2]] | Kratky_list[[2]]) / (Full_list[[3]] | Kratky_list[[3]] | Full_list[[4]] | Kratky_list[[4]]) / (Full_list[[5]] | Kratky_list[[5]] | Full_list[[6]] | Kratky_list[[6]]) / (Full_list[[7]] | Kratky_list[[7]])

(Full_list[[1]] | Full_list[[2]]) / (Full_list[[3]] | Full_list[[4]]) / (Full_list[[5]] | Full_list[[6]]) / (Full_list[[7]])
```
