## Supplementary material for "Disordered proteins interact with the chemical environment to tune their protective function during drying": File S3: Figure_S7.R

```
setwd('~/Documents/Research_Projects/LEA_CAHS_Oligomerization/Submission')

#call in packages
library(tidyverse)
library(dplyr)
library(purrr)
library(broom)
library(stringr)
library(ggplot2)
library(plotly)
library(readxl)
library(RColorBrewer)
library(patchwork)
library(grid)
library(ggsignif)
#DSC 12mg Trehalose-----
T12 <- read_xlsx('Figure_S6.xlsx', sheet = 'Trehalose')
colnames(T12)<- c('Temperature', 'Heat_Flow', 'Sample')
DSC_min <- T12 %>% 
  group_by(Sample)%>%
  summarise_at(vars(Heat_Flow),
               list(min = min))
#create df with discrete heat_flow values
Sample <- c('CT0', 'CT1', 'CT10', 'CT100', 'CT500')
adj_heatflow <- c(-.324, -.33, -.338, -.342, -.348)
adjusted_heatflow <- data.frame(Sample, adj_heatflow)
#combine dataframe
Adjusted <- inner_join(DSC_min, adjusted_heatflow)
#find difference
Adjusted <- Adjusted %>% mutate(Diff= adj_heatflow - min)
#select and merge with DSC 
Adjusted <- Adjusted %>% select(c(Sample, Diff))
#merge
Adjusted_DSC <- inner_join(T12, Adjusted)
Adjusted_DSC <- Adjusted_DSC %>% mutate(adjusted_heatflow = Heat_Flow + Diff)
#import data for enthalpy selection
Trehalose_markers <- read_xlsx('Figure_S6.xlsx', sheet = 'Trehalose_points')
DSC_T12<-ggplot(data = Adjusted_DSC, aes(x = Temperature, y = adjusted_heatflow))+
  geom_line(aes(color = Sample))+
  geom_point(data = Trehalose_markers, aes(x = Temperature, y = adjusted_heatflow), shape = 3, size = 1)+
  ylim(-.35,-.31)+
  theme_bw()+
  theme(axis.text = element_text(size = 6),
        plot.title = element_text(hjust = 0.95, size = 6, vjust = -10),
        axis.title.y = element_text(size = 6),
        axis.title.x = element_text(size = 6),
        legend.position = 'none', aspect.ratio = 1,
        panel.border = element_rect(colour = "black", fill=NA, size=.5),
        panel.grid = element_blank(),,
        axis.text.y = element_blank(),
        axis.ticks.y = element_blank())+
  labs(color = '',
       y = 'Heat flow (W/g)',
       title = '12 mg/mL',
       x = 'Temperature (\u00B0C)')+
  xlim(20,60)+
  
  scale_color_manual(values = c('black', '#B0E0E6','#8CB8DE','#6393BF','#36628B'))
DSC_T12
#12mg Sucrose
S12 <- read_xlsx('Figure_S6.xlsx', sheet = 'Sucrose')
colnames(S12)<- c('Temperature', 'Heat_Flow', 'Sample')
DSC_min <- S12 %>% 
  group_by(Sample)%>%
  summarise_at(vars(Heat_Flow),
               list(min = min))
#create df with discrete heat_flow values
Sample <- c('CS0', 'CS1', 'CS10', 'CS100', 'CS500')
adj_heatflow <- c(-.324, -.33, -.336, -.342, -.348)
adjusted_heatflow <- data.frame(Sample, adj_heatflow)
#combine dataframe
Adjusted <- inner_join(DSC_min, adjusted_heatflow)
#find difference
Adjusted <- Adjusted %>% mutate(Diff= adj_heatflow - min)
#select and merge with DSC 
Adjusted <- Adjusted %>% select(c(Sample, Diff))
#merge
Adjusted_DSC <- inner_join(S12, Adjusted)
Adjusted_DSC <- Adjusted_DSC %>% mutate(adjusted_heatflow = Heat_Flow + Diff)
#import data for enthalpy selection
Sucrose_markers <- read_xlsx('Figure_S6.xlsx', sheet = 'Sucrose_points')
DSC_S12<-ggplot(data = Adjusted_DSC, aes(x = Temperature, y = adjusted_heatflow))+
  geom_line(aes(color = Sample))+
  geom_point(data = Sucrose_markers, aes(x = Temperature, y = adjusted_heatflow), shape = 3, size = 1)+
  
  ylim(-.35,-.31)+
  theme_bw()+
  theme(axis.text = element_text(size = 6),
        plot.title = element_text(hjust = 0.95, size = 6, vjust = -10),
        axis.title.y = element_text(size = 6),
        axis.title.x = element_text(size = 6),
        legend.position = 'none', aspect.ratio = 1,
        panel.border = element_rect(colour = "black", fill=NA, size=.5),
        panel.grid = element_blank(),
        axis.text.y = element_blank(),
        axis.ticks.y = element_blank())+
  labs(color = '',
       y = 'Heat flow (W/g)',
       title = '12 mg/mL',
       x = 'Temperature (\u00B0C)')+
  xlim(20,55)+
  
  scale_color_manual(values = c('black', '#CAE7D3','#9CDBC7','#63B89C','#448B74'))

df <- read_xlsx('Figure_S6.xlsx', sheet = 'Trehalose_12_Enthalpy')
#find mean of different trials
df <- df %>% filter(Sample %in% c('CT1', 'CT10', 'CT100', 'CT500'))
FigS6D <-ggplot(data = df, aes(x = Sample, y = Avg_C_Enthalpy*1000, fill = Sample))+
  geom_bar(position = position_dodge(), stat = 'identity', color = 'black', size = .3)+
  theme_bw()+
  theme(axis.text = element_text(size = 6),
        plot.title = element_text(hjust = 0.95, size = 6, vjust = -10),
        axis.title.y = element_text(size = 6),
        axis.title.x = element_text(size = 6),
        legend.position = 'none', aspect.ratio = 1,
        panel.border = element_rect(colour = "black", fill=NA, size=.5))+
  labs(title = '12 mg/mL',
       x = 'Molar Ratio (Trehalose:CAHS D)',
       y = expression(Delta*'Enthalpy (mJ/g)'))+
  scale_fill_manual(values = c('#B0E0E6','#8CB8DE','#6393BF','#36628B'))+
  scale_x_discrete(breaks=c("CTO","CT1","CT10", "CT100", "CT500"),
                   labels=c("CAHS D","1:1","10:1", "100:1", "500:1"))+
  geom_errorbar(aes(ymin = Avg_C_Enthalpy*1000 - Stdev_C_Enthalpy*1000, ymax = Avg_C_Enthalpy*1000 + Stdev_C_Enthalpy*1000), width = .2, position = position_dodge(.9), size = .3)+
  ylim(-5,90)
FigS6D


df <- read_xlsx('Figure_S6.xlsx', sheet = 'Sucrose_12_Enthalpy')
#find mean of different trials
df <- df %>% filter(Sample %in% c('CS1', 'CS10', 'CS100', 'CS500'))
FigS6E <-ggplot(data = df, aes(x = Sample, y = Avg_C_Enthalpy*1000, fill = Sample))+
  geom_bar(position = position_dodge(), stat = 'identity', color = 'black', size = .3)+
  theme_bw()+
  theme(axis.text = element_text(size = 6),
        plot.title = element_text(hjust = 0.95, size = 6, vjust = -10),
        axis.title.y = element_text(size = 6),
        axis.title.x = element_text(size = 6),
        legend.position = 'none', aspect.ratio = 1,
        panel.border = element_rect(colour = "black", fill=NA, size=.5))+
  labs(title = '12 mg/mL',
       x = 'Molar Ratio (Sucrose:CAHS D)',
       y = expression(Delta*'Enthalpy (mJ/g)'))+
  scale_fill_manual(values = c('#CAE7D3','#9CDBC7','#63B89C','#448B74'))+
  scale_x_discrete(breaks=c("CS0","CS1","CS10", "CS100", "CS500"),
                   labels=c("CAHS D","1:1","10:1", "100:1", "500:1"))+
  geom_errorbar(aes(ymin = Avg_C_Enthalpy*1000 - Stdev_C_Enthalpy*1000, ymax = Avg_C_Enthalpy*1000 + Stdev_C_Enthalpy*1000), width = .2, position = position_dodge(.9), size = .3)+
  ylim(-5,90)

##Tm 6mg Trehalose----
my.cols <- brewer.pal(5, "Blues")
my.cols[1] <- "#000000"
#read in data
Trehalose_6 <-read_xlsx('Figure_S6.xlsx', sheet = 'Trehalose_6_Enthalpy')
CT6<-ggplot(data = Trehalose_6, aes(x = Sample, y = Avg_Tm, fill = Sample))+
  geom_col( 
    position = position_dodge(width = .9), 
    color = 'black', size = .2)+
  #Set graph labels
  labs(x = 'Molar Ratio (Trehalose:CAHS D)',
       y = 'Peak Temperature (\u00B0C)',
       title = '6 mg/mL')+
  ylim(0,45)+
  #Make Black and white theme
  theme_bw()+
  #Adjust graph settings
  theme(axis.text = element_text(size = 6),
        plot.title = element_text(hjust = 0.95, size = 6, vjust = -10),
        axis.title.y = element_text(size = 6),
        axis.title.x = element_text(size = 6),
        legend.position = 'none', aspect.ratio = 1,
        panel.border = element_rect(colour = "black", fill=NA, size=.5))+
  coord_cartesian(ylim=c(30,45))+
  #add error bars based on standard deviation
  geom_errorbar(aes(ymin = Avg_Tm - Stdev_Tm, ymax = Avg_Tm + Stdev_Tm), width = .3, position = position_dodge(.9))+
  scale_x_discrete(labels = c('CT0' = 'CAHS D',
                              'CT1' = "1:1",
                              'CT10' = '10:1',
                              'CT100' = '100:1',
                              'CT500' = '500:1'))+
  scale_fill_manual(values = c('black', '#B0E0E6','#8CB8DE','#6393BF','#36628B'))
##Tm 6mg Sucrose----
my.cols <- brewer.pal(5, "Greens")
my.cols[1] <- "#000000"
#read in data
Sucrose_6 <-read_xlsx('Figure_S6.xlsx', sheet = 'Sucrose_6_Enthalpy')
CS6<-ggplot(data = Sucrose_6, aes(x = Sample, y = Avg_Tm, fill = Sample))+
  geom_col( 
    position = position_dodge(width = .9), 
    color = 'black', size = .2)+
  #Set graph labels
  labs(x = 'Molar Ratio (Sucrose:CAHS D)',
       y = 'Peak Temperature (\u00B0C)',
       title = '6 mg/mL')+
  ylim(0,45)+
  #Make Black and white theme
  theme_bw()+
  #Adjust graph settings
  theme(axis.text = element_text(size = 6),
        plot.title = element_text(hjust = 0.95, size = 6, vjust = -10),
        axis.title.y = element_text(size = 6),
        axis.title.x = element_text(size = 6),
        legend.position = 'none', aspect.ratio = 1,
        panel.border = element_rect(colour = "black", fill=NA, size=.5))+
  coord_cartesian(ylim=c(30,45))+
  
  #add error bars based on standard deviation
  geom_errorbar(aes(ymin = Avg_Tm - Stdev_Tm, ymax = Avg_Tm + Stdev_Tm), width = .3, position = position_dodge(.9))+
  scale_x_discrete(labels = c('CS0' = 'CAHS D',
                              'CS1' = "1:1",
                              'CS10' = '10:1',
                              'CS100' = '100:1',
                              'CS500' = '500:1'))+
  scale_fill_manual(values = c('black', '#CAE7D3','#9CDBC7','#63B89C','#448B74'))
#Tm Trehalose 12 mg
Trehalose_12 <-read_xlsx('Figure_S6.xlsx', sheet = 'Trehalose_12_Enthalpy')
CT12<-ggplot(data = Trehalose_12, aes(x = Sample, y = Avg_Tm, fill = Sample))+
  geom_col( 
    position = position_dodge(width = .9), 
    color = 'black', size = .2)+
  #Set graph labels
  labs(x = 'Molar Ratio (Trehalose:CAHS D)',
       y = 'Peak Temperature (\u00B0C)',
       title = '12 mg/mL')+
  ylim(0,45)+
  #Make Black and white theme
  theme_bw()+
  #Adjust graph settings
  theme(axis.text = element_text(size = 6),
        plot.title = element_text(hjust = 0.95, size = 6, vjust = -10),
        axis.title.y = element_text(size = 6),
        axis.title.x = element_text(size = 6),
        legend.position = 'none', aspect.ratio = 1,
        panel.border = element_rect(colour = "black", fill=NA, size=.5))+
  coord_cartesian(ylim=c(30,45))+
  #add error bars based on standard deviation
  geom_errorbar(aes(ymin = Avg_Tm - Stdev_Tm, ymax = Avg_Tm + Stdev_Tm), width = .3, position = position_dodge(.9))+
  scale_x_discrete(labels = c('CT0' = 'CAHS D',
                              'CT1' = "1:1",
                              'CT10' = '10:1',
                              'CT100' = '100:1',
                              'CT500' = '500:1'))+
  scale_fill_manual(values = c('black', '#B0E0E6','#8CB8DE','#6393BF','#36628B'))
#12 mg/mL Sucrose 
#read in data
Sucrose_12 <-read_xlsx('Figure_S6.xlsx', sheet = 'Sucrose_12_Enthalpy')
CS12<-ggplot(data = Sucrose_12, aes(x = Sample, y = Avg_Tm, fill = Sample))+
  geom_col( 
    position = position_dodge(width = .9), 
    color = 'black', size = .2)+
  #Set graph labels
  labs(x = 'Molar Ratio (Sucrose:CAHS D)',
       y = 'Peak Temperature (\u00B0C)',
       title = '12 mg/mL')+
  ylim(0,45)+
  #Make Black and white theme
  theme_bw()+
  #Adjust graph settings
  theme(axis.text = element_text(size = 6),
        plot.title = element_text(hjust = 0.95, size = 6, vjust = -10),
        axis.title.y = element_text(size = 6),
        axis.title.x = element_text(size = 6),
        legend.position = 'none', aspect.ratio = 1,
        panel.border = element_rect(colour = "black", fill=NA, size=.5))+
  coord_cartesian(ylim=c(30,45))+
  
  #add error bars based on standard deviation
  geom_errorbar(aes(ymin = Avg_Tm - Stdev_Tm, ymax = Avg_Tm + Stdev_Tm), width = .3, position = position_dodge(.9))+
  scale_x_discrete(labels = c('CS0' = 'CAHS D',
                              'CS1' = "1:1",
                              'CS10' = '10:1',
                              'CS100' = '100:1',
                              'CS500' = '500:1'))+
  scale_fill_manual(values = c('black', '#CAE7D3','#9CDBC7','#63B89C','#448B74'))


DSC_T12 +DSC_S12+FigS6D+FigS6E+ CT6 + CS6 + CT12+ CS12+ plot_layout(ncol = 4)+
plot_annotation(tag_levels = list(c('B)','C)','D)','E)','F)','G)','H)','I)'))) & theme(plot.tag = element_text(size = 8))
```
