## Supplementary material for "Disordered proteins interact with the chemical environment to tune their protective function during drying": File S3: Betaine Script.R

```
library (tidyverse)
library (patchwork)
library(readxl)

prot_list <- c("CAHSD + Betaine")
Full_list <- c()
Guinier_list <- c()
Kratky_list <- c()
ymax_list <- c(5)
rmarg_list <- c(0.1)

for(i in 1:length(prot_list))
{
  num = i
  protein = prot_list[i]
  setwd(paste("C:/Users/Vincent/OneDrive/Documents/csv SAXS Dataset/", protein, sep = ""))
  
  Dim_Kratky <- read_csv("Dimensionless_Kratky_(Rg).csv")
  
  vals <- read_excel("C:/Users/Vincent/OneDrive/Documents/Boothby Lab Work/Team CAHSD/Oligomerization Project/Oligomerization SAXS Detailed Guinier.xlsx")
  vals <- subset(vals, Protein == protein)
  
  file_list <- c("Tris - 4.csv", "10B - 4.csv", "100B - 4.csv", "500B - 4.csv")
  fit_list <- c("Tris Fit - 4.csv", "10B Fit - 4.csv", "100B Fit - 4.csv", "500B Fit - 4.csv")
  col_list <- c("#cccccc", "#A63A2E66", "#A63A2EAA", "#A63A2EFF")
  
  Full <- ggplot() +
    geom_line(data = read_csv(file_list[1]), aes(x = (`Q` * `Q`), y = log(`I(Q)`)), color = col_list[1], pch = 16, size = 1.0) +
    geom_line(data = read_csv(file_list[2]), aes(x = (`Q` * `Q`), y = log(`I(Q)`)), color = col_list[2], pch = 16, size = 1.0) +
    geom_line(data = read_csv(file_list[3]), aes(x = (`Q` * `Q`), y = log(`I(Q)`)), color = col_list[3], pch = 16, size = 1.0) +
    geom_line(data = read_csv(file_list[4]), aes(x = (`Q` * `Q`), y = log(`I(Q)`)), color = col_list[4], pch = 16, size = 1.0) +
    theme_bw()+
    theme(
      plot.title = element_text(hjust = 0.1, size = 10, vjust = -11),
      axis.title = element_text(size = 8),
      axis.text.y = element_text(size = 8),
      axis.text.x = element_text(size = 8),
      legend.text=element_text(size=14),
      legend.position = 'none', aspect.ratio = 1,
      legend.title = element_blank(),
      plot.margin = unit(c(0.001, 1, 0.001, 0.001), "inches"),
      panel.border = element_rect(colour = "black", fill=NA, size=1)) +
    labs(x = expression('q'^2 * ' ' * (Å ^ -1)), y = 'ln(I(q))', title = paste(protein, sep = "")) +
    ylim(-5, 8)
  
  Kratky <- ggplot() +
    geom_line(data = Dim_Kratky, aes(x = Dim_Kratky$`Tris - 4.dat_qRg`, y = Dim_Kratky$`Tris - 4.dat_(qRg)^2*I(q)/I(0)`), color = col_list[1], size = 1.0) +
    geom_line(data = Dim_Kratky, aes(x = Dim_Kratky$`10B - 4.dat_qRg`, y = Dim_Kratky$`10B - 4.dat_(qRg)^2*I(q)/I(0)`), color = col_list[2], size = 1.0) +
    geom_line(data = Dim_Kratky, aes(x = Dim_Kratky$`100B - 4.dat_qRg`, y = Dim_Kratky$`100B - 4.dat_(qRg)^2*I(q)/I(0)`), color = col_list[3], size = 1.0) +
    geom_line(data = Dim_Kratky, aes(x = Dim_Kratky$`500B - 4.dat_qRg`, y = Dim_Kratky$`500B - 4.dat_(qRg)^2*I(q)/I(0)`), color = col_list[4], size = 1.0) +
    theme_bw()+
    theme(
      plot.title = element_text(hjust = 0.05, size = 10, vjust = -8),
      axis.title = element_text(size = 8),
      axis.text.y = element_text(size = 8),
      axis.text.x = element_text(size = 8),
      legend.text=element_text(size=14),
      legend.position = 'none', aspect.ratio = 1,
      legend.title = element_blank(),
      plot.margin = unit(c(0.001, 1, 0.001, 0.001), "inches"),
      panel.border = element_rect(colour = "black", fill=NA, size=1)) +
    labs(x = expression('qRg'), y = expression('(I(q)/I(0))' * '*' *'qRg'^2), title = "") +
    ylim(-1, 8)
  
  Full_list[[i]] <- Full 
  Guinier_list[[i]] <- Guinier
  Kratky_list[[i]] <- Kratky
}

Guinier_list[[1]]

Kratky_list[[1]]

Full_list[[1]]
```
