## Supplementary material for "Disordered proteins interact with the chemical environment to tune their protective function during drying": File S3: Fig_S8E_Betaine_control.R

```
setwd("~/Documents")

#call packages
library(dplyr)
library(tidyverse)
library(ggplot2)
library(readxl)
library(ggsci)
library(reshape2)
library(drc)
library(scales)
library(patchwork)
library(wesanderson)

data <-read_xlsx('Betaine_Control_Review.xlsx')
Betaine_Control <-ggplot(data = data, aes(x = Seconds, y = Abs, color = Sample))+
  geom_point(size = .5)+
  theme_bw()+
  theme(
    plot.title = element_text(hjust = 0.5, size = 6),
    axis.title = element_text(size =6),
    axis.text.y = element_text(size =6),
    axis.text.x = element_text(size =6),
    legend.text=element_text(size=14),
    legend.position = 'none', aspect.ratio = 1,
    legend.title = element_blank(),
    plot.background = element_blank(),
    panel.grid.major = element_blank(),
    panel.grid.minor = element_blank(),
    panel.border = element_blank())+
  theme(axis.line = element_line(color = 'black'))+
  labs(x = 'Time (seconds)', y = 'Absorbance (340 nm)')+
  scale_color_manual(values = c('#8B0000','grey'))
Betaine_Control + plot_layout(ncol = 4)
```
