## Supplementary material for "Disordered proteins interact with the chemical environment to tune their protective function during drying": File S4

**LEA motif sequences**

**>At11**

AKSKADETLES

**>Aav11**

LRDTAAEKLHQ

**>He11**

LKDKAGSAWNQ

**>Av11**

KDKAAEALDAI

**>At22**

AKSKADETLESAKSKADETLES

**>At44**

AKSKADETLESAKSKADETLESAKSKADETLESAKSKADETLES

**>At20**

AKEKLNIGGAKAQGHAEKTM

**Full-length proteins**

**>AtLEA3-3**

MASHQEQSYKAGETRGKAQEKTGEAMGTMGDKTQAAKDKTQETAQSAQQKAHETAQSAKDKTSQAAQTTQERAQESKDKTGSYMSETGEAIKNKAHDAAEYTKETAEAGKEKTSGILGQTGEQVKQMAMGATDAVKHTLGLRTDEGNKEHVSSAPSTTTTTTTRETQRK

**>AavLEA1**

MSSQQNQNRQGEQQEQGYMEAAKEKVVNAWESTKETLSSTAQAAAEKTAEFRDSAGETIRDLTGQAQEKGQEFKERAGEKAEETKQRAGEKMDETKQRAGEMRENAGQKMEEYKQQGKGKAEELRDTAAEKLHQAGEKVKGRD

**>HeLEA68614**

MFLARNVSRVALRSVSLSPAAIPQQQHAGVAAVYAVRFASSSGSGRPADNWAESQKEKAKAGLKDAQAEVGKVAREVKDKAAGGIEQAKDAVKQGANDLKRSGSRTFENAKDDIQAKAQHAKSDLKGAKHQAEGVVENVKEAAENAWEKTKDVAENLKDKVQSPGGLADKAANAWETVKDRAQDAASEVKHKAGDLKDKAQQVIHDATTQSGDNRKQDQQQRRDSQGSQSGQNSRSRN

**>AvLEA1C**

MNKFLSILCLVLCISATFAKQSATEQAVNAAADLKDKVKDAASAAYDAASPKVAEGAEFIKDKAEQAYETGSKVAGEYADVAKEKLAKVADDVKASAQNFANDASKTGQEYAQEGLKQGQKLGEQAFEVGKDKANEALKAAQKSGADAYEAALEYGADGVKRAKQLPEQTVELSRDKANEALKAARHAAGDSIDSATEYVQETRKQASKKAKETTEEASEKAQKAKRNADL

**>AtLEA4-2**

MQSAKEKISDMASTAKEKLNIGGAKAQGHAEKTMARTKKEKKLAQEREKSKEAQAKADLHQSKAEHAADAQVHGHHLPGHSTYPTRATGANYPPGQI

**>CAHS D**

MSGRNVESHMERNEKVVVNNSGHADVKKQQQQVEHTEFTHTEVKAPLIHPAPPIISTGAAGLAEEIVGQGFTASAARISGGTAEVHLQPSAAMTEEARRDQERYRQEQESIAKQQEREMEKKTEAYRKTAEAEAEKIRKELEKQHARDVEFRKDLIESTIDRQKREVDLEAKMAKRELDREGQLAKEALERSRLATNVEVNFDSAAGHTVSGGTTVSTSDKMEIKRN
